# Spatiospectral brain networks reflective of improvisational experience

**DOI:** 10.1101/2021.02.25.432633

**Authors:** Josef Faller, Andrew Goldman, Yida Lin, James R. McIntosh, Paul Sajda

## Abstract

Musical improvisers are trained to categorize certain musical structures into functional classes, which is thought to facilitate improvisation. Using a novel auditory oddball paradigm (Goldman et al., 2020) which enables us to disassociate a deviant (i.e. musical cord inversion) from a consistent functional class, we recorded scalp EEG from a group of musicians who spanned a range of improvisational and classically trained experience. Using a spatiospectral based inter and intra network connectivity analysis, we found that improvisers showed a variety of differences in connectivity within and between large-scale cortical networks compared to classically trained musicians, as a function of deviant type. Inter-network connectivity in the alpha band, for a time window leading up to the behavioural response, was strongly linked to improvisation experience, with the default mode network acting as a hub. Spatiospectral networks post response were substantially different between improvisers and classically trained musicians, with greater inter-network connectivity (specific to the alpha and beta bands) seen in improvisers whereas those with more classical training had largely reduced inter-network activity (mostly in the gamma band). More generally, we interpret our findings in the context of network-level correlates of expectation violation as a function of subject expertise, and we discuss how these may generalize to other and more ecologically valid scenarios.

## 1. Introduction

Improvisation has received scholarly attention in recent years from a variety of disciplinary perspectives. While often associated with musical performance, improvisation is theorized to underlie a wide variety of human behaviors ranging from artistic practices to organizational management to the performance of gender (Lewis & Piekut, 2016). Following from definitions of creativity in the psychology literature, improvisation can be characterized as the spontaneous formation of novel, high quality output, that is novel and useful (Sternberg et al., 2004). Recent work has begun to coalesce knowledge and models from electroencephalography (EEG) studies (Stevens Jr & Zabelina, 2019), the involvement of the motor system (Bashwiner & Bacon, 2019), the importance of expertise (Pinho et al., 2014; Braun, 2008), perception-action coupling (Loui, 2018), top-down and bottom-up networks (Faber & McIntosh, 2020), and network neuroscience (Beaty et al., 2019; Belden et al., 2020).

Western musical improvisation offers an important model for the more general study of improvisation. Western musical improvisers can create and play music spontaneously, guided only (if at all) by notation that does not specify exact notes, but instead specifies functional classes of harmonies and melodies with multiple possible realizations, or instantiations as notes (e.g., jazz lead sheets, or figured bass notations).

Improvisers are free to play any notes that fit these functional classes, subject to certain constraints, such as musical syntax, aesthetic considerations, and style or appropriateness for the audience (Berliner, 1994). Intriguingly, Western classically trained musicians, following a musical aesthetics that reifies specific series of notes as musical works (Goehr, 1992), are trained to perform these works strictly following the musical score and rarely ever improvise harmonic or melodic aspects of the music; to change those aspects would be to change the work of music, contradicting the aesthetics of the classical music tradition. Presumably as a result of the specific nature of this training, a classically trained musician who may have trained playing an instrument just as many years as an improviser - just in a different way - may not be able to improvise music.

Previous work found that jazz improvisers showed more pronounced, larger early right anterior negativity (ERAN) to rare and unexpected targets (Przysinda et al., 2017). Magnitudes of these ERAN responses correlated with metrics for improvisation experience and P3b and ERAN correlated with fluency and originality in divergent thinking tasks. Aligned with these findings Zabelina & Ganis (2018) reported that individuals with greater ability in divergent thinking showed shorter response times and a stronger N2 ERP deflection for rare target trials which the authors interpret as higher attentional flexibility and stronger engagement of cognitive control processes in divergent thinkers. Musicians with higher improvisation experience were further found to show lower BOLD activation in the right motor area (inferior frontal gyrus or IFG, anterior insula), regions associated with the default mode network or DMN (angular gyrus), the dorsolateral prefrontal cortex or DLPFC (Pinho et al., 2014) and higher upper-alpha power frontally during improvisation relative to control conditions (Lopata et al., 2017). These findings are supported by studies which contrasted brain activity during musical improvisation relative to control tasks within individuals in fMRI (Limb & Braun, 2008; Bengtsson et al., 2007; de Manzano & Ullén, 2012; Liu et al., 2012; Kouneiher et al., 2009), and complemented by electro- and magnetoencephalography-based studies which, in slightly different tasks, reported increased theta, alpha and beta power (Sasaki et al., 2019), decreased theta, alpha and beta power (Adhikari et al., 2016), or increased alpha and theta, but decreased beta power (Boasen et al., 2018).

When studying improvisation experience in terms of differences in brain connectivity, Pinho et al. (2014) reported that individuals with more improvisation experience showed greater connectivity between DLPFC and motor regions (dorsal premotor cortex or dPMC, pre-supplementary motor area or pre-SMA) based on BOLD-based functional connectivity. Work by the same authors (Pinho et al., 2015) supported the original findings when brain connectivity was studied within-subject during improvisatory activity relative to control conditions. Work by other authors in fMRI (Dhakal et al., 2019) and EEG (Adhikari et al., 2016) on the other hand reported on evidence for decreased granger causality-based connectivity.

Very recent work has focused on studying connectivity between large-scale cortical networks with Belden et al. (2020) showing that musical improvisation experience can be predicted from resting state fMRI in that improvisers showed higher connectivity between primary visual network and DMN/ECN (executive control network) as well as higher connectivity between DMN and ECN while classically trained musicians on the other hand showed higher connectivity between vDMN and frontal pole. Earlier studies on creativity in non-music related contexts support these findings, reporting that creative individuals may be able to simultaneously engage large-scale networks that normally work in opposition, like default mode, salience and executive control networks (Beaty et al., 2018b). Further support comes from studies that showed that the interaction between large-scale networks predicted openness (Beaty et al., 2018a), was associated with high figural creativity (Liu et al., 2018) and may underlie the inhibition of prepotent responses (Beaty et al., 2017).

Goldman et al. (2020) theorized that the specific way western musical improvisers are trained to categorize notes into higher level structures like functional-harmonic classes of chords may facilitate their ability to improvise. In music theory, harmonies can be classified by their function; roughly, in a series of harmonies, various chords play the role of “tonic” harmonies, some can function as “pre-dominant harmonies” and some as “dominant harmonies,” depending on their placement within syntactically ordered series of harmonies. Different chords can play these different functional roles: for example, in some musical contexts, an improviser can substitute a chord with the notes G-B-D for one with the notes Db-F-Ab; these two chords share no notes, but can serve the same dominant function. Being able to substitute one harmony for another within the same functional class constitutes an important part of widely practiced forms of improvisation, and would underlie other important skills like recognizing a bandmate’s substitutions in order to more fluently respond and interact with them. Thus, in the study, the authors hypothesized that trained improvisers may perceive different chords within a functional class as more similar than chords that belong to different functional classes, whereas musicians without improvisatory training would not show the influence of such categorizations on their harmonic perception.

The authors tested this hypothesis in an EEG study using an auditory oddball paradigm where improvisers and classically trained musicians listened to progressions of three chords where the middle chord was either a deviant in terms of its musical inversion, but still picked from within the same functional class, referred to as “exemplar deviant” (7.5% probability), a deviant that also lay outside the functional class, referred to as “function deviant” (7.5% probability), or a standard (no change in inversion; same functional class; 85% probability). In support of their hypotheses, Goldman et al. found that musicians with more improvisation experience were slower and less accurate at detecting exemplar deviants relative to function deviants, i.e., deviant harmonic stimuli outside of the functional class were more salient than deviants within the functional class. In addition, more experienced improvisers also showed less pronounced N2c and P3b event-related potential (ERP) responses to exemplar deviants relative to function deviants, interpreted as a relatively lower violation of expectancy.

Here we build on the data collected by Goldman et al. (2020) to investigate whether connectivity between cortical networks could help explain how musicians perceive and process musical structures, and whether improvisatory training leads to characteristic differences in such processing. We use connectivity and band power to isolate and measure spatiospectral brain networks and processes related to how musicians perceive chords within and across functional-harmonic categorical boundaries. We focus on whether the amount of improvisatory training can predict differences between these measurements. Again, as described by Goldman et al. (2020), this difference helps explain an important aspect of improvisatory training, perception, and performance. We focus on canonical cortical networks (Williams, 2016), some of which have been implicated in improvisation by previous studies (Belden et al., 2020), specifically networks related to attention (including frontoparietal network and dorsal attention network; e.g. Marek & Dosenbach (2018), Fornito et al. (2012) and Vossel et al. (2014)), cognitive control (e.g. Niendam et al. (2012)), salience (also including cingulo opercular network; e.g. Seeley (2019), Seeley et al. (2007) and Dosenbach et al. (2006)) and the default mode network (e.g. Fornito et al. (2012)). In an analysis inspired by Hanada et al. (2019) we derived connectivity within and between these networks as follows: We first recovered neuroelectrical source activity for every constituent region of given networks (e.g. ACC, DLPFC, etc.) using inverse methods (cortically constrained low resolution tomography; Pascual-Marqui et al. (2002)). We then computed directed connectivity between regions within and between networks using a validated signal processing pipeline (Mahjoory et al., 2017) that made use of a connectivity metric (phase slope index, PSI, Nolte et al. (2008)) that was theoretically and empirically shown (Nolte et al., 2008) to be robust to volume conduction effects as they appear in EEG (Haufe et al., 2013). These network metrics were then separately computed for exemplar and function deviants and the difference between these scalar values was used to linearly predict self-reported weekly improvisation hours, weekly hours spent training classical music and a behavioral metric (Goldman et al., 2020; Townsend & Ashby, 1978) that reflected the difference in task performance between exemplar and function deviants. We analyzed the resulting spatiospectral networks for three time windows: 1) between presentation of the second and third chord (*between chords*), 2) prior to the response (*pre-response*) and 3) after the response (*post-response*) (see Fig. 1).

**Figure 1:**
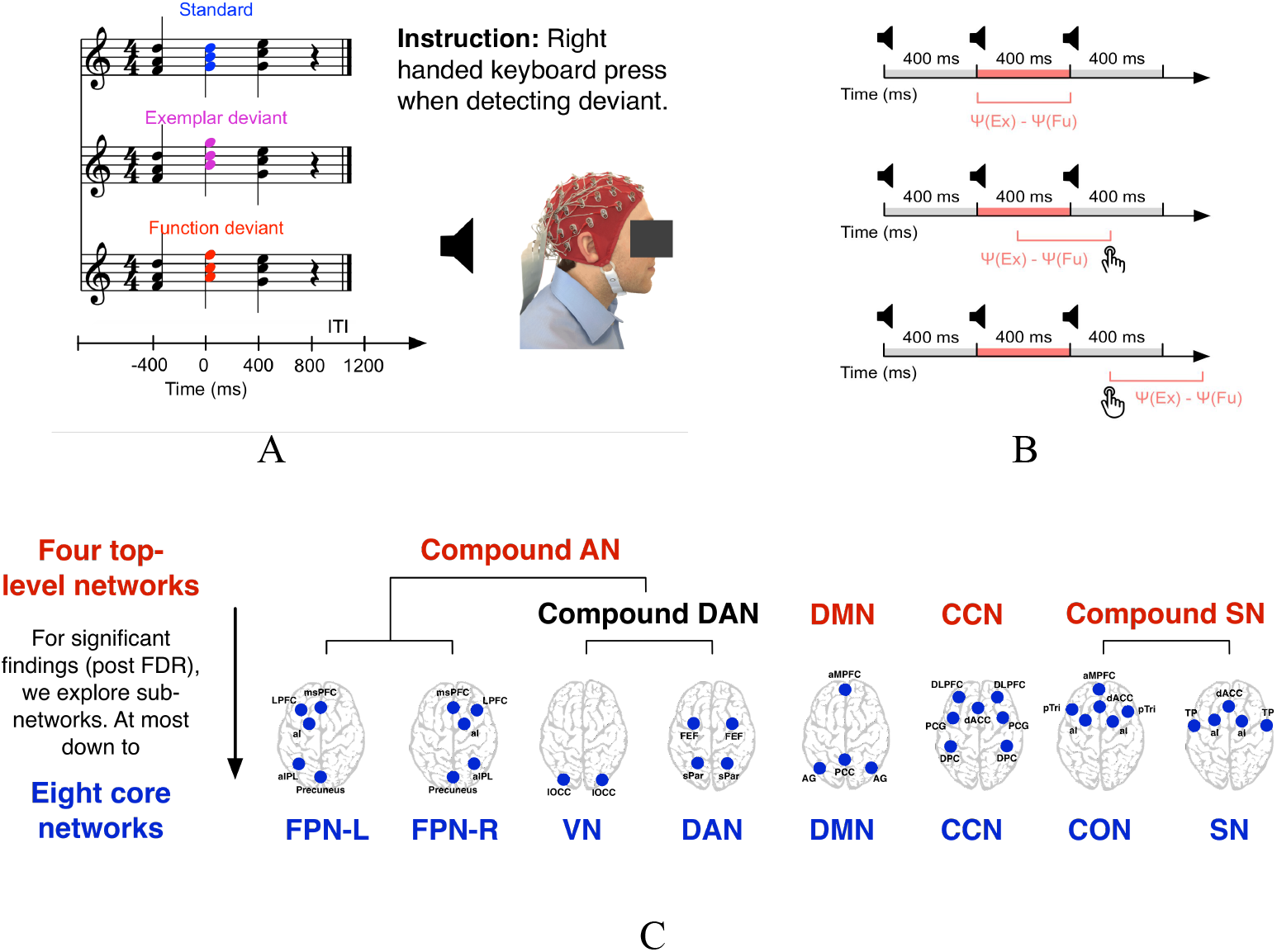
Experimental Paradigm. (A) Subjects (all musicians) where instructed to listen to chord progressions, each consisting of three chords, and respond with a button press if they heard a deviant. There were two types of deviants, one being “exemplar” and one “functional” (see main text for details). Each chord progression was considered a trial and EEG was recorded during the entire experiment. (B) Analysis of the data, with respect to differences in network connectivity between exemplar (Ψ(Ex)) and functional (Ψ(Fu)) deviants, was focused on three time windows, the 400 ms between the second and third chord (*between chords*), the 400ms before the behavioural response (*pre-response*) and finally the 400ms after the behavioural response (*post-response*). (C) The canonical brain networks investigated, both in terms of inter and inter-network connectivity, using phase-slope index measures (PSI). Networks include the left (FPN-L) and right (FPN-R) fronto-parietal network, the visual network (VN), the dorsal attention network (DAN) the default mode network (DMN) the cognitive control network (CCN) the cingulo opercular network (CON) and the salience network (SN). Three compound networks were also considered: the compound DAN, the compound SN and the compound attention network (AN). Networks were fully connected.

## 2. Materials and Methods

### 2.1. Study participants

The data for this analysis has been collected by Goldman et al. (2020): A total of 40 musicians with formal training and/or significant professional experience (mean age 25.3, s.d. 5.5; 24 male) completed the experiment, with 25 of the subjects reporting ≥ 1 hour/week improvisation training on average since age 18. The musicians’ primary instruments were piano (*N_p_*=14), wind (*N_w_*=15) and string instruments (*N_s_*=11). Eight musicians reported being able to perfectly assess pitch of musical notes in absence of a reference tone (“absolute pitch”, Ward (1999)). All participants reported normal hearing and no history of neurological disorders. The study was approved by the institutional review board of Columbia University (NY, USA) and all subjects provided written informed consent prior to participation in the experiments.

### 2.2. Auditory oddball task

The musicians were instructed to listen to chord progressions, that each consisted of three chords. We refer to one instance of such a progression in the recording as a trial. Every one of the three chords in one trial sounded in sequence, each for 400ms in piano timbre, after which each trial ended with another 400 ms silence. This resulted in a fixed, total trial length of 1600 ms. The only progressions used in the experiment were ii-IV-I, ii-V-I, ii-IV6-I and ii-V6-I (this notation reflects chord configurations as shown in Figure 1A). Each experimental block consisted of 180 trials. For each such block one of the four aforementioned progressions were chosen as “standard”, resulting in four types of blocks (see Goldman et al. (2020) for details). These “block types” were used to counterbalance the effect of other features of the individual progressions such as intervallic content that may have been in themselves salient (refer to Goldman et al. (2020) for further explanation). An experimental block always started with at least eight “standard” trials for the purpose of allowing participants to learn what type of progression would be the standard for the current block. There were two types of deviant trials that each occurred at a probability of 7.5% (in total 15%). Every deviant trial was followed by at least three standard trials. Deviant trials only differed from standard trials in terms of the middle chord: (1) Exemplar deviants, where the middle chord was replaced with a chord of identical notes but different inversion. For example, if the middle chord for a standard trial in that experimental block was V then the middle chord for the exemplar deviant in that block would be V6. For (2) function deviants, the middle chord was replaced by a chord from a different functional class. For example, if the middle chord for a standard was again V, then the middle chord for the corresponding function deviant in that block would be IV (again, see Figure 1A). Importantly, the key for each trial’s chord progression was picked at random. This meant that musicians needed to examine the second chord of every trial relative to the first and/or third to identify whether the trial was a standard or deviant. The order of standards and deviants within every one of the four types of experimental blocks was generated once only, and was thus identical across subjects within these block types. For the experiment, every one of the block types occurred twice, thus resulting in a total of eight blocks per subject. The order of the eight blocks was shuffled for every subject. In total, there were 1440 trials per subject of which 222 were functional and 218 were exemplar deviants. See Goldman et al. (2020) for further details.

### 2.3. Data collection

While the musicians performed the oddball task, their EEG was recorded from 64 gel-based, active electrodes at standard scalp locations (10/20 system; Oostenveld & Praamstra (2001)) at a sampling rate of 2048Hz using a biosignal amplifier (Biosemi ActiveTwo, Biosemi, The Netherlands). The subjects were seated comfortably at a desk inside a shielded room as the auditory oddball paradigm was played to them via noise-cancelling, in-ear headphones (Quiet Comfort 20, Bose Corp., MA, USA). Subjects were instructed to respond to deviant chords as quickly and accurately as possible, by pressing the space-bar on a computer keyboard on the desk in front of them using the index finger of their right hand. This auditory stream was also recorded as a separate channel via the biosignal amplifier to assure highly accurate synchronization of paradigm timing, EEG and behavioral responses.

### 2.4. Preprocessing

Figure 2 shows an overview of the signal processing pipeline, where every participant’s EEG was first filtered bi-directionally with the pass-band configured from 0.5 to 45 Hz (finite-impulse response filter; order 6144, tripling the raw sampling rate). The filtered signal was then down-sampled from 2048 to 256 Hz.

**Figure 2:**
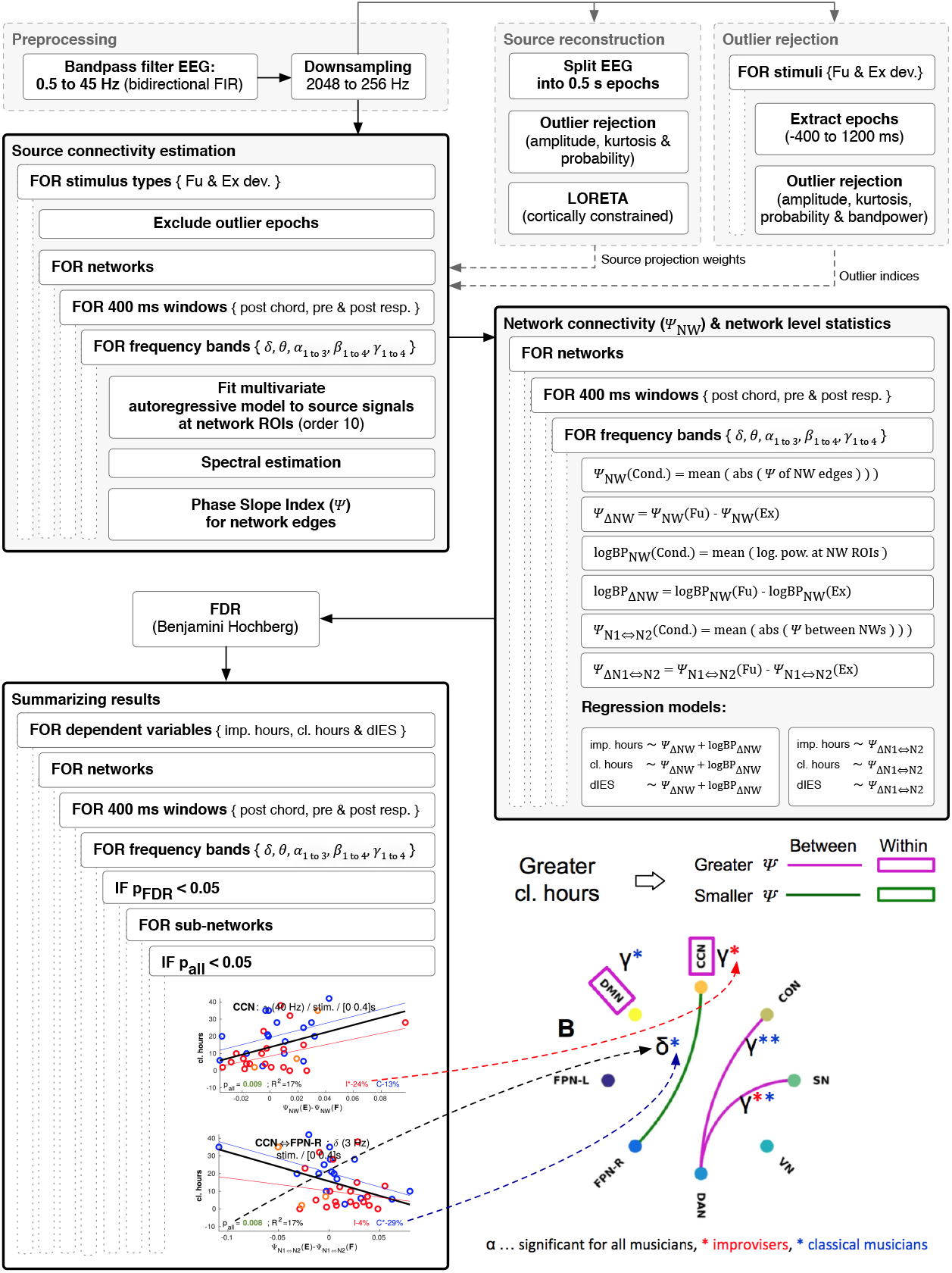
Flowchart summarizing data processing and analysis used in the study. Each block includes a summary of steps for the data processing and analysis that was done: EEG Preprocessing, Source reconstruction, Outlier rejection, Source connectivity estimation, Network connectivity and network level statistics and methodology for Summarizing results. The lower right figure shows how the results are presented in terms of intra and inter-network interactions. This example network analysis is for the dependent variable cl. hours, so the number of reported weekly hours spent training classical performance. Results of the PSI analysis are shown with boxes (for intra-network connectivity) and edges (inter-network connectivity) with color indicating the direction of the effect. Pink indicates that musicians with greater reported weekly hours spent training classical performance (cl. hours) also showed greater connectivity for exemplar relative to function deviants. Green on the other hand indicates lower connectivity for exemplar relative to function deviants. Each connectivity measure for a network (either box/intra or edge/inter) is associated with one or more spectral bands, indicating the frequencies at which the connectivity is significant. Black greek letters indicate significant effects (i.e. *p* < 0.05) across all musicians. Colored asterisks indicate which connectivity (box/intra or edge/inter) is additionally significant for improvisers only (red *) or classical musicians only (blue *). One, two and three * correspond to treshold levels for p-values of 0.05, 0.01 and 0.001. Further details are provided in the main text.

### 2.5. Reconstruction of electrical activity at specific brain regions

Neuroelectrical signals at specific cortical regions of interest (ROIs) in the brain, from hereon referred to as cortical current source density (CSD) signals, were inferred from the observed EEG by applying the inverse method anatomically constrained low resolution brain electromagnetic tomography (cLORETA, Pascual-Marqui et al. (2002)) to a boundary element method (BEM) based “forward model” of how current propagates from a cortical neuronal source through neural tissue, cerebrospinal fluid, skull and out to the scalp. The first step in the procedure was automatic epoch-based outlier rejection based on the Matlab toolbox EEGLAB (Delorme & Makeig, 2004), where the subject’s EEG was split into epochs of 0.5s and epochs were rejected when their signal exceeded commonly used thresholds for amplitude (smaller or greater 200 *μV*), kurtosis (> 5.5 × *SD* for the subject) or probability (> 4.0 × *SD* for the subject). The procedure for estimating CSD was identical to García-Cordero et al. (2017), where the BEM solution was computed using OpenMEEG (Gramfort et al., 2010; Kybic et al., 2005) using the MRI based brain anatomy model “Colin 27” (Holmes et al., 1998) that was non-linearly mapped into MNI305 space (Evans et al., 1993) and associated with standard EEG electrode locations using BrainStorm (Tadel et al., 2011). Inverse modelling was accomplished through cLORETA, by which the 64 scalp EEG channels were first linearly mapped to a 5003-vertex cortical mesh and from there to 202 regions according to a sub parcellated version of the Desikan-Killiany atlas (Desikan et al., 2006).

### 2.6. Trial based outlier rejection

After outlier rejection was first performed prior to source reconstruction, the obtained source space projection matrix was then applied to raw EEG signal. Prior to actual analysis of experimental trials, outlier epochs were identified separately for the three conditions of standards, function and exemplar deviants. For each condition, epochs were extracted from −400 to 1200 ms relative to the onset of the second chord in a progression and epochs were rejected according to the previously mentioned criteria for amplitude, kurtosis, probability and additionally as per a custom iterative band power based method (Faller et al., 2012). For the iterative method log-transformed band power was computed for frequency bands in delta, theta, alpha, beta and gamma up to 50 Hz. Trials were marked as outliers if average log-transformed power for the trials in any of the bands fell outside the mean ± 4 standard deviations of how all trials in that band and subject were distributed. If more than 0 outlier trials were marked, then the procedure was repeated based on a mean and standard deviation that did not take the outlier trials into account.

### 2.7. Connectivity estimation between brain regions

Conceptually, our analysis starts with four top-level brain networks (related to attention, cognitive control, default mode and salience; see Figure 1C). Some of these top-level networks (e.g. the network we refer to as the “compound” Attention Network), are composed of sub-networks, and ultimately of eight “core” networks (see Figure 1C). When statistically significant effects (post FDR) are observed in top-level networks, we continue analysis in sub-networks in an effort to localize effects. Specifically in terms of computation, the first step in our approach is to calculate the directed connectivity metric PSI separately for every subject, every trial type (standards and both deviants), for every brain network (starting with the four top-level networks), for twelve EEG frequency bands, three time windows (0 to 400ms, relative to the second chord, as well as −400 to 0 and 0 to 400 ms relative to the response) and for every edge within the fully connected networks. CSD time series for the nodes in every network were obtained by averaging across signals that corresponded to subparcellations as per the mapping from reconstructed source signals using the Desikan-Killiany atlas (Desikan et al., 2006) as described above. A separate multivariate autoregressive model (order 10) was then fit to these CSD time series separately for every network, every time window and trial type using the Levinson-Wiggens-Robinson algorithm (Morf et al., 1978) as implemented in the Biosig toolbox (Vidaurre et al., 2011) used by Fieldtrip (Oostenveld et al., 2011). Through Fourier transform, we obtained cross spectral densities for the pairs of source time series for which we wanted to study connectivity relationships (i.e. edges in the network graphs; see Fig. 1C). The phase of these cross spectral densities was then analyzed to derive PSI (denoted as Ψ)for the corresponding network edges according to Nolte et al. (2008) using default parameters in Fieldtrip for EEG frequency bands ± 2 Hz relative to the center frequencies shown in Table 1.

**Table 1:** Center frequencies for each band used in the PSI analysis

| | $\delta$ | $\theta$ | $\alpha_1$ | $\alpha_2$ | $\alpha_3$ | $\beta_1$ | $\beta_2$ | $\beta_3$ | $\beta_4$ | $\gamma_1$ | $\gamma_2$ | $\gamma_3$ |
| --- | --- | --- | --- | --- | --- | --- | --- | --- | --- | --- | --- | --- |
| Center frequency (Hz) | 3 | 6 | 8 | 10 | 12 | 16 | 20 | 24 | 28 | 32 | 36 | 40 |

PSI makes use of the fact that if a signal in a frequency band that spans the adjacent frequencies *f*_1_ to *f_n_* in *x_a_*(*t*) is reproduced with a time delay *τ* later in another signal *x_b_*(*t*), then the phase spectrum of complex coherency is linear over this contiguous range of frequencies *f*1 to *f_n_* with a positive slope proportional to the time delay *τ*. If signal *x_b_*(*t*) instead would lead signal *x_a_*(*t*) in time, then a negative slope would be observed. A more formal definition for PSI as per Nolte et al. (2008) is

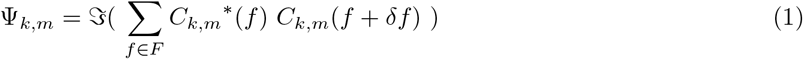

where *k* and *m* indicate the indices of the signals between which to calculate connectivity, 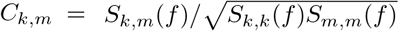 represents complex coherency, *S* the cross-spectral density matrix, *f* is one out of a set *F* of frequencies in a small band for which to calculate PSI, *f* the frequency resolution, the asterisk denotes taking the conjugate transpose and 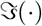 denotes taking the imaginary part of a complex number.

### 2.8. Estimation of connectivity within brain networks

To capture connectivity regardless of directionality across edges over a whole cortical network in a robust manner we defined a simple metric Ψ*NW*, for which the absolute value was taken for the PSI value for every edge of a network before all these absolute values were simply averaged. More formally, and based on definitions by Nolte et al. (2008) this can be represented as

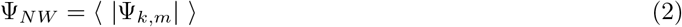

where Ψ, indexed by *k* and *m* represents the PSI between the brain signals *k* and *m* that correspond to pairs of nodes within the network, |·| denotes taking the absolute value and < · > denotes expected value.

### 2.9. Estimation of connectivity between brain networks

Connectivity between networks was assessed by first computing PSI between the nodes of different networks. For example, connectivity was computed between one ROI in network 1 and every ROI in network 2 and so forth. Then we again took the absolute value for all these PSI results, and finally averaged across all the results. That way we obtained one scalar value reflective of overall connectivity between one pair of networks.

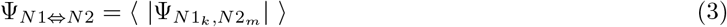

### 2.10. Estimation of band power within brain networks

Average activity across a network as expressed in signal amplitude was captured by computing logarithm transformed bandpower for every region of interest (node) in the network and then averaging across the results for these nodes. More formally,

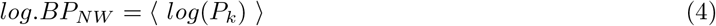

where *P* are the band power values, averaged across trials, for brain signals *k* that correspond to network constituent nodes and < · > again denotes the expected value.

### 2.11. Statistical prediction of experience and behavior from network connectivity

Robust regression (Holland & Welsch, 1977) was used to separately predict improvisation experience and behavioral performance in the oddball task from two independent variables that were based on overall connectivity NW in large-scale canonical cortical networks for function and exemplar deviants. Improvisation experience (imp. hours) was represented by average weekly hours of practice in musical improvisation since age 18 as reported by the musicians in a questionnaire prior to the experiment, and non-improvisatory experience (cl. hours) was represented by average weekly hours of non-improvisatory (e.g., classical-style) practice (Goldman et al., 2020). As per the hypotheses of Goldman and colleagues, improvisers should react more slowly and less accurately to detecting exemplar relative to function deviants, since improvisers regularly train to substitute chords with other chords from the same functional class and standards and exemplar deviants were from within the same functional class. This was captured in the following behavioral metric

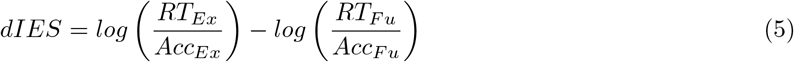

where *RT* and *Acc* represent average response time and accuracy for the respective deviant conditions of exemplar and function deviants. *A* positive value of *dIES* corresponds to function deviants being easier to detect, while a negative value corresponds to exemplar deviants being easier to detect. The following regression models were thus evaluated across networks (starting with the four top-level networks; see Fig 1C), three time windows and twelve frequency bands:

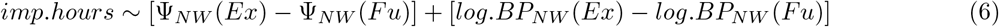

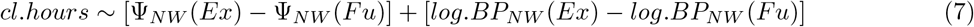

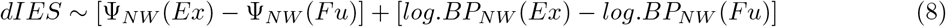

where expressions in [·] represent one variable and the abbreviations *Fu*, *Ex* and *Sta* represent the three stimulus conditions.

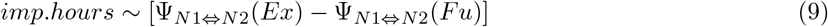

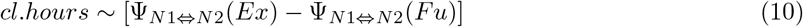

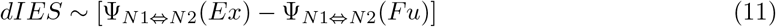

After false-discovery rate (FDR; Benjamini & Hochberg (1995)) based correction on model level (number of comparisons: 3 dependent variables x number of networks x 3 time windows x 12 frequencies), models that resulted in FDR-corrected p-values < 0.05 were further studied using robust regression directly on the independent variables; on that level p-values < 0.05 were considered statistically significant. Whenever we were fitting data for improvisers alone, three improvisers were conservatively excluded since we found that they, on occasion, represented overly influential data points (represented as orange instead of red circles in scatter plots in the supplemental material).

## 3. Results

We present results in terms of the time windows of analysis, shown in Figure 1B: *between chords*, *pre-response*, *post-response*. As we are discussing greater or lower connectivity, we are specifically referring to greater connectivity for exemplar relative to function deviants (i.e. *ψ*(*Ex*) – *ψ*(*Fu*)), consistent with Equations (6) to (11).

### 3.1. Stimulus locked analysis between chords

#### 3.1.1. Reduced connectivity between DAN and CON networks for improvisers relative to classically trained musicians

In a time window of 400 ms directly following the onset of the audio of deviant chords, musicians with greater improvisation experience showed lower connectivity between canonical brain networks in the alpha and beta band for the exemplar relative to the function deviant (see Fig. 3A). Opposing effects between musical disciplines were observed for connectivity between cingulo opercular and dorsal attention network, where greater improvisation experience, was associated with lower connectivity in the beta band (*p_FDR_* = 0.036, *R*^2^ = 17.8%; Fig. 3A), while greater experience with classical music, in comparison, was associated with greater connectivity in the gamma band (*p_FDR_* = 0.046, *R*^2^ = 15.5%; Fig. 3B). Further noteworthy effects were found when predicting improvisation experience between cognitive control and right frontoparietal network in the alpha band (*p_FDR_* = 0.036, *R*^2^ = 16.8%) and finally between salience and visual network in the alpha (*p_FDR_* = 0.044, *R*^2^ = 16.3%) and beta band (*p_FDR_* = 0.017, *R*^2^ = 25.5%), all shown in Fig. 3A.

**Figure 3:**
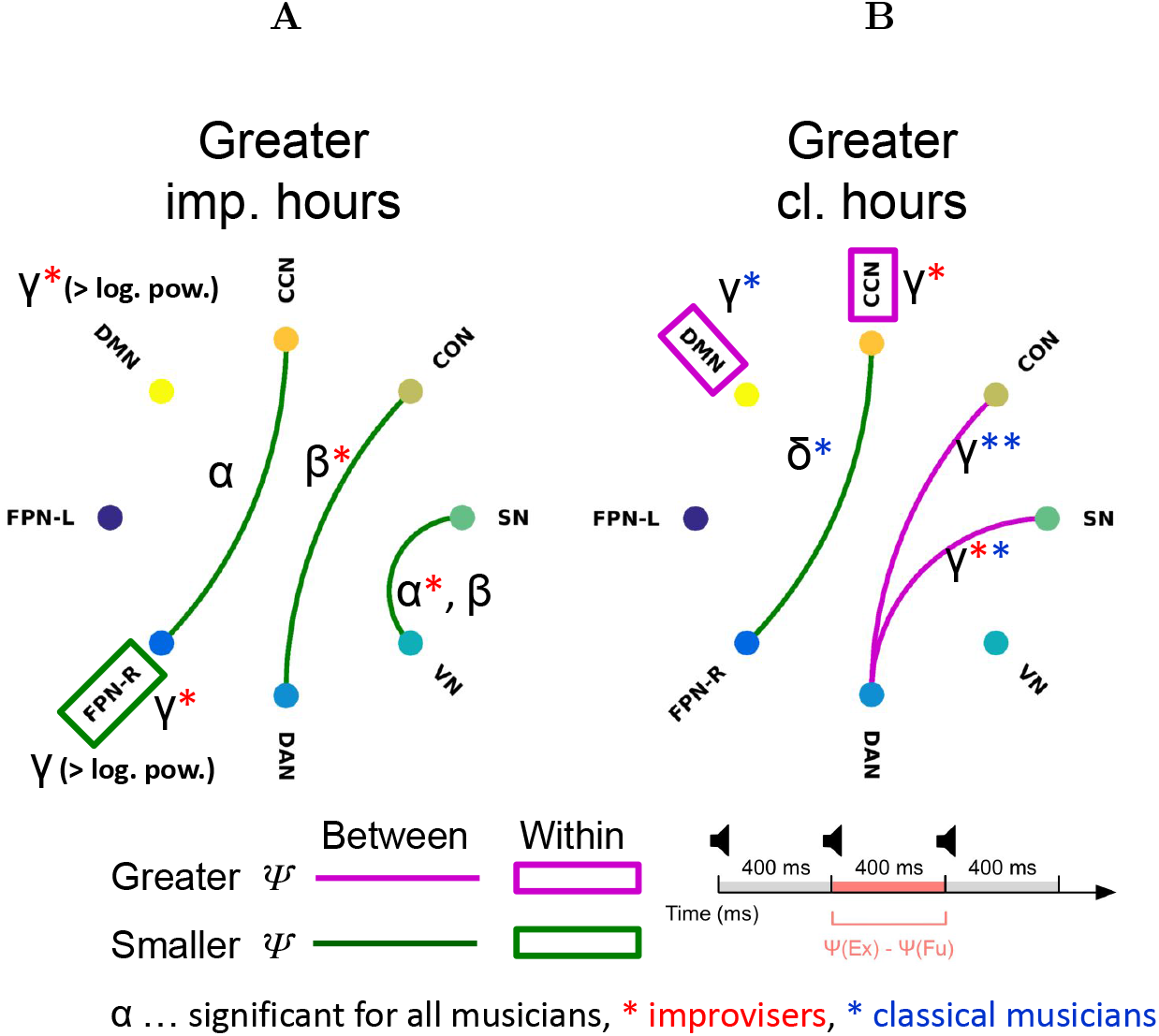
Spatiospectral networks for between chords analysis. Bottom right shows the time window of the analysis (refer back to Fig 1B). (A) Musicians with greater improvisation experience showed lower inter-network connectivity between canonical brain networks in the alpha and beta band for the exemplar relative to the function deviant. Specifically these effects were found between cognitive control (CCN) and right frontoparietal (FRN-R) networks in the alpha band and between the cingulo opercular (CON) and dorsal attention (DAN) networks in the beta band and between salience (SN) and visual (VN) networks in the alpha and beta bands. Intra-network connectivity was lower in the FRN-P. In addition both the FRN-P and default mode network (DMN) showed greater logarithmic gamma power. (B) Greater experience performing classical music was likewise associated with lower inter-network connectivity between CCN and FPN-R, though the effect was in the delta rather than alpha band. Greater inter-network connectivity was seen between DAN and CON and DAN and SN, both in the gamma band. Increased intra-network activity was seen in both the DMN and CCN, once again specifically for the gamma band.

#### 3.1.2. Greater experience irrespective of discipline was associated with reduced connectivity between CCN and FPN-R

In this time window directly following the audio of the chord, we further found effects within the right frontoparietal network (*p_FDR_* = 0.010, *R*^2^ = 40.8%; see Fig. 3A). Specifically, greater improvisation experience was associated with lower connectivity within the network in the gamma band (*p* = 0.013, *R*^2^ = 16.7%) and greater logarithmic power also in the gamma band (*p* = 0.007, *R*^2^ = 19.4%). Within the default mode network, greater improvisation experience was associated with a significant effect (*p_FDR_* = 0.013, *R*^2^ = 40.8 %), specifically greater logarithmic power in the gamma band (*p* = 0.006, *R*^2^ = 24.6%).

Greater experience performing classical music was likewise associated with lower connectivity between cognitive control and right frontoparietal network for exemplar relative to function deviants (*p_FDR_* = 0.049, *R*^2^ = 15.2%; Fig. 3B). However, the effect was found in the delta band whereas for improvisation experience the effect was found in the alpha band. In short, the higher the average weekly hours of experience, irrespective of musical discipline, the lower the connectivity between cognitive control and right frontoparietal network for exemplar relative to function deviants (see Fig. 3A and B).

Greater experience in performing classical music was also associated with greater connectivity within the cognitive control (*p_FDR_* = 0.049, *R*^2^ = 23.5%) and within the default mode network (*p_FDR_* = 0.045, *R*^2^ = 24.8%; Fig.3B).

Furthermore, while musicians with more improvisation experience had exhibited lower connectivity between salience and visual network in the alpha (*p_FDR_* = 0.044, *R*^2^ = 16.3%) and beta band (*p_FDR_* = 0.017, *R*^2^ = 25.5%; Fig. 3A), musicians with greater experience in classical music showed greater connectivity between salience and dorsal attention network in the gamma band (*p_FDR_* = 0.046, *R*^2^ = 15.0%; Fig. 3B).

In this time window directly following the onset of the deviant chords, greater brain connectivity between networks for the exemplar relative to the function deviant tended to be associated with greater dIES, meaning a slower and less accurate response to exemplar relative to function deviants (see Figures S.5 and S.6). We found behavioral effects for most connections where we found effects related to experience with improvisation and classical music, except between the cognitive control and right frontoparietal network. Results were less consistent for within-network effects in this time window. Specificially, it was only for the default mode network that we found a behavioral effect that also matched the finding related to self reported average weekly hours training classical music.

In summary, musicians who reported greater average weekly hours of training for either musical discipline showed lower connectivity between cognitive control and right frontoparietal network in the 400 ms following the onset of an exemplar deviant relative to the same time window for a function deviant. Between cingulo opercular and dorsal attention network, greater improvisation experience was associated with lower connectivity, while greater experience in classical music was associated with higher connectivity. Finally, improvisers exhibited lower connectivity between salience and visual network, while musicians with greater classical experience showed greater connectivity between salience and dorsal attention network.

### 3.2. Pre-response analysis: Improvisers show distinctive inter-network connectivity in the alpha band with robust effects between DMN and VN

In the 400 ms before the motor response to an exemplar deviant chord - a chord that was experimentally manipulated to fall in the same functional class as the standard, but was otherwise like the function deviant chord - musicians with greater improvisation experience showed greater connectivity between brain networks, all relative to when the musicians responded to a function deviant and exclusively in the alpha band (see Fig. 4).

**Figure 4:**
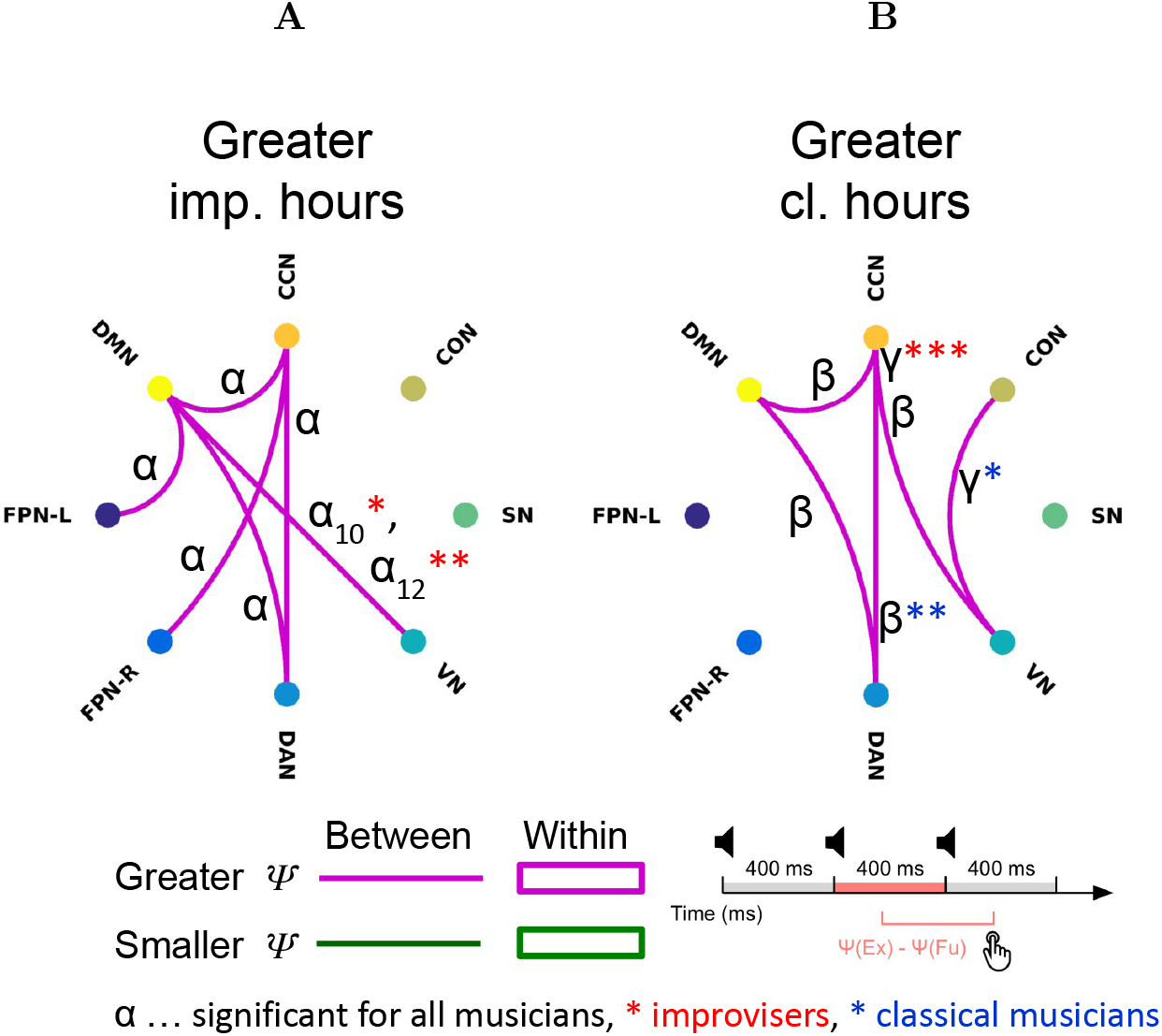
Spatiospectral networks for pre-response analysis. Bottom right shows the time window of the analysis (refer back to Fig 1B). (A) Musicians with greater improvisation experience showed greater inter-network connectivity between a number of canonical brain networks in the alpha band for the exemplar relative to the function deviant. In this case the default mode network (DMN) acted as a “hub”. (B) Greater experience performing classical music was likewise associated with greater inter-network connectivity between a number of the canonical networks, though this effect was found in the beta and gamma bands. There were no significant intra-network connectivity changes seen for either (A) or (B).

The default mode network acted as a hub with greater connectivity to the left frontoparietal, cognitive control, dorsal attention and visual network. The effect between default mode and visual network stood out as it was not only significant for all musicians (10 Hz: *p_FDR_* = 0.042, *R*^2^ = 17.4%; 12 Hz: *p_FDR_* = 0.016, *R*^2^ = 24.8%; Fig4A) but also for the smaller subset of “improvisers” alone (i.e. only musicians with self-reported average weekly hours spent improvising > 0.5), where the effect was most robust for a center frequency of 12hz (*p* = 0.008, *R*^2^ = 42.4%), followed by a center frequency of 10Hz (*p* = 0.050, *R*^2^ = 25.8%). Notably, musicians with more improvisation experience also showed greater connectivity between the cognitive control and the right frontoparietal network (*p* = 0.003, *R*^2^ = 22.0%; Fig.4A).

We also identified a group of three fully interconnected networks (i.e. a “clique” or “rich club” from a graph-theoretical perspective; Griffa & Van den Heuvel (2018)) that was composed of the default mode, cognitive control and dorsal attention network. Interestingly, when studying how between network connectivity related to musicians’ experience with classical music we observed the same sub structure such that musicians with greater self reported average weekly hours of practice in classical music since age 18 showed greater connectivity between default mode, cognitive control and dorsal attention network, so just like for improvisation experience - except in the beta rather than alpha band (see Fig. 4A and B).

Furthermore, in terms of associations with experience in classical music, we found no effect between default mode and visual network, but instead musicians with greater self-reported experience in classical music showed greater connectivity between the cognitive control and visual network in the beta (*p_FDR_* = 0.048, *R*^2^ = 14.6%; Fig. 4B) and particularly the gamma band (*p_FDR_* = 0.021, *R*^2^ = 23.6%; Fig. 4B). Interestingly, the latter effect was particularly robust for the sub group of improvisers alone (*p* = 5.89e^-4^, *R*^2^ = 51.1%).

Statistically significant associations between task performance (*dIES*) and inter-network connectivity were found broadly in the alpha, beta and gamma band as well as less often (< 5 times) in the delta and theta band (see Fig. S.8). In almost all cases the association was such that greater connectivity between networks was associated with greater *dIES*, meaning a slower and less accurate response for exemplar relative to function deviants. Only less than five cases showed an effect in the opposite direction.

Importantly, for all effects found for improvisation experience (i.e. self-reported hours of improvisation experience), except between default mode and dorsal attention network, we found effects for task performance that matched in timing, frequency and direction of the effect (see Fig. S.8). This means that connectivity between these networks was not only directly proportional to self reported improvisation experience, but also directly proportional to slower and less accurate responding to exemplar deviants, thus supporting the hypothesized link between inter-network connectivity, improvisation experience and modified behavior.

In summary, in the 400 ms before responding to a deviant chord that was experimentally manipulated to fall in the same functional class as the standard chord in an oddball task, musicians who reported greater improvisation experience showed greater connectivity between canonical cortical brain networks in the alpha band with the default mode network acting as a hub and particularly robust effects found between default mode and visual network. Greater experience in classical music was likewise associated with greater inter-network connectivity, however consistently in the beta and gamma as opposed to the alpha band. Inter-network connectivity effects between three networks, default mode, cognitive control and dorsal attention network overlapped between musical disciplines. Greater inter-network connectivity that was observed with greater improvisation experience, was consistently also associated with slower and less accurate responding to the manipulated exemplar deviant relative to the function deviant supporting the hypothesized link between improvisation experience and slower and less accurate responding to audio of chords that improvisers are trained to categorize differently (Goldman et al., 2020).

### 3.3. Post-response analysis: For improvisers DAN and VN acted as network hubs whereas for classically trained musicians, CCN acted as a hub

In the 400 ms after responding to an exemplar as compared to a function deviant, improvisers with greater improvisation experience tended to exhibit greater connectivity between networks with the dorsal attention and visual network acting as hubs (Fig. 5A), all mainly in the beta and gamma band, while they showed lower connectivity between the default mode and visual network (*p_FDR_* = 0.034, *R*^2^ = 18.6%).

**Figure 5:**
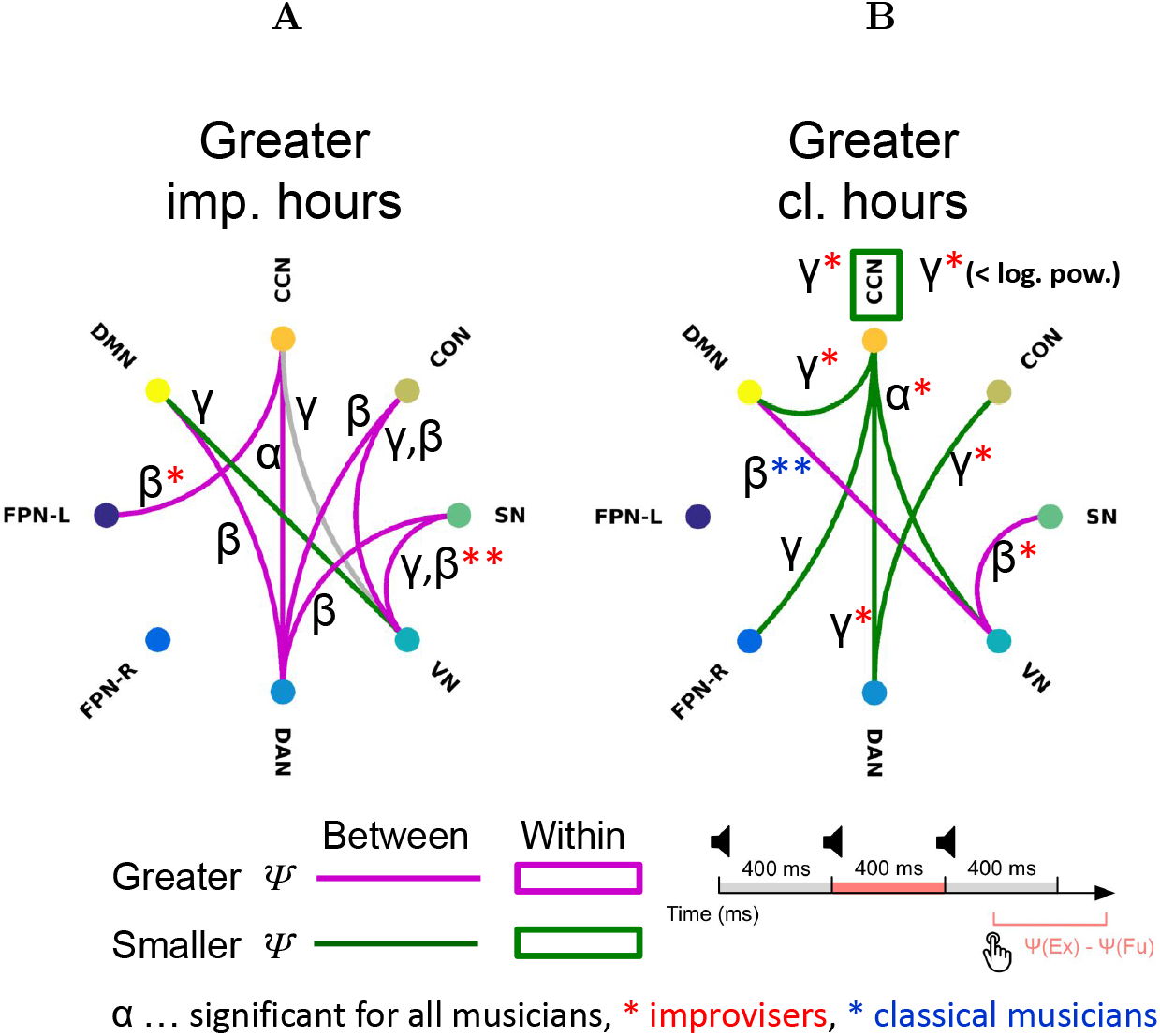
Spatiospectral networks for post-response analysis. Bottom right shows the time window of the analysis (refer back to Fig 1B). (A) Musicians with greater improvisation experience showed greater inter-network connectivity (for the exemplar relative to the function deviant) with the dorsal attention (DAN) and visual (VN) networks acting as hubs. This inter-network connectivity was mainly in the beta and gamma band. The grey link between the cognitive control network (CCN) and VN indicates a rare case where two frequency bands (*γ*1 and *γ*3) within the gamma range show an effect in opposing directions. (B) We observed an opposite effect for musicians with greater experience performing classical music, namely lower inter-network connectivity, with the CCN and VN acting as hubs. Also observed was lower intra-network connectivity in the CCN and reduced low power in the CCN, both in the gamma band.

Meanwhile, the effects observed for musicians with greater experience in classical music tended to point in the opposite direction such that greater experience was associated with lower connectivity between networks where the cognitive control and visual network acted as hubs (Fig. 5B). Musicians with greater experience in classical music also exhibited greater connectivity between default mode and visual network, so again the opposite of what was found for improvisation experience.

While connectivity from the cognitive control network to other networks was lower for greater self reported experience with classical music (Fig. 5B), we also found significant effects within the cognitive control network (*p_FDR_* = 0.035, *R*^2^ = 26.8%), specifically lower within network connectivity (*p* = 0.032, *R*^2^ = 23.2%) and lower logarithmic bandpower (*p* = 0.024, *R*^2^ = 13.0%) in the gamma band.

For this time window directly following the motor response, we further found that greater musical expertise was associated with greater connectivity between the salience and visual network both for improvisation (beta: *p_FDR_* = 0.034, *R*^2^ = 18.4%; gamma: *p_FDR_* = 0.016, *R*^2^ = 26.9%; Fig. 5A) and classical performance (*p_FDR_* = 0.050, *R*^2^ = 14.5%; Fig.5B).

Behavioral effects in this time window between default mode and visual network as well as between the salience and visual network and more broadly were such that greater connectivity for exemplar relative to function deviants was associated with slower and less accurate responses to exemplar relative to function deviants. Hypothetically, slower and less accurate responses to exemplar relative to function deviants are linked to the training improvisers receive, so that we assumed a musician who responds slower and less accurately to an exemplar deviant may have received more training in improvisation. For the connection between default mode and visual network we observe that lower connectivity for exemplar relative to function deviants was associated with greater improvisation experience, which constitutes a disagreement (Fig. 5A). For the connection between salience and visual network as well as more broadly for other effects related to improvisation experience in this time window we tended to find agreement.

In summary, for the 400 ms following motor response to the experimentally manipulated exemplar deviant as compared to a function deviant, we found that improvisers showed lower connectivity between default mode and visual network, greater connectivity between salience and visual network as well as an overall increased connectivity between networks, where the dorsal attention, the visual network and to a lesser degree the cognitive control network acted as hubs. Greater experience in classical performance training was likewise associated with greater connectivity between salience and visual network, but also with greater connectivity between default mode and visual network as well as lower connectivity widely between networks where the cognitive control network acted as a hub. Within the cognitive control network, both connectivity and logarithmic power in the gamma band were lower for musicians with greater experience in training classical performance.

## 4. Discussion

Leveraging the high temporal resolution of EEG (Rosen et al., 2020; Zabelina & Ganis, 2018; Marek & Dosenbach, 2018), and through our focus on network connectivity guided by fMRI findings, (Belden et al., 2020; Beaty et al., 2018b; Pinho et al., 2014), we asked what networked neural processes, if any, may underlie how improvisers perceive and process chords differently, given their training to think about harmony categorically (Goldman et al., 2020). We took into account activity that manifests as average EEG band power across a network as well as connectivity within or between large-scale cortical networks (Cohen & D’Esposito, 2016).

The exemplar deviant chord in the oddball task in this experiment was designed to be part of the same functional class as the frequent and expected standard chord, while the function deviant was equivalent to the exemplar deviant, except that the function deviant belonged to a functional class other than the standard. Improvisers are trained to substitute chords within a functional class, and thus we hypothesized that improvisers would categorize the exemplar deviant as being more similar to the standard, which we assumed should cause improvisers to respond slower and less accurately to exemplar relative to the function deviants. This idea is supported empirically also by findings by Goldman et al. (2020), who reported a statistically significant relationship such that greater *dIES* corresponded to greater self-reported weekly hours of improvisation training since age 18.

In our purely auditory task, musicians responded with their right hand to chords that were deviants in terms of chord inversion, but musicians were successfully kept blind (as verified by post-experiment interviews) to the fact that there were two types of deviants and that one of these types, referred to as exemplar deviant, was modified such that it fell within the same functional class as the standard chord (Goldman et al., 2020). Improvisers are trained to categorize chords within the same functional class separately, as being usable interchangeably in improvisatory performance. We studied neural responses surrounding exemplar deviants but specifically after subtracting the response for function deviants, such that we could expect that any effects we observe should be specifically tied to our experimental manipulation related to categorization of musical structures.

One finding that stood out was that connectivity related effects between networks before improvisers responded to an exemplar relative to a deviant chord were consistently and exclusively found in the alpha band. In contrast, connectivity related effects associated with experience in classical music before the response were only found in the beta and gamma band. It’s noteworthy that significant findings in the alpha band were otherwise rare and most findings were either in the beta or gamma band.

To our knowledge this is the first report indicating that improvisers may exhibit greater between network connectivity specifically in the alpha band even by just responding to a rare chord that was manipulated to fall in the same functional class as the standard chord in an oddball task. In fact we are not aware of any report on connectivity between networks in improvisers in any brain-state occurring primarily in the alpha band. Finding an alpha related effect for improvisers in the connectivity between networks is not implausible though, given that there is ample evidence implicating the alpha oscillation in musical improvisation with reports of both increased (Sasaki et al., 2019; Boasen et al., 2018) or decreased (Adhikari et al., 2016) alpha power while musicians improvise in slightly different experiments. Beyond musical improvisation, amplitude changes in the alpha oscillation have been robustly linked to domain general creativity as measured for example by divergent thinking tasks (Zabelina & Ganis, 2018; Fink et al., 2007; Jauk et al., 2012; Schwab et al., 2014) or compound remote associates tasks (Rothmaler et al., 2017), with a relatively high heterogeneity in the direction of effects (Dietrich & Kanso, 2010; Arden et al., 2010) ascribed to the diversity in tasks and methods (Fink et al., 2014), but with findings overall leaning toward increased frontal and parietal alpha power for greater creativity (Dietrich & Kanso, 2010), where one interpretation pointed toward a hypothetical function of alpha in attenuating top-down control (Lustenberger et al., 2015). Given however, that our results are based on connectivity between brain regions rather than amplitude at certain regions, we think what we observe may be most consistent with changes in network organization and/or function that may be caused by intense training in musical improvisation. Results from graph-analyses based on fMRI (Belden et al., 2020) and EEG (N=4; Wan et al. (2014)) point to greater global network integration for improvisers as opposed to a more densely connected local organization for musicians with greater training in classical music. These findings in turn are consistent with the idea that improvisers may, through training, become very efficient at flexibly engaging and balancing a variety of mental processes with substrates in distributed brain regions (de Manzano & Ullén, 2012) related to executive control and accessing long-term/working memory in realtime (Lopata et al., 2017; Belden et al., 2020) without the necessity of conscious mediation (Limb & Braun, 2008; Liu et al., 2012). Our findings of effects of inter-network connectivity in the alpha band for improvisers in contrast with effects in higher frequency bands for classically trained musicians, support the idea that long range oscillatory communication may be an important factor in creative cognition (Stevens Jr & Zabelina, 2019). According to this idea, also referenced by Boasen et al. (2018), different EEG frequency bands are thought to be linked to different scales of cortical integration (Von Stein & Sarnthein, 2000) such that high frequency oscillations represent local communication while theta and alpha oscillations are linked to long-range/inter-areal integration (Haegens et al., 2010; Klimesch et al., 2007; Clayton et al., 2015). In summary, we interpret the observed effects in the alpha band for improvisers to indicate that even when improvisers merely respond to an “in-class” chord (a chord in the same functional class as the standard) they co-engage cortical resources more broadly than classically trained musicians or musicians with less extensive training in improvisation. This supports the idea that music genre specific training may be accompanied by significant genre-specific changes in neurophysiology (Loui, 2018; Bianco et al., 2017) and the outcome of our experiment indicates that this may extend to how improvisers categorize musical structures.

Also leading up to the right handed response, we observed that greater reported weekly hours, irrespective of type of training were associated with greater connectivity between a group of three fully connected networks (a ‘clique” or “rich-club” in terms of graph theory; Griffa & Van den Heuvel (2018)) consisting of default mode, cognitive control and dorsal attention network was found for both disciplines, which we interpret to mean that connectivity between these networks is task related and linked to training in musical performance in general (Loui, 2018; Bianco et al., 2017), irrespective of discipline. We consider the existence of such an effect plausible and potentially scientifically interesting by itself. Given how improvisers and classically trained musicians are different groups with relatively little overlap in this sample of musicians, this finding might also be interpretable as evidence in support of the fidelity of this method.

Given that our focus lies on neurophysiological differences specific to improvisation we direct our attention to effects outside this clique of networks leading up to the manual response. Another effect that stood out in that time window was that improvisers showed greater connectivity between default mode and visual network leading up to the response, but less connectivity between these two networks after the response. Classically trained musicians also showed greater connectivity between default mode and visual network, however only after the response.

Activity in the default mode and other large-scale cortical networks including the dorsal attention network has typically been found to be anti-correlated in fMRI studies (Fornito et al., 2012). Finding increased connectivity between these networks here is consistent with the idea that creativity may depend on the flexible engagement of generative and evaluative processes (Sowden et al., 2015; Zabelina & Robinson, 2010) and aligns with reports in fMRI literature, where positively correlated engagement of large-scale cortical networks was linked to experience in musical improvisation (Belden et al., 2020), greater creativity (Beaty et al., 2018b, 2019) or openness to experience (Beaty et al., 2017).

The default mode network specifically, is traditionally associated with self-referential processing (Kim & Johnson, 2014), but as outlined by Belden et al. (2020) also with musical behaviors like tracking of musical tonality (Janata et al., 2002), associating music with autobiographical memories (Janata, 2009) or aesthetic response to episodic memory retrieval (Schacter & Addis, 2007). Specifically, the DMN’s role in memory retrieval as part of a greater role in creative cognition (Benedek et al., 2014) may be of particular interest for this investigation. Overall, a number of studies have linked default mode network activity (Beaty et al., 2015; Rosen et al., 2017) and interaction between default mode and other networks such as the frontoparietal network (Beaty et al., 2018b, 2019; Belden et al., 2020) to creativity and musical improvisation.

Occipital areas that overlap what we defined here as visual network on the other hand, have been previously implicated in creativity, as reviewed by Belden et al. (2020), where greater white (Takeuchi et al., 2017) or grey (Fink et al., 2014) matter density in the occipital lobe, as well as greater white matter connectivity in the inferior occipitofrontal fasciculus (Zamm et al., 2013) were found to be associated with greater creativity. Belden et al. (2020) also found greater connectivity between the visual network and the default mode as well as a network similar to what we here defined as the frontoparietal network (Belden referred to it as executive control network), in resting-state recordings of musicians with improvisation experience. Belden et al. contrasted their findings to Beaty et al. (2018b) who had found no evidence for involvement of occipital regions in a network linked to creativity in resting-state fMRI.

Given the findings of Belden et al. (2020) we assume that there may exist a baseline effect between default mode and visual network at rest for improvisers. However, since that should be present for function deviants as well, for which we correct by subtracting the signal acquired for function deviants, we assume that the observed effect is in fact tied to our experimental manipulation related to categorization of musical structures. One explanation for the observed effects could be that connectivity between default mode and visual network reflects an access to long-term memory that is engaged only or stronger for “in-class” chords and supports how improvisers categorize musical structures according to functional classes, maybe here concretely by supporting the comparison of categories between the standard chord in working memory and categorization related information about the just perceived exemplar deviant from long-term memory. However, the fact that classically trained musicians or less extensively trained improvisers show greater connectivity between default mode and visual network as well, but post-response, indicates that this categorization related phenomenon may not by itself necessarily exclusively subserve creative demands reserved only for improvisers. Instead, a more plausible explanation could be that strongly trained improvisers adapt, through training, to prioritize this process to a degree where it is executed before the manual response since an improviser’s response in an ecologically valid setting may have to strongly depend on the result of this process. In other words, as per this theory, as a musician improvises they may permanently check that the chord they just heard (or played) is a constituent of the currently appropriate functional class and/or need to make sure the chord they are playing next is likewise part of that or whatever next appropriate functional class. Somebody who is not a strongly trained improviser may not or less strongly engage this process before a response to an “in-class” chord.

For improvisers we further observed greater connectivity between default mode and left frontoparietal network, which aligns with previous accounts that implicated these networks (Bashwiner et al., 2016; Mok, 2014; Shi et al., 2018) and in particular increases in connectivity between them (Kenett et al., 2018) in supporting creative cognition (Belden et al., 2020) and high creative ability (Zabelina & Robinson, 2010). One idea is that these networks may represent cortical hubs that underlie the dual-process model of creative cognition (Sowden et al., 2015; Stanovich, 1999; Evans, 2008, 2009) with the default mode network supporting creative processes and the frontoparietal network, which includes lateral prefrontal brain areas like the dorsolateral prefrontal cortex, dorsal premotor cortex and inferior frontal gyrus, supporting evaluative processes (Belden et al., 2020). Given that this time window leads up to a right handed response, we think that this effect could be related to motor planning, which would imply that when improvisers are merely asked to respond to an “in-class” chord, they co-engage the default mode network pointing toward a context of this motor response that is biased toward creativity. Assuming a more ecologically valid context, this tight integration with the default mode network could enable a more direct and flexible access to musical structures and motor patterns which would seem conducive to greater mastery in musical improvisation. Post response, improvisers exhibited greater connectivity between the cognitive control and the left frontoparietal network which may be reflective of evaluative processes. The fact that we found no effects for the left frontoparietal network in association with classical training, supports the idea that the left frontoparietal network plays a particular role for improvisers here in this experiment and potentially more generally in more ecologically valid contexts.

In the same time window leading up to the right handed response, improvisers further showed greater connectivity between the cognitive control and the right frontoparietal network.

The cognitive control network is thought to be a superordinate network that supports executive control functions (Cole & Schneider, 2007; Niendam et al., 2012). As Cole and Schneider explain, this may include vigilance or sustained attention (Pennington & Ozonoff, 1996; Smith & Jonides, 1999), initiation of complex goal-directed behaviors (Lezak, 1995), inhibition of prepotent but incorrect responses (Smith & Jonides, 1999; Luna et al., 2010), flexibility to shift easily between goal states (Ravizza & Carter, 2008), planning necessary steps to achieve goal (Smith & Jonides, 1999) and the ability to hold information in working memory and to manipulate the information to guide response selection (Goldman-Rakic, 1996).

Since at this point in the trial, improvisers have not yet performed a motor action, it does not seem plausible that this phenomenon is related to an evaluative process in accordance with the dual-process model of creative cognition, even though cognitive control structures are involved. Thus it seems more likely that this phenomenon, which at this time point is specific to improvisers is also related to motor planning.

After the manual response, improvisers showed greater connectivity between networks with the dorsal attention and visual network acting as hubs and consistent effects were being also observed between salience and dorsal attention related networks.

The function of the dorsal attention network has been described as mediating top-down guided voluntary allocation of (primarily visual) attention to locations or features (Vossel et al., 2014) or the endogenous deployment of attention (Corbetta & Shulman, 2002), while Marek & Dosenbach (2018) suggest it may play a more general role in adaptive task control. The dorsal attention network has been found to be activated during voluntary attention shifts during search for salient visual stimuli (Shulman et al., 2003) and more recent findings indicate that the dorsal attention network may also play a role in external attention, either independently or in task-dependent interaction with the ventral attention network (Ahrens et al., 2019). The ventral attention network has been associated with (exogenous) re-orienting towards task-relevant events that appear at unexpected locations (Ahrens et al., 2019; Corbetta & Shulman, 2002). In experimental design, predictive (symbolic) cues are usually used to engage endogenous attention, as opposed to transient/non-predictive events to test exogenous attention (Ahrens et al., 2019).

One potential explanation of the observed effects around the dorsal attention network could be that for improvisers, a situation where the musician merely responds to an “in-class” chord triggers increased deployment of endogenous attention. To an improviser an “in-class” chord, particularly in the context of this experiment (where such chords are rare) but maybe more generally even during performance could represent something akin to a predictive cue. The increased engagement of endogenous attention could be linked to processes that are vital for successful improvisation. For example, what is the harmony or functional class of this chord I just heard and what is a suitable, adaptive response right now (i.e. for pressing the button in the experiment or playing the next tone or chord during performance). Major parts of the dorsal attention network also overlap the right parietal areas where Rosen et al. (2020) found greater power to be associated with greater improvisation experience. As potential explanations these authors referenced processes related to multimodal sensory processing and integration (Mihaly, 1996), long-term memory access (Wagner et al., 2005) or spatial coding, sensory-motor transformation and attention (Kaas & Stepniewska, 2016).

Musicians with greater experience in classical performance, but particularly those who were also improvisers consistently showed effects indicating decreased engagement and integration of the cognitive control network, specifically, lower connectivity and logarithmic power within the cognitive control network as well as lower connectivity between the cognitive control network and other networks like default mode, right frontoparietal, dorsal attention and visual network. This also means, that improvisers with particularly little experience in training classical music showed particularly high reliance on and integration of the cognitive control network after the manual response. This is for the most part consistent with what we find in terms of significant effects related to improvisation experience.

What we observe here may be an interaction effect between training in improvisation and classical music, such that improvisers with particularly little experience in training classical music require greater engagement of the cognitive control network to determine whether the response was accurate. One possible explanation for why this could be the case, could be that improvisers more so than classically trained musicians engage cognitive control resources after the response as an evaluative behavior consistent with the dual-process theory of cognition toward creative behavior (Belden et al., 2020; Sowden et al., 2015). According to this idea creative behaviors may be implemented by alternating between generative and evaluative behaviors (Belden et al., 2020). These generative behaviors are thought to be spontaneous and intuitive (Belden et al., 2020) and referred to more formally as system 1 (Stanovich, 1999) or type 1 (Evans, 2008, 2009) processes. Evaluative behaviors on the other hand are thought to be related to deliberate and analytical processing and referred to more formally as system 2 (Stanovich, 1999) or type 2 processes (Evans, 2008, 2009). Improvisers are strongly conditioned to engage evaluative processes after actions. Classically trained musicians on the other hand, usually already know exactly what they are going to play. This makes it less important for classically trained musicians to evaluate the output they just generated.

Among the earliest effects, directly following the onset of the exemplar deviant chord, improvisers showed greater power in the default mode network, while classically trained musicians showed greater connectivity within the default mode network. This could be indicative of processes related to early memory retrieval, that are engaged more intensively the more intensely the musicians has been trained irrespective of discipline. Greater connectivity within network for classically trained musicians aligns with previous findings of greater local efficiency for classically trained musicians (Belden et al., 2020), while greater gamma power for improvisers could be a result of greater cortical thickness in areas of the default mode network which has been found for musical improvisers (Kühn et al., 2014).

Musicians who trained more extensively, irrespective of musical domain further showed lower connectivity between cognitive control and right frontoparietal network, with improvisers also showing lower connectivity but greater power in gamma within the frontoparietal network and classically trained musicians showing greater connectivity within the cognitive control network. Taken together these findings point to a difference in executive control processes between the types of musical disciplines when faced with an exemplar deviant. While classically trained musicians seem to more strongly engage cognitive control resources, again exhibiting stronger within-network connectivity suggestive of high local efficiency (Belden et al., 2020), improvisers in contrast, showed lower connectivity and again greater power within the right frontoparietal network, which could be linked to more globally connected cortical organization (Belden et al., 2020).

Another difference between the two types of training directly after perceiving an exemplar deviant, may lie in how salience related networks configure dorsal attention related networks, with improvisers showing less connectivity between cingulo opercular and dorsal attention network as well as between the salience and visual network. Classically trained musicians on the other hand showed greater connectivity between dorsal attention and both cingulo opercular and the salience network. In accordance with our hypothesis (Goldman et al., 2020), this could be interpreted as improvisers perceiving the exemplar deviant as more similar to the standard since both chords are constituents of the same functional class. For more extensively trained classical musicians on the other hand, their training may make them more sensitive to the subtle difference between exemplar and function deviant, which in turn leads salience related networks to more strongly engage processes related to endogenous attention.

Interpreting the involvement of the visual network should take into account that musicians in this experiment were performing a target detection task, for which Mantini et al. (2009) showed, based on simultaneously recorded EEG and BOLD data, that activity in the dorsal and ventral attention network correlated significantly with the P300 reference time course and thus was interpreted to best account for sustained and transient activity in a visual oddball task. Thus one could consider as an alternative explanation that improvisers may have been merely more surprised for the exemplar, relative to the function deviant for an unknown reason other than our manipulation related to categorization of musical structures. But this would not explain the increased connectivity between default mode and visual network. On the contrary, connectivity between cortical networks, particularly also including the default mode network has been robustly linked to improvisation, particularly at rest (Belden et al., 2020).

Behavioral effects were mostly found in the alpha, beta and gamma band and were more numerous than effects related to either of the two types of musical expertise. Apart from very few exceptions the nature of associations was such that greater connectivity between or within the networks was associated with slower and less accurate responding to exemplar relative to function deviants. Overall this is in line with previous work that also observed links between behavioral performance and connectivity within and between brain networks as reviewed by Cohen (2018). One hypothesis for this experiment was that improvisers would respond slower and less accurately for exemplar relative to function deviants. This holds, in that we found effects in behavior that matched - in time, frequency and direction - those effects that were most convincingly tied to improvisation expertise. However, we also found behavioral effects that matched effects related to expertise in classical music, supporting the idea that more intense training in the classical domain, may as well decrease task performance for exemplar deviants, likely for reasons different from those found in improvisers. In addition we found behavioral effects for which we found no corresponding effects for training in improvisation or classical music, which could mean that these behavioral effects capture phenomena unrelated to musical expertise, or that there is a matching effect related to musical expertise, but that self-reporting is too noisy to establish a significant effect. Any other mismatch between effects found for behavior and self-reported experience could be a result of behavioral effects being strongly tied to motor-related brain activity, while effects for self-reported experience may be more strongly related to cognitive aspects. In summary, given that our experimental manipulation strongly narrows resulting effects for exemplar relative to function deviants to categorization of musical structures, and that we find behavioral effects that match the effects that were most strongly tied to self-reported improvisation expertise, we think we found robust evidence in support of the idea that categorization of musical structures is tied to how large-scale cortical brain networks are engaged and interact, and that improvisers implement these processes differently compared to classically trained musicians. While we found these effects here in a target detection task, we argue, supported by literature, that these or similar mechanisms may be employed when musicians actually improvise on their instrument, may facilitate improvisation as a skill and should be a result of improvisers’ intense and specific training regimen.

## Acknowledgments

This work was supported in part by the United States Army Research Laboratory under Cooperative Agreements W911NF-10-2-0022 and W911NF-16-2-0008 as well as a Vannevar Bush Faculty Fellowship from the US Department of Defense (N00014-20-1-2027) and the Presidential Scholars in Society and Neuroscience program at Columbia University. Author contributions: J.F., J.M., A.G., Y.L. and P.S. designed research; J.F., Y.L. and J.M. performed research; J.F. and Y.L. contributed analytic tools; J.F. and Y.L. analyzed data; J.F., P.S., Y.L., A.G. and J.M. wrote the paper.

