## Supplemental Information for "Spatiospectral brain networks reflective of improvisational experience"

### 1. Effects within brain networks

#### 1.1. Overview of within network effects

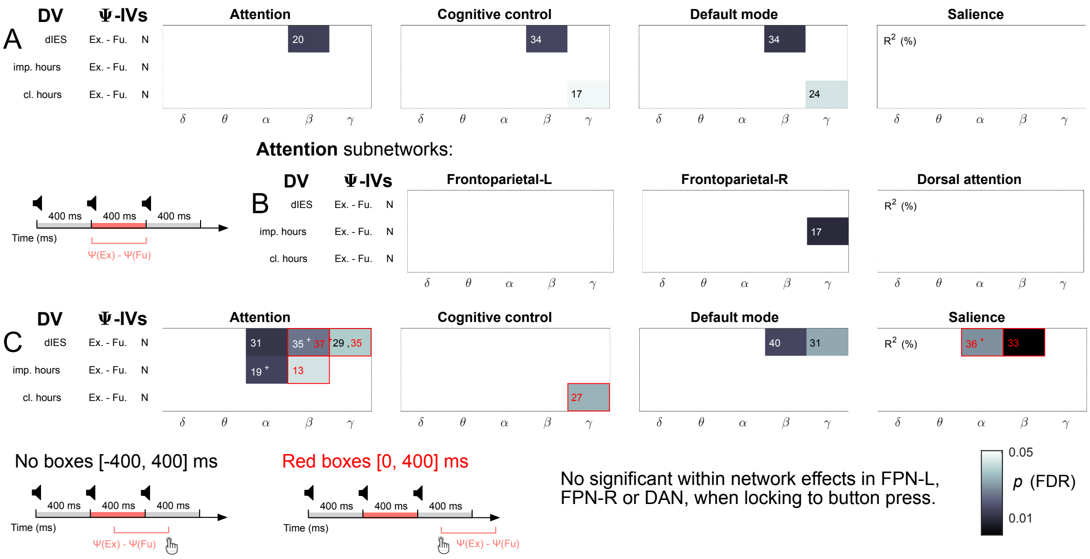


Figure 1: Overview of model level effects within brain networks. (A) Post-stimulus connectivity results for the four top-level networks. (B) Again, post-stimulus results, but for the subnetworks of the compound attention network. The subnetworks of the compound attention network were investigated since statistically significant effects were found for the dependent improviation hours pre and post button press, see panel (C).

### *1.2. Within network effects - scatter plots*

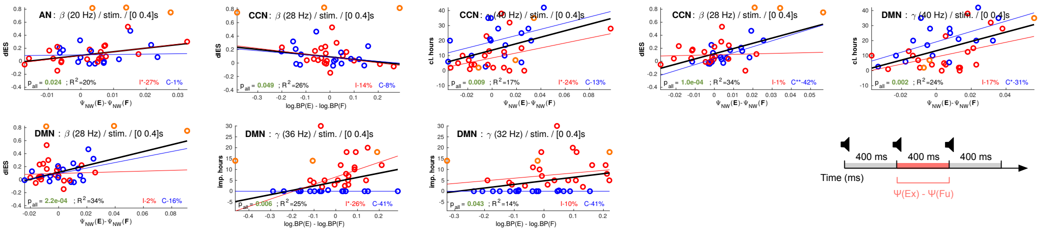

Figure 2: Regression analyses for significant model-level effects within the four top-level networks, post-stimulus (stim.). The black line represents a fit for all musicians, while the red and blue lines represent fits to sub-groups of improvisers (red rings; weekly improvisation hours $\geq 0.5$) and classically trained performers (blue rings; weekly improvisation hours $< 0.5$). For fits on subgroups one, two and three * represent p-value levels of 0.05, 0.01 and 0.001 respectively.

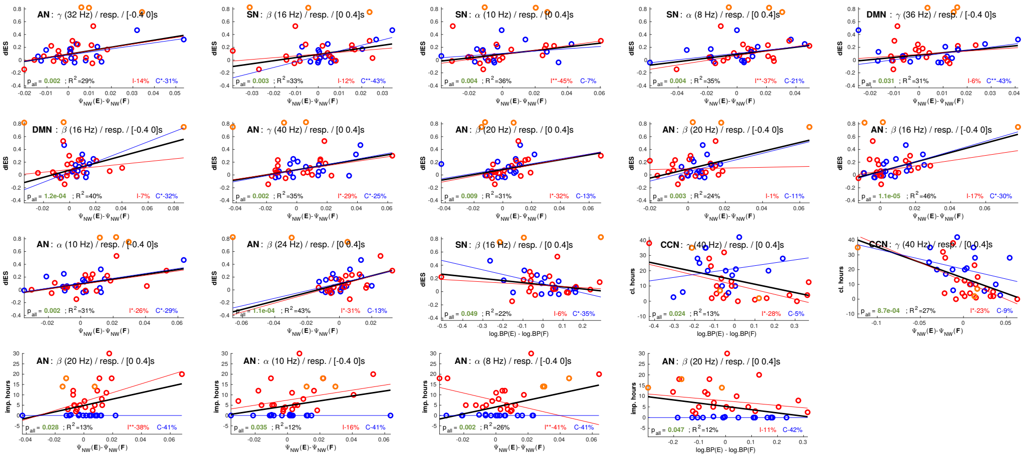

Figure 3: Regression analyses for significant model-level effects within the four top-level networks, response-locked (resp.). The black line represents a fit for all musicians, while the red and blue lines represent fits to sub-groups of improvisers (red rings; weekly improvisation hours $\geq 0.5$) and classically trained performers (blue rings; weekly improvisation hours $< 0.5$). For fits on subgroups one, two and three * represent p-value levels of 0.05, 0.01 and 0.001 respectively.

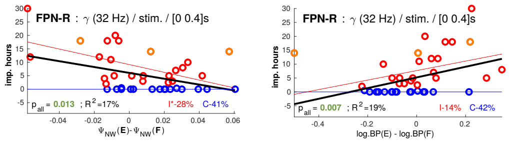


Figure 4: Regression analyses for significant model-level effects within the compound attention network, post-stimulus (stim.). The black line represents a fit for all musicians, while the red and blue lines represent fits to sub-groups of improvisers (red rings; weekly improvisation hours $\geq 0.5$) and classically trained performers (blue rings; weekly improvisation hours $< 0.5$). For fits on subgroups one, two and three * represent p-value levels of 0.05, 0.01 and 0.001 respectively.

| Locked to | Network | Time window | DV | IV | Frequency band | All musicians $p_{FDR}$ | $R^2$ |
|---|---|---|---|---|---|---|---|
| chord | AN | [0, 400] | dIES | $[\Psi_{NW}(E) - \Psi_{NW}(F)] + [log.BP(E) - log.BP(F)]$ | $\beta_2$ | 0.015 | 33.1 % |
| chord | CCN | [0, 400] | dIES | $[\Psi_{NW}(E) - \Psi_{NW}(F)] + [log.BP(E) - log.BP(F)]$ | $\beta_4$ | 0.016 | 32.1 % |
| chord | CCN | [0, 400] | cl. hours | $[\Psi_{NW}(E) - \Psi_{NW}(F)] + [log.BP(E) - log.BP(F)]$ | $\gamma_3$ | 0.049 | 23.5 % |
| chord | DMN | [0, 400] | dIES | $[\Psi_{NW}(E) - \Psi_{NW}(F)] + [log.BP(E) - log.BP(F)]$ | $\beta_4$ | 0.013 | 35.4 % |
| chord | DMN | [0, 400] | cl. hours | $[\Psi_{NW}(E) - \Psi_{NW}(F)] + [log.BP(E) - log.BP(F)]$ | $\gamma_3$ | 0.045 | 24.8 % |
| response | AN | [-400, 0] | dIES | $[\Psi_{NW}(E) - \Psi_{NW}(F)] + [log.BP(E) - log.BP(F)]$ | $\alpha_2$ | 0.013 | 34.1 % |
| response | AN | [-400, 0] | dIES | $[\Psi_{NW}(E) - \Psi_{NW}(F)] + [log.BP(E) - log.BP(F)]$ | $\beta_1$ | 4.94e-04 | 48.7 % |
| response | AN | [-400, 0] | dIES | $[\Psi_{NW}(E) - \Psi_{NW}(F)] + [log.BP(E) - log.BP(F)]$ | $\beta_2$ | 0.016 | 31.5 % |
| response | AN | [-400, 0] | dIES | $[\Psi_{NW}(E) - \Psi_{NW}(F)] + [log.BP(E) - log.BP(F)]$ | $\gamma_1$ | 0.039 | 26.1 % |
| response | AN | [-400, 0] | imp. hours | $[\Psi_{NW}(E) - \Psi_{NW}(F)] + [log.BP(E) - log.BP(F)]$ | $\alpha_1$ | 0.005 | 39.7 % |
| response | AN | [-400, 0] | imp. hours | $[\Psi_{NW}(E) - \Psi_{NW}(F)] + [log.BP(E) - log.BP(F)]$ | $\alpha_2$ | 0.026 | 29.3 % |
| response | AN | [0, 400] | dIES | $[\Psi_{NW}(E) - \Psi_{NW}(F)] + [log.BP(E) - log.BP(F)]$ | $\beta_2$ | 0.045 | 24.5 % |
| response | AN | [0, 400] | dIES | $[\Psi_{NW}(E) - \Psi_{NW}(F)] + [log.BP(E) - log.BP(F)]$ | $\beta_3$ | 0.005 | 40.1 % |
| response | AN | [0, 400] | dIES | $[\Psi_{NW}(E) - \Psi_{NW}(F)] + [log.BP(E) - log.BP(F)]$ | $\gamma_3$ | 0.033 | 27.4 % |
| response | AN | [0, 400] | imp. hours | $[\Psi_{NW}(E) - \Psi_{NW}(F)] + [log.BP(E) - log.BP(F)]$ | $\beta_2$ | 0.044 | 25.1 % |
| response | CCN | [0, 400] | cl. hours | $[\Psi_{NW}(E) - \Psi_{NW}(F)] + [log.BP(E) - log.BP(F)]$ | $\gamma_3$ | 0.035 | 26.8 % |
| response | DMN | [-400, 0] | dIES | $[\Psi_{NW}(E) - \Psi_{NW}(F)] + [log.BP(E) - log.BP(F)]$ | $\beta_1$ | 0.016 | 32.0 % |
| response | DMN | [-400, 0] | dIES | $[\Psi_{NW}(E) - \Psi_{NW}(F)] + [log.BP(E) - log.BP(F)]$ | $\gamma_2$ | 0.033 | 27.3 % |
| response | SN | [0, 400] | dIES | $[\Psi_{NW}(E) - \Psi_{NW}(F)] + [log.BP(E) - log.BP(F)]$ | $\alpha_1$ | 0.026 | 29.1 % |
| response | SN | [0, 400] | dIES | $[\Psi_{NW}(E) - \Psi_{NW}(F)] + [log.BP(E) - log.BP(F)]$ | $\alpha_2$ | 0.032 | 27.9 % |
| response | SN | [0, 400] | dIES | $[\Psi_{NW}(E) - \Psi_{NW}(F)] + [log.BP(E) - log.BP(F)]$ | $\beta_1$ | 1.15e-04 | 54.5 % |
| chord | FPN-R | [0, 400] | imp. hours | $[\Psi_{NW}(E) - \Psi_{NW}(F)] + [log.BP(E) - log.BP(F)]$ | $\gamma_1$ | 0.010 | 40.8 % |
| chord | DAN-core | [0, 400] | dIES | $[\Psi_{NW}(E) - \Psi_{NW}(F)] + [log.BP(E) - log.BP(F)]$ | $\beta_4$ | 0.018 | 31.5 % |
| chord | DAN-core | [0, 400] | cl. hours | $[\Psi_{NW}(E) - \Psi_{NW}(F)] + [log.BP(E) - log.BP(F)]$ | $\gamma_3$ | 0.018 | 31.8 % |
| response | DAN-core | [-400, 0] | dIES | $[\Psi_{NW}(E) - \Psi_{NW}(F)] + [log.BP(E) - log.BP(F)]$ | $\gamma_2$ | 0.050 | 26.8 % |
| response | VN | [-400, 0] | imp. hours | $[\Psi_{NW}(E) - \Psi_{NW}(F)] + [log.BP(E) - log.BP(F)]$ | $\alpha_3$ | 0.018 | 31.9 % |
| response | VN | [0, 400] | imp. hours | $[\Psi_{NW}(E) - \Psi_{NW}(F)] + [log.BP(E) - log.BP(F)]$ | $\gamma_1$ | 0.013 | 38.4 % |

Table 1: Details for significant findings within networks on model level (corresponds to overview plots). DV indicates the dependent variable, IV the independent variables, where an expression within square brackets represents one factor.

| Locked to | Network | Time window | DV | IV | Frequency band | All musicians p | All musicians $R^2$ | Improvisers p | Improvisers $R^2$ | Classical performers p | Classical performers $R^2$ |
|---|---|---|---|---|---|---|---|---|---|---|---|
| chord | AN | [0, 400] | dIES | $\Psi_{NW}(E) - \Psi_{NW}(F)$ | $\beta_2$ | 0.024 | 19.9 % | 0.021 | 26.8 % | 0.891 | 0.7 % |
| chord | CCN | [0, 400] | dIES | $\Psi_{NW}(E) - \Psi_{NW}(F)$ | $\beta_4$ | 9.96e-05 | 34.4 % | 0.811 | 1.1 % | 0.007 | 42.5 % |
| chord | CCN | [0, 400] | dIES | $log.BP(E) - log.BP(F)$ | $\beta_4$ | 0.049 | 25.9 % | 0.152 | 13.7 % | 0.348 | 7.7 % |
| chord | CCN | [0, 400] | cl. hours | $\Psi_{NW}(E) - \Psi_{NW}(F)$ | $\gamma_3$ | 0.009 | 17.2 % | 0.039 | 24.4 % | 0.169 | 13.1 % |
| chord | DMN | [0, 400] | imp. hours | $log.BP(E) - log.BP(F)$ | $\gamma_1$ | 0.043 | 13.5 % | 0.365 | 10.1 % | 1.000 | 40.6 % |
| chord | DMN | [0, 400] | imp. hours | $log.BP(E) - log.BP(F)$ | $\gamma_2$ | 0.006 | 24.6 % | 0.042 | 25.8 % | 1.000 | 40.8 % |
| chord | DMN | [0, 400] | dIES | $\Psi_{NW}(E) - \Psi_{NW}(F)$ | $\beta_4$ | 2.24e-04 | 33.7 % | 0.838 | 1.7 % | 0.121 | 16.4 % |
| chord | DMN | [0, 400] | cl. hours | $\Psi_{NW}(E) - \Psi_{NW}(F)$ | $\gamma_3$ | 0.002 | 23.9 % | 0.075 | 16.7 % | 0.025 | 31.2 % |
| response | AN | [-400, 0] | imp. hours | $\Psi_{NW}(E) - \Psi_{NW}(F)$ | $\alpha_1$ | 0.002 | 26.3 % | 0.009 | 41.5 % | 1.000 | 41.4 % |
| response | AN | [-400, 0] | imp. hours | $\Psi_{NW}(E) - \Psi_{NW}(F)$ | $\alpha_2$ | 0.035 | 12.3 % | 0.121 | 16.4 % | 1.000 | 40.7 % |
| response | AN | [-400, 0] | dIES | $\Psi_{NW}(E) - \Psi_{NW}(F)$ | $\alpha_2$ | 0.002 | 31.2 % | 0.023 | 25.8 % | 0.033 | 28.6 % |
| response | AN | [-400, 0] | dIES | $\Psi_{NW}(E) - \Psi_{NW}(F)$ | $\beta_1$ | 1.13e-05 | 46.1 % | 0.133 | 16.6 % | 0.030 | 29.7 % |
| response | AN | [-400, 0] | dIES | $\Psi_{NW}(E) - \Psi_{NW}(F)$ | $\beta_2$ | 0.003 | 24.0 % | 0.849 | 1.0 % | 0.287 | 10.8 % |
| response | AN | [-400, 0] | dIES | $\Psi_{NW}(E) - \Psi_{NW}(F)$ | $\gamma_1$ | 0.002 | 29.1 % | 0.112 | 14.1 % | 0.026 | 30.6 % |
| response | AN | [0, 400] | imp. hours | $\Psi_{NW}(E) - \Psi_{NW}(F)$ | $\beta_2$ | 0.028 | 13.2 % | 0.005 | 38.2 % | 1.000 | 41.1 % |
| response | AN | [0, 400] | imp. hours | $log.BP(E) - log.BP(F)$ | $\beta_2$ | 0.047 | 11.7 % | 0.308 | 11.1 % | 1.000 | 41.5 % |
| response | AN | [0, 400] | dIES | $\Psi_{NW}(E) - \Psi_{NW}(F)$ | $\beta_2$ | 0.009 | 31.4 % | 0.010 | 31.9 % | 0.173 | 13.3 % |
| response | AN | [0, 400] | dIES | $\Psi_{NW}(E) - \Psi_{NW}(F)$ | $\beta_3$ | 1.07e-04 | 43.4 % | 0.011 | 30.7 % | 0.169 | 13.2 % |
| response | AN | [0, 400] | dIES | $\Psi_{NW}(E) - \Psi_{NW}(F)$ | $\gamma_3$ | 0.002 | 35.2 % | 0.017 | 29.4 % | 0.047 | 25.3 % |
| response | CCN | [0, 400] | cl. hours | $\Psi_{NW}(E) - \Psi_{NW}(F)$ | $\gamma_3$ | 8.72e-04 | 26.8 % | 0.032 | 23.2 % | 0.251 | 9.3 % |
| response | CCN | [0, 400] | cl. hours | $log.BP(E) - log.BP(F)$ | $\gamma_3$ | 0.024 | 13.0 % | 0.017 | 27.7 % | 0.411 | 4.9 % |
| response | DMN | [-400, 0] | dIES | $\Psi_{NW}(E) - \Psi_{NW}(F)$ | $\beta_1$ | 1.18e-04 | 40.3 % | 0.343 | 6.6 % | 0.036 | 31.5 % |
| response | DMN | [-400, 0] | dIES | $\Psi_{NW}(E) - \Psi_{NW}(F)$ | $\gamma_2$ | 0.031 | 31.3 % | 0.424 | 5.7 % | 0.009 | 43.4 % |
| response | SN | [0, 400] | dIES | $\Psi_{NW}(E) - \Psi_{NW}(F)$ | $\alpha_1$ | 0.004 | 35.2 % | 0.006 | 36.9 % | 0.076 | 21.4 % |
| response | SN | [0, 400] | dIES | $\Psi_{NW}(E) - \Psi_{NW}(F)$ | $\alpha_2$ | 0.004 | 36.3 % | 0.002 | 44.6 % | 0.325 | 7.4 % |
| response | SN | [0, 400] | dIES | $\Psi_{NW}(E) - \Psi_{NW}(F)$ | $\beta_1$ | 0.003 | 33.1 % | 0.157 | 12.3 % | 0.006 | 42.8 % |
| response | SN | [0, 400] | dIES | $log.BP(E) - log.BP(F)$ | $\beta_1$ | 0.049 | 22.5 % | 0.364 | 5.8 % | 0.015 | 35.4 % |
| chord | FPN-R | [0, 400] | imp. hours | $\Psi_{NW}(E) - \Psi_{NW}(F)$ | $\gamma_1$ | 0.013 | 16.7 % | 0.017 | 27.7 % | 1.000 | 41.2 % |
| chord | FPN-R | [0, 400] | imp. hours | $log.BP(E) - log.BP(F)$ | $\gamma_1$ | 0.007 | 19.4 % | 0.133 | 13.8 % | 1.000 | 41.9 % |
| chord | DAN-core | [0, 400] | dIES | $\Psi_{NW}(E) - \Psi_{NW}(F)$ | $\beta_4$ | 6.36e-05 | 36.4 % | 0.228 | 8.0 % | 0.037 | 27.6 % |
| chord | DAN-core | [0, 400] | cl. hours | $\Psi_{NW}(E) - \Psi_{NW}(F)$ | $\gamma_3$ | 0.005 | 19.8 % | 0.068 | 17.6 % | 0.191 | 11.9 % |
| response | DAN-core | [-400, 0] | dIES | $\Psi_{NW}(E) - \Psi_{NW}(F)$ | $\gamma_2$ | 0.004 | 32.9 % | 0.248 | 10.2 % | 0.005 | 43.8 % |
| response | VN | [-400, 0] | imp. hours | $\Psi_{NW}(E) - \Psi_{NW}(F)$ | $\alpha_3$ | 5.42e-04 | 29.3 % | 0.033 | 24.5 % | 1.000 | 40.6 % |
| response | VN | [0, 400] | imp. hours | $\Psi_{NW}(E) - \Psi_{NW}(F)$ | $\gamma_1$ | 2.30e-04 | 46.6 % | 0.916 | 3.9 % | 1.000 | 43.8 % |
| response | VN | [0, 400] | cl. hours | $log.BP(E) - log.BP(F)$ | $\theta$ | 0.004 | 20.2 % | 0.008 | 33.2 % | 0.122 | 16.2 % |

Table 2: Details for significant findings within networks on the level of factors (corresponds to scatter plots). DV indicates the dependent variable, IV the independent variables, where an expression within square brackets represents one factor.

### 2. Effects between brain networks

#### *2.1. Overview of between network effects*

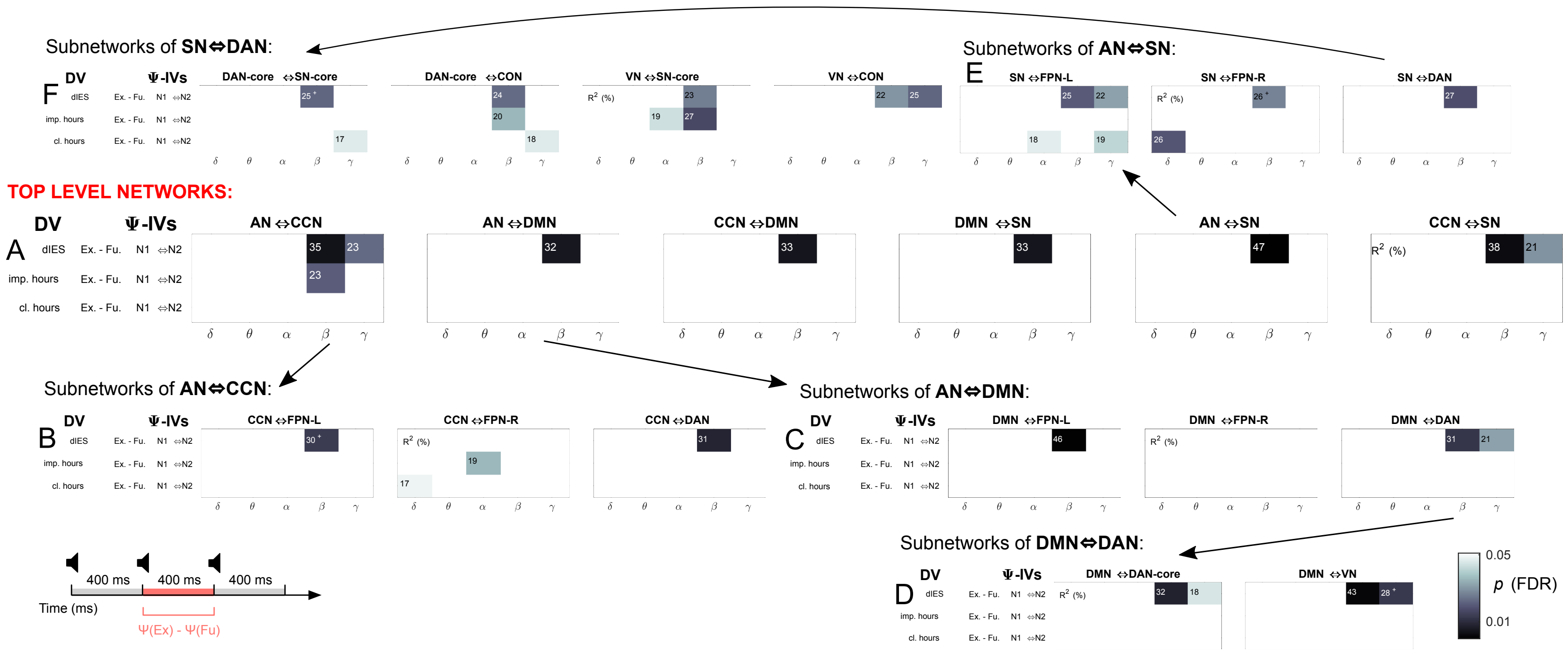


Figure 5: Overview of model level effects between brain networks, post-stimulus. (A) Connectivity results between the four top-level networks (so six plots). Panels (B) to (F) show results for subnetworks that were explored as a result of significant findings at higher network levels for dependent variables weekly hours spent training improvisation or classical performance.

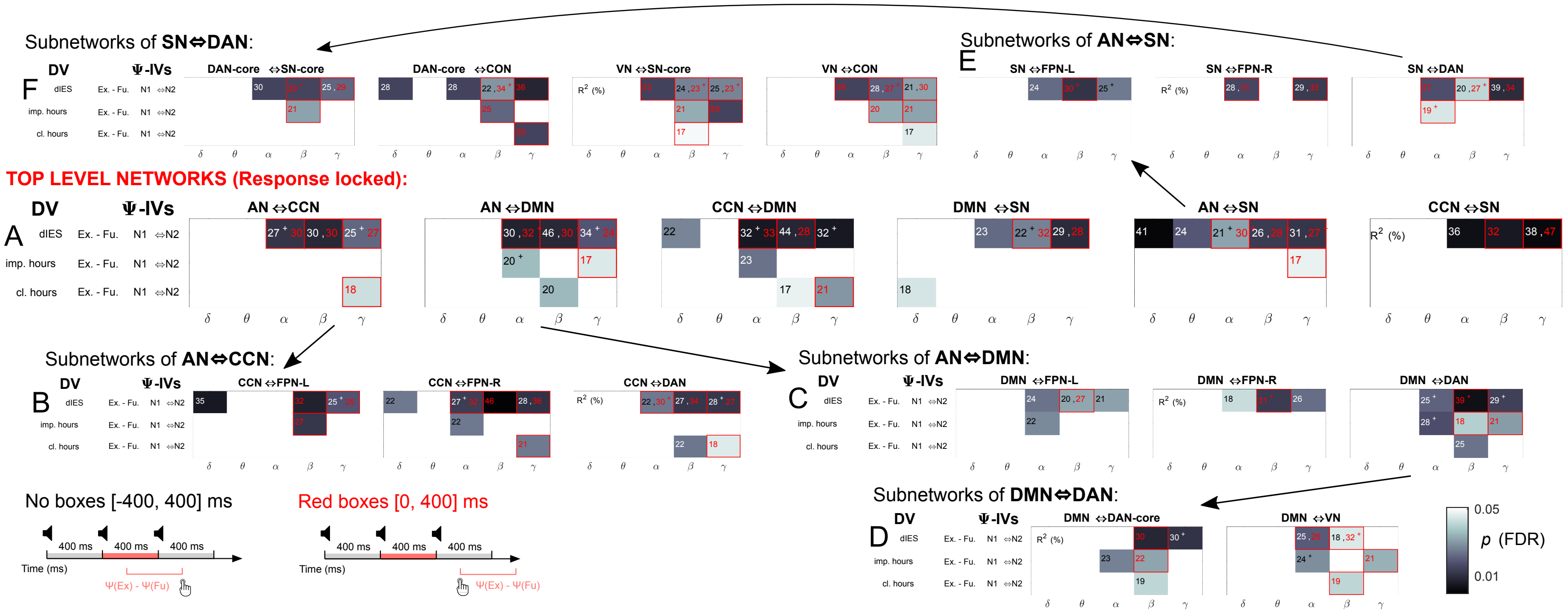

Figure 6: Overview of model level effects between brain networks, response locked. (A) Connectivity results between the four top-level networks (so six plots). Panels (B) to (F) show results for subnetworks that were explored as a result of significant findings at higher network levels for the dependent variables weekly hours spent training improvisation or classical performance.

### *2.2. Between network effects - scatter plots*

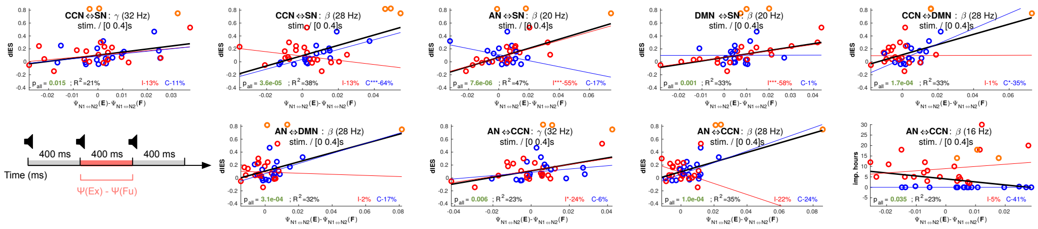

Figure 7: Regression analyses for significant model-level effects between the four top-level networks, post-stimulus (stim.). The black line represents a fit for all musicians, while the red and blue lines represent fits to sub-groups of improvisers (red rings; weekly improvisation hours $\geq 0.5$) and classically trained performers (blue rings; weekly improvisation hours $< 0.5$). For fits on subgroups one, two and three * represent p-value levels of 0.05, 0.01 and 0.001 respectively.

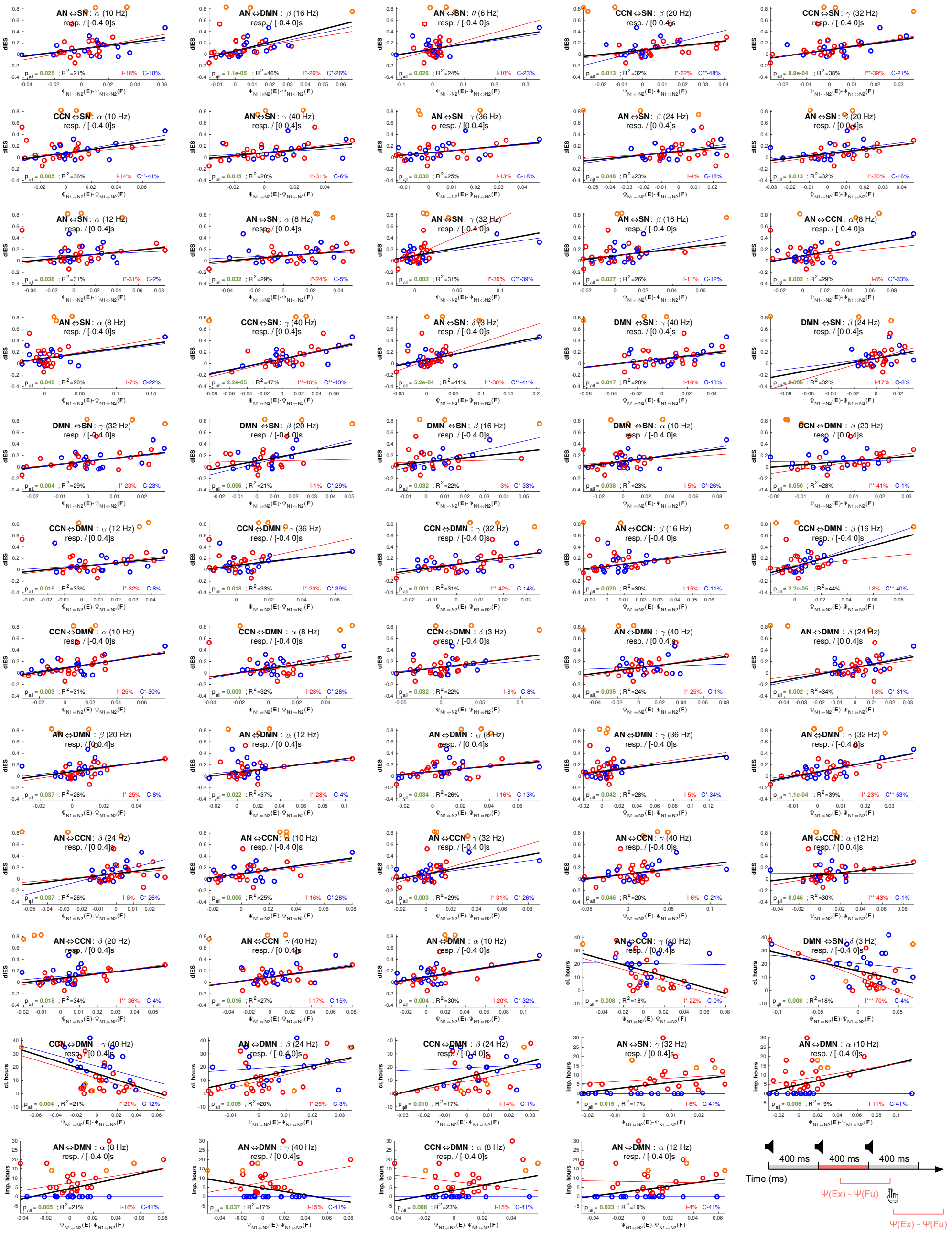


Figure 8: Regression analyses for significant model-level effects between the four top-level networks, response-locked (resp.). The black line represents a fit for all musicians, while the red and blue lines represent fits to sub-groups of improvisers (red rings; weekly improvisation hours $\geq 0.5$) and classically trained performers (blue rings; weekly improvisation hours $< 0.5$). For fits on subgroups one, two and three * represent p-value levels of 0.05, 0.01 and 0.001 respectively.

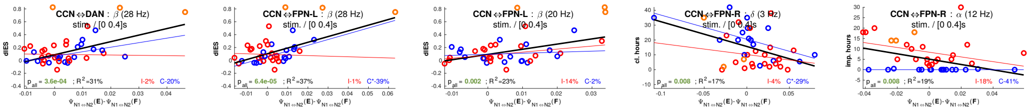

Figure 9: Regression analyses for significant model-level effects between subnetworks of the compound attention network and the cognitive control network, post-stimulus (stim.). The black line represents a fit for all musicians, while the red and blue lines represent fits to sub-groups of improvisers (red rings; weekly improvisation hours $\geq 0.5$) and classically trained performers (blue rings; weekly improvisation hours $< 0.5$). For fits on subgroups one, two and three * represent p-value levels of 0.05, 0.01 and 0.001 respectively.

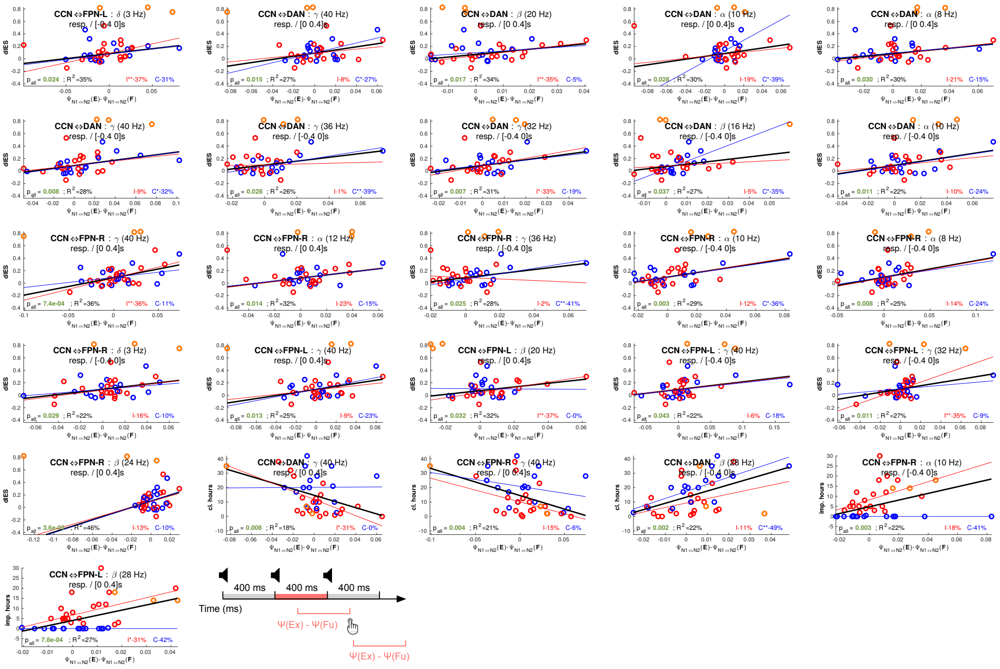

Figure 10: Regression analyses for significant model-level effects between subnetworks of the compound attention network and the cognitive control network, response-locked (resp.). The black line represents a fit for all musicians, while the red and blue lines represent fits to sub-groups of improvisers (red rings; weekly improvisation hours $\geq 0.5$) and classically trained performers (blue rings; weekly improvisation hours $< 0.5$). For fits on subgroups one, two and three * represent p-value levels of 0.05, 0.01 and 0.001 respectively.

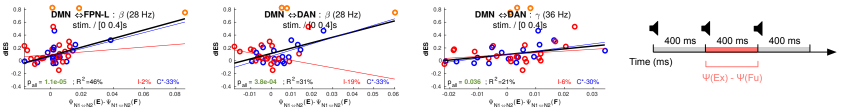

Figure 11: Regression analyses for significant model-level effects between subnetworks of the compound attention network and the default mode network, post-stimulus (stim.). The black line represents a fit for all musicians, while the red and blue lines represent fits to sub-groups of improvisers (red rings; weekly improvisation hours $\geq 0.5$) and classically trained performers (blue rings; weekly improvisation hours $< 0.5$). For fits on subgroups one, two and three * represent p-value levels of 0.05, 0.01 and 0.001 respectively.

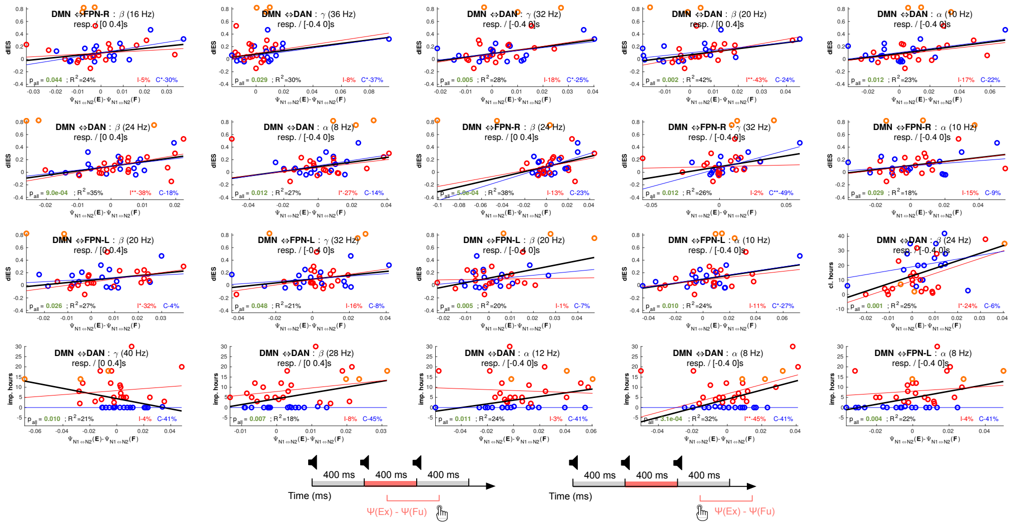

Figure 12: Regression analyses for significant model-level effects between subnetworks of the compound attention network and the default mode network, response-locked (resp.). The black line represents a fit for all musicians, while the red and blue lines represent fits to sub-groups of improvisers (red rings; weekly improvisation hours $\geq 0.5$) and classically trained performers (blue rings; weekly improvisation hours $< 0.5$). For fits on subgroups one, two and three * represent p-value levels of 0.05, 0.01 and 0.001 respectively.

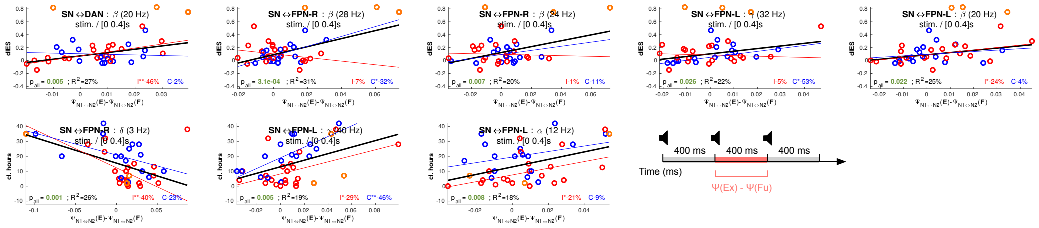

Figure 13: Regression analyses for significant model-level effects between subnetworks of the compound attention network and the salience network, post-stimulus (stim.). The black line represents a fit for all musicians, while the red and blue lines represent fits to sub-groups of improvisers (red rings; weekly improvisation hours $\geq 0.5$) and classically trained performers (blue rings; weekly improvisation hours $< 0.5$). For fits on subgroups one, two and three * represent p-value levels of 0.05, 0.01 and 0.001 respectively.

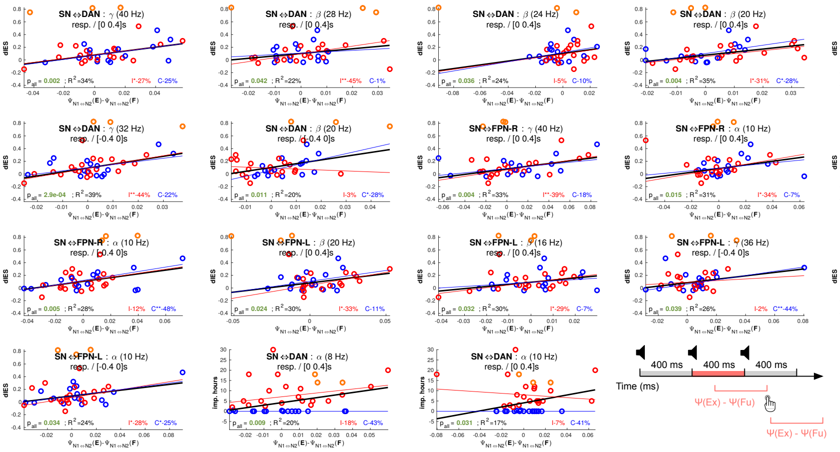


Figure 14: Regression analyses for significant model-level effects between subnetworks of the compound attention network and the salience network, response-locked (resp.). The black line represents a fit for all musicians, while the red and blue lines represent fits to sub-groups of improvisers (red rings; weekly improvisation hours $\geq 0.5$) and classically trained performers (blue rings; weekly improvisation hours $< 0.5$). For fits on subgroups one, two and three * represent p-value levels of 0.05, 0.01 and 0.001 respectively.

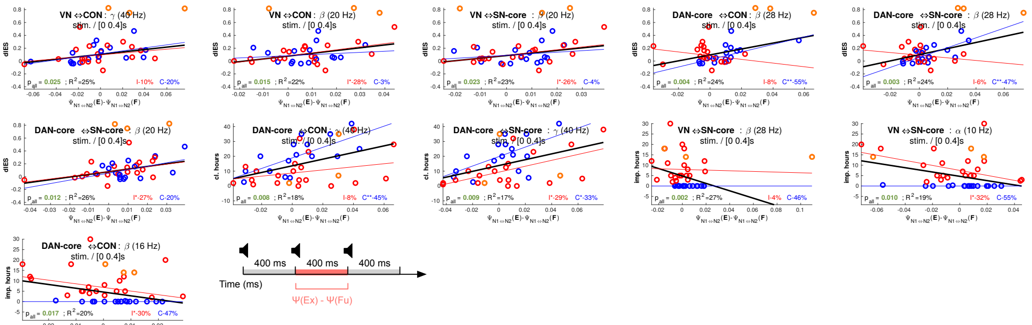


Figure 15: Regression analyses for significant model-level effects between subnetworks of the dorsal attention network and the salience network, post-stimulus (stim.). The black line represents a fit for all musicians, while the red and blue lines represent fits to sub-groups of improvisers (red rings; weekly improvisation hours $\geq 0.5$) and classically trained performers (blue rings; weekly improvisation hours $< 0.5$). For fits on subgroups one, two and three * represent p-value levels of 0.05, 0.01 and 0.001 respectively.

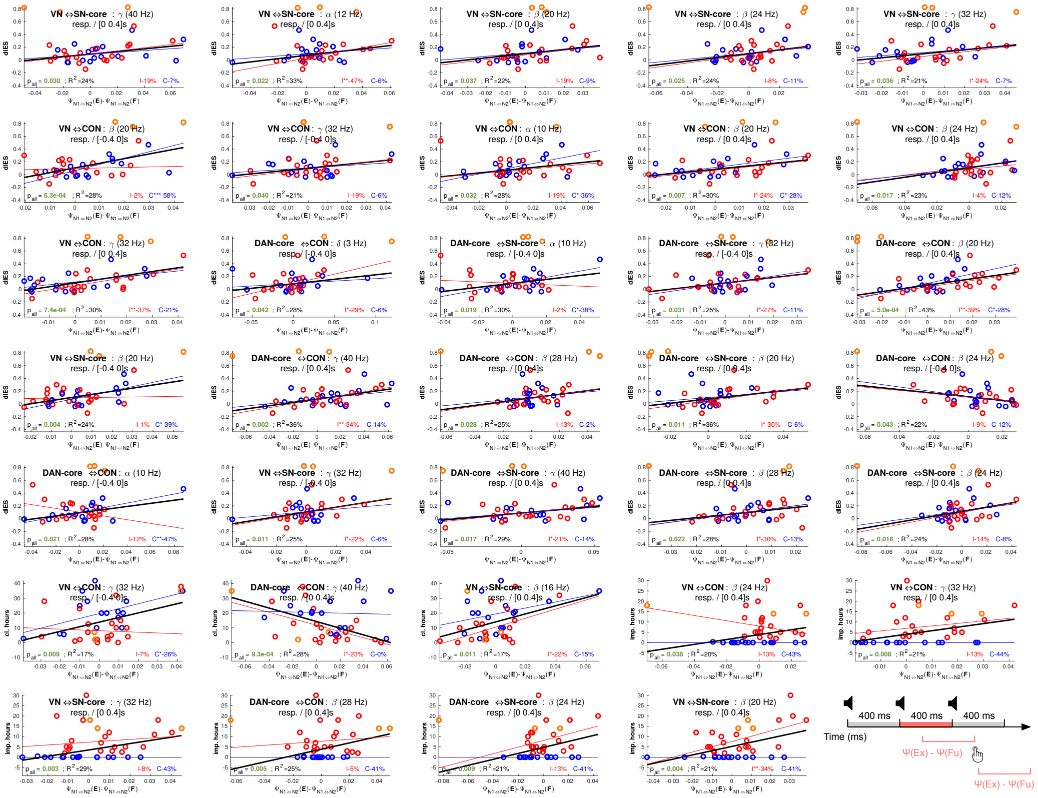


Figure 16: Regression analyses for significant model-level effects between subnetworks of the dorsal attention network and the salience network, response-locked (resp.). The black line represents a fit for all musicians, while the red and blue lines represent fits to sub-groups of improvisers (red rings; weekly improvisation hours $\geq 0.5$) and classically trained performers (blue rings; weekly improvisation hours $< 0.5$). For fits on subgroups one, two and three * represent p-value levels of 0.05, 0.01 and 0.001 respectively.

Table 3: Details for significant findings between networks on model level (corresponds to overview plots). DV indicates the dependent variable, IV the independent variables, where an expression within square brackets represents one factor.

| Locked to | Network | Time window | DV | IV | Frequency band | All musicians $p_{FDR}$ | All musicians $R^2$ |
|---|---|---|---|---|---|---|---|
| chord | AN ⇔CCN | [0, 400] | dIES | $\Psi_{N1\Leftrightarrow N2}(E) - \Psi_{N1\Leftrightarrow N2}(F)$ | $\beta_4$ | 0.004 | 33.3 % |
| chord | AN ⇔CCN | [0, 400] | dIES | $\Psi_{N1\Leftrightarrow N2}(E) - \Psi_{N1\Leftrightarrow N2}(F)$ | $\gamma_1$ | 0.023 | 20.7 % |
| chord | AN ⇔CCN | [0, 400] | imp. hours | $\Psi_{N1\Leftrightarrow N2}(E) - \Psi_{N1\Leftrightarrow N2}(F)$ | $\beta_1$ | 0.021 | 21.3 % |
| chord | AN ⇔DMN | [0, 400] | dIES | $\Psi_{N1\Leftrightarrow N2}(E) - \Psi_{N1\Leftrightarrow N2}(F)$ | $\beta_4$ | 0.005 | 30.5 % |
| chord | CCN ⇔DMN | [0, 400] | dIES | $\Psi_{N1\Leftrightarrow N2}(E) - \Psi_{N1\Leftrightarrow N2}(F)$ | $\beta_4$ | 0.005 | 31.5 % |
| chord | DMN ⇔SN | [0, 400] | dIES | $\Psi_{N1\Leftrightarrow N2}(E) - \Psi_{N1\Leftrightarrow N2}(F)$ | $\beta_2$ | 0.005 | 31.6 % |
| chord | AN ⇔SN | [0, 400] | dIES | $\Psi_{N1\Leftrightarrow N2}(E) - \Psi_{N1\Leftrightarrow N2}(F)$ | $\beta_2$ | 4.10e-04 | 45.4 % |
| chord | CCN ⇔SN | [0, 400] | dIES | $\Psi_{N1\Leftrightarrow N2}(E) - \Psi_{N1\Leftrightarrow N2}(F)$ | $\beta_4$ | 0.003 | 35.9 % |
| chord | CCN ⇔SN | [0, 400] | dIES | $\Psi_{N1\Leftrightarrow N2}(E) - \Psi_{N1\Leftrightarrow N2}(F)$ | $\gamma_1$ | 0.030 | 18.9 % |
| response | AN ⇔CCN | [-400, 0] | dIES | $\Psi_{N1\Leftrightarrow N2}(E) - \Psi_{N1\Leftrightarrow N2}(F)$ | $\alpha_1$ | 0.008 | 27.4 % |
| response | AN ⇔CCN | [-400, 0] | dIES | $\Psi_{N1\Leftrightarrow N2}(E) - \Psi_{N1\Leftrightarrow N2}(F)$ | $\alpha_2$ | 0.017 | 22.6 % |
| response | AN ⇔CCN | [-400, 0] | dIES | $\Psi_{N1\Leftrightarrow N2}(E) - \Psi_{N1\Leftrightarrow N2}(F)$ | $\beta_1$ | 0.007 | 28.1 % |
| response | AN ⇔CCN | [-400, 0] | dIES | $\Psi_{N1\Leftrightarrow N2}(E) - \Psi_{N1\Leftrightarrow N2}(F)$ | $\gamma_1$ | 0.008 | 27.5 % |
| response | AN ⇔CCN | [-400, 0] | dIES | $\Psi_{N1\Leftrightarrow N2}(E) - \Psi_{N1\Leftrightarrow N2}(F)$ | $\gamma_3$ | 0.037 | 17.3 % |
| response | AN ⇔CCN | [0, 400] | dIES | $\Psi_{N1\Leftrightarrow N2}(E) - \Psi_{N1\Leftrightarrow N2}(F)$ | $\alpha_3$ | 0.007 | 28.4 % |
| response | AN ⇔CCN | [0, 400] | dIES | $\Psi_{N1\Leftrightarrow N2}(E) - \Psi_{N1\Leftrightarrow N2}(F)$ | $\beta_2$ | 0.005 | 32.0 % |
| response | AN ⇔CCN | [0, 400] | dIES | $\Psi_{N1\Leftrightarrow N2}(E) - \Psi_{N1\Leftrightarrow N2}(F)$ | $\beta_3$ | 0.013 | 23.8 % |
| response | AN ⇔CCN | [0, 400] | dIES | $\Psi_{N1\Leftrightarrow N2}(E) - \Psi_{N1\Leftrightarrow N2}(F)$ | $\gamma_3$ | 0.012 | 24.6 % |
| response | AN ⇔CCN | [0, 400] | cl. hours | $\Psi_{N1\Leftrightarrow N2}(E) - \Psi_{N1\Leftrightarrow N2}(F)$ | $\gamma_3$ | 0.044 | 15.8 % |
| response | AN ⇔DMN | [-400, 0] | dIES | $\Psi_{N1\Leftrightarrow N2}(E) - \Psi_{N1\Leftrightarrow N2}(F)$ | $\alpha_2$ | 0.007 | 28.0 % |
| response | AN ⇔DMN | [-400, 0] | dIES | $\Psi_{N1\Leftrightarrow N2}(E) - \Psi_{N1\Leftrightarrow N2}(F)$ | $\beta_1$ | 4.10e-04 | 44.8 % |
| response | AN ⇔DMN | [-400, 0] | dIES | $\Psi_{N1\Leftrightarrow N2}(E) - \Psi_{N1\Leftrightarrow N2}(F)$ | $\gamma_1$ | 0.002 | 37.8 % |
| response | AN ⇔DMN | [-400, 0] | dIES | $\Psi_{N1\Leftrightarrow N2}(E) - \Psi_{N1\Leftrightarrow N2}(F)$ | $\gamma_2$ | 0.009 | 26.2 % |
| response | AN ⇔DMN | [-400, 0] | imp. hours | $\Psi_{N1\Leftrightarrow N2}(E) - \Psi_{N1\Leftrightarrow N2}(F)$ | $\alpha_1$ | 0.029 | 19.1 % |
| response | AN ⇔DMN | [-400, 0] | imp. hours | $\Psi_{N1\Leftrightarrow N2}(E) - \Psi_{N1\Leftrightarrow N2}(F)$ | $\alpha_2$ | 0.040 | 16.6 % |
| response | AN ⇔DMN | [-400, 0] | imp. hours | $\Psi_{N1\Leftrightarrow N2}(E) - \Psi_{N1\Leftrightarrow N2}(F)$ | $\alpha_3$ | 0.040 | 16.8 % |

Continued on next page

**Table 3 – continued from previous page**

| Locked to | Network | Time window | DV | IV | Frequency band | All musicians $p_{FDR}$ | $R^2$ |
|---|---|---|---|---|---|---|---|
| response | AN ⇔DMN | [-400, 0] | cl. hours | $\Psi_{N1\Leftrightarrow N2}(E) - \Psi_{N1\Leftrightarrow N2}(F)$ | $\beta_3$ | 0.036 | 17.7 % |
| response | AN ⇔DMN | [0, 400] | dIES | $\Psi_{N1\Leftrightarrow N2}(E) - \Psi_{N1\Leftrightarrow N2}(F)$ | $\alpha_1$ | 0.013 | 24.2 % |
| response | AN ⇔DMN | [0, 400] | dIES | $\Psi_{N1\Leftrightarrow N2}(E) - \Psi_{N1\Leftrightarrow N2}(F)$ | $\alpha_3$ | 0.003 | 35.1 % |
| response | AN ⇔DMN | [0, 400] | dIES | $\Psi_{N1\Leftrightarrow N2}(E) - \Psi_{N1\Leftrightarrow N2}(F)$ | $\beta_2$ | 0.013 | 23.9 % |
| response | AN ⇔DMN | [0, 400] | dIES | $\Psi_{N1\Leftrightarrow N2}(E) - \Psi_{N1\Leftrightarrow N2}(F)$ | $\beta_3$ | 0.005 | 32.2 % |
| response | AN ⇔DMN | [0, 400] | dIES | $\Psi_{N1\Leftrightarrow N2}(E) - \Psi_{N1\Leftrightarrow N2}(F)$ | $\gamma_3$ | 0.019 | 22.0 % |
| response | AN ⇔DMN | [0, 400] | imp. hours | $\Psi_{N1\Leftrightarrow N2}(E) - \Psi_{N1\Leftrightarrow N2}(F)$ | $\gamma_3$ | 0.046 | 15.1 % |
| response | CCN ⇔DMN | [-400, 0] | dIES | $\Psi_{N1\Leftrightarrow N2}(E) - \Psi_{N1\Leftrightarrow N2}(F)$ | $\delta$ | 0.027 | 19.7 % |
| response | CCN ⇔DMN | [-400, 0] | dIES | $\Psi_{N1\Leftrightarrow N2}(E) - \Psi_{N1\Leftrightarrow N2}(F)$ | $\alpha_1$ | 0.006 | 29.7 % |
| response | CCN ⇔DMN | [-400, 0] | dIES | $\Psi_{N1\Leftrightarrow N2}(E) - \Psi_{N1\Leftrightarrow N2}(F)$ | $\alpha_2$ | 0.006 | 29.4 % |
| response | CCN ⇔DMN | [-400, 0] | dIES | $\Psi_{N1\Leftrightarrow N2}(E) - \Psi_{N1\Leftrightarrow N2}(F)$ | $\beta_1$ | 6.38e-04 | 42.7 % |
| response | CCN ⇔DMN | [-400, 0] | dIES | $\Psi_{N1\Leftrightarrow N2}(E) - \Psi_{N1\Leftrightarrow N2}(F)$ | $\gamma_1$ | 0.006 | 29.3 % |
| response | CCN ⇔DMN | [-400, 0] | dIES | $\Psi_{N1\Leftrightarrow N2}(E) - \Psi_{N1\Leftrightarrow N2}(F)$ | $\gamma_2$ | 0.005 | 30.8 % |
| response | CCN ⇔DMN | [-400, 0] | imp. hours | $\Psi_{N1\Leftrightarrow N2}(E) - \Psi_{N1\Leftrightarrow N2}(F)$ | $\alpha_1$ | 0.024 | 20.4 % |
| response | CCN ⇔DMN | [-400, 0] | cl. hours | $\Psi_{N1\Leftrightarrow N2}(E) - \Psi_{N1\Leftrightarrow N2}(F)$ | $\beta_3$ | 0.048 | 14.7 % |
| response | CCN ⇔DMN | [0, 400] | dIES | $\Psi_{N1\Leftrightarrow N2}(E) - \Psi_{N1\Leftrightarrow N2}(F)$ | $\alpha_3$ | 0.005 | 31.5 % |
| response | CCN ⇔DMN | [0, 400] | dIES | $\Psi_{N1\Leftrightarrow N2}(E) - \Psi_{N1\Leftrightarrow N2}(F)$ | $\beta_2$ | 0.009 | 26.4 % |
| response | CCN ⇔DMN | [0, 400] | cl. hours | $\Psi_{N1\Leftrightarrow N2}(E) - \Psi_{N1\Leftrightarrow N2}(F)$ | $\gamma_3$ | 0.030 | 18.7 % |
| response | DMN ⇔SN | [-400, 0] | dIES | $\Psi_{N1\Leftrightarrow N2}(E) - \Psi_{N1\Leftrightarrow N2}(F)$ | $\alpha_2$ | 0.023 | 20.8 % |
| response | DMN ⇔SN | [-400, 0] | dIES | $\Psi_{N1\Leftrightarrow N2}(E) - \Psi_{N1\Leftrightarrow N2}(F)$ | $\beta_1$ | 0.025 | 20.3 % |
| response | DMN ⇔SN | [-400, 0] | dIES | $\Psi_{N1\Leftrightarrow N2}(E) - \Psi_{N1\Leftrightarrow N2}(F)$ | $\beta_2$ | 0.029 | 19.1 % |
| response | DMN ⇔SN | [-400, 0] | dIES | $\Psi_{N1\Leftrightarrow N2}(E) - \Psi_{N1\Leftrightarrow N2}(F)$ | $\gamma_1$ | 0.008 | 26.9 % |
| response | DMN ⇔SN | [-400, 0] | cl. hours | $\Psi_{N1\Leftrightarrow N2}(E) - \Psi_{N1\Leftrightarrow N2}(F)$ | $\delta$ | 0.045 | 15.5 % |
| response | DMN ⇔SN | [0, 400] | dIES | $\Psi_{N1\Leftrightarrow N2}(E) - \Psi_{N1\Leftrightarrow N2}(F)$ | $\beta_3$ | 0.006 | 30.2 % |
| response | DMN ⇔SN | [0, 400] | dIES | $\Psi_{N1\Leftrightarrow N2}(E) - \Psi_{N1\Leftrightarrow N2}(F)$ | $\gamma_3$ | 0.010 | 25.5 % |
| response | AN ⇔SN | [-400, 0] | dIES | $\Psi_{N1\Leftrightarrow N2}(E) - \Psi_{N1\Leftrightarrow N2}(F)$ | $\delta$ | 0.001 | 39.5 % |
| response | AN ⇔SN | [-400, 0] | dIES | $\Psi_{N1\Leftrightarrow N2}(E) - \Psi_{N1\Leftrightarrow N2}(F)$ | $\theta$ | 0.020 | 21.7 % |
| response | AN ⇔SN | [-400, 0] | dIES | $\Psi_{N1\Leftrightarrow N2}(E) - \Psi_{N1\Leftrightarrow N2}(F)$ | $\alpha_1$ | 0.033 | 18.2 % |

Continued on next page

**Table 3 – continued from previous page**

| Locked to | Network | Time window | DV | IV | Frequency band | All musicians $p_{FDR}$ | All musicians $R^2$ |
|---|---|---|---|---|---|---|---|
| response | AN ⇔SN | [-400, 0] | dIES | $\Psi_{N1\Leftrightarrow N2}(E) - \Psi_{N1\Leftrightarrow N2}(F)$ | $\alpha_2$ | 0.032 | 18.4 % |
| response | AN ⇔SN | [-400, 0] | dIES | $\Psi_{N1\Leftrightarrow N2}(E) - \Psi_{N1\Leftrightarrow N2}(F)$ | $\beta_1$ | 0.014 | 23.5 % |
| response | AN ⇔SN | [-400, 0] | dIES | $\Psi_{N1\Leftrightarrow N2}(E) - \Psi_{N1\Leftrightarrow N2}(F)$ | $\gamma_1$ | 0.006 | 29.3 % |
| response | AN ⇔SN | [0, 400] | dIES | $\Psi_{N1\Leftrightarrow N2}(E) - \Psi_{N1\Leftrightarrow N2}(F)$ | $\alpha_1$ | 0.008 | 27.3 % |
| response | AN ⇔SN | [0, 400] | dIES | $\Psi_{N1\Leftrightarrow N2}(E) - \Psi_{N1\Leftrightarrow N2}(F)$ | $\alpha_3$ | 0.006 | 29.2 % |
| response | AN ⇔SN | [0, 400] | dIES | $\Psi_{N1\Leftrightarrow N2}(E) - \Psi_{N1\Leftrightarrow N2}(F)$ | $\beta_2$ | 0.005 | 30.6 % |
| response | AN ⇔SN | [0, 400] | dIES | $\Psi_{N1\Leftrightarrow N2}(E) - \Psi_{N1\Leftrightarrow N2}(F)$ | $\beta_3$ | 0.021 | 21.4 % |
| response | AN ⇔SN | [0, 400] | dIES | $\Psi_{N1\Leftrightarrow N2}(E) - \Psi_{N1\Leftrightarrow N2}(F)$ | $\gamma_2$ | 0.015 | 23.1 % |
| response | AN ⇔SN | [0, 400] | dIES | $\Psi_{N1\Leftrightarrow N2}(E) - \Psi_{N1\Leftrightarrow N2}(F)$ | $\gamma_3$ | 0.010 | 25.8 % |
| response | AN ⇔SN | [0, 400] | imp. hours | $\Psi_{N1\Leftrightarrow N2}(E) - \Psi_{N1\Leftrightarrow N2}(F)$ | $\gamma_1$ | 0.048 | 14.7 % |
| response | CCN ⇔SN | [-400, 0] | dIES | $\Psi_{N1\Leftrightarrow N2}(E) - \Psi_{N1\Leftrightarrow N2}(F)$ | $\alpha_2$ | 0.004 | 34.1 % |
| response | CCN ⇔SN | [-400, 0] | dIES | $\Psi_{N1\Leftrightarrow N2}(E) - \Psi_{N1\Leftrightarrow N2}(F)$ | $\gamma_1$ | 0.002 | 36.8 % |
| response | CCN ⇔SN | [0, 400] | dIES | $\Psi_{N1\Leftrightarrow N2}(E) - \Psi_{N1\Leftrightarrow N2}(F)$ | $\beta_2$ | 0.006 | 30.0 % |
| response | CCN ⇔SN | [0, 400] | dIES | $\Psi_{N1\Leftrightarrow N2}(E) - \Psi_{N1\Leftrightarrow N2}(F)$ | $\gamma_3$ | 4.10e-04 | 45.3 % |
| chord | CCN ⇔FPN-L | [0, 400] | dIES | $\Psi_{N1\Leftrightarrow N2}(E) - \Psi_{N1\Leftrightarrow N2}(F)$ | $\beta_2$ | 0.022 | 21.1 % |
| chord | CCN ⇔FPN-L | [0, 400] | dIES | $\Psi_{N1\Leftrightarrow N2}(E) - \Psi_{N1\Leftrightarrow N2}(F)$ | $\beta_4$ | 0.006 | 34.8 % |
| chord | CCN ⇔FPN-R | [0, 400] | imp. hours | $\Psi_{N1\Leftrightarrow N2}(E) - \Psi_{N1\Leftrightarrow N2}(F)$ | $\alpha_3$ | 0.036 | 16.8 % |
| chord | CCN ⇔FPN-R | [0, 400] | cl. hours | $\Psi_{N1\Leftrightarrow N2}(E) - \Psi_{N1\Leftrightarrow N2}(F)$ | $\delta$ | 0.049 | 15.2 % |
| chord | CCN ⇔DAN | [0, 400] | dIES | $\Psi_{N1\Leftrightarrow N2}(E) - \Psi_{N1\Leftrightarrow N2}(F)$ | $\beta_4$ | 0.009 | 28.9 % |
| response | CCN ⇔FPN-L | [-400, 0] | dIES | $\Psi_{N1\Leftrightarrow N2}(E) - \Psi_{N1\Leftrightarrow N2}(F)$ | $\delta$ | 0.006 | 33.4 % |
| response | CCN ⇔FPN-L | [-400, 0] | dIES | $\Psi_{N1\Leftrightarrow N2}(E) - \Psi_{N1\Leftrightarrow N2}(F)$ | $\gamma_1$ | 0.013 | 24.6 % |
| response | CCN ⇔FPN-L | [-400, 0] | dIES | $\Psi_{N1\Leftrightarrow N2}(E) - \Psi_{N1\Leftrightarrow N2}(F)$ | $\gamma_3$ | 0.025 | 19.6 % |
| response | CCN ⇔FPN-L | [0, 400] | dIES | $\Psi_{N1\Leftrightarrow N2}(E) - \Psi_{N1\Leftrightarrow N2}(F)$ | $\beta_2$ | 0.008 | 30.3 % |
| response | CCN ⇔FPN-L | [0, 400] | dIES | $\Psi_{N1\Leftrightarrow N2}(E) - \Psi_{N1\Leftrightarrow N2}(F)$ | $\gamma_3$ | 0.017 | 22.5 % |
| response | CCN ⇔FPN-L | [0, 400] | imp. hours | $\Psi_{N1\Leftrightarrow N2}(E) - \Psi_{N1\Leftrightarrow N2}(F)$ | $\beta_4$ | 0.013 | 25.2 % |
| response | CCN ⇔FPN-R | [-400, 0] | dIES | $\Psi_{N1\Leftrightarrow N2}(E) - \Psi_{N1\Leftrightarrow N2}(F)$ | $\delta$ | 0.025 | 20.0 % |
| response | CCN ⇔FPN-R | [-400, 0] | dIES | $\Psi_{N1\Leftrightarrow N2}(E) - \Psi_{N1\Leftrightarrow N2}(F)$ | $\alpha_1$ | 0.017 | 22.7 % |
| response | CCN ⇔FPN-R | [-400, 0] | dIES | $\Psi_{N1\Leftrightarrow N2}(E) - \Psi_{N1\Leftrightarrow N2}(F)$ | $\alpha_2$ | 0.012 | 26.7 % |

Continued on next page

**Table 3 – continued from previous page**

| Locked to | Network | Time window | DV | IV | Frequency band | All musicians $p_{FDR}$ | $R^2$ |
|---|---|---|---|---|---|---|---|
| response | CCN ⇔FPN-R | [-400, 0] | dIES | $\Psi_{N1\Leftrightarrow N2}(E) - \Psi_{N1\Leftrightarrow N2}(F)$ | $\gamma_2$ | 0.012 | 26.3 % |
| response | CCN ⇔FPN-R | [-400, 0] | imp. hours | $\Psi_{N1\Leftrightarrow N2}(E) - \Psi_{N1\Leftrightarrow N2}(F)$ | $\alpha_2$ | 0.025 | 19.9 % |
| response | CCN ⇔FPN-R | [0, 400] | dIES | $\Psi_{N1\Leftrightarrow N2}(E) - \Psi_{N1\Leftrightarrow N2}(F)$ | $\alpha_3$ | 0.008 | 30.5 % |
| response | CCN ⇔FPN-R | [0, 400] | dIES | $\Psi_{N1\Leftrightarrow N2}(E) - \Psi_{N1\Leftrightarrow N2}(F)$ | $\beta_3$ | 8.02e-04 | 44.0 % |
| response | CCN ⇔FPN-R | [0, 400] | dIES | $\Psi_{N1\Leftrightarrow N2}(E) - \Psi_{N1\Leftrightarrow N2}(F)$ | $\gamma_3$ | 0.006 | 34.1 % |
| response | CCN ⇔FPN-R | [0, 400] | cl. hours | $\Psi_{N1\Leftrightarrow N2}(E) - \Psi_{N1\Leftrightarrow N2}(F)$ | $\gamma_3$ | 0.025 | 19.4 % |
| response | CCN ⇔DAN | [-400, 0] | dIES | $\Psi_{N1\Leftrightarrow N2}(E) - \Psi_{N1\Leftrightarrow N2}(F)$ | $\alpha_2$ | 0.025 | 19.6 % |
| response | CCN ⇔DAN | [-400, 0] | dIES | $\Psi_{N1\Leftrightarrow N2}(E) - \Psi_{N1\Leftrightarrow N2}(F)$ | $\beta_1$ | 0.013 | 25.4 % |
| response | CCN ⇔DAN | [-400, 0] | dIES | $\Psi_{N1\Leftrightarrow N2}(E) - \Psi_{N1\Leftrightarrow N2}(F)$ | $\gamma_1$ | 0.009 | 28.8 % |
| response | CCN ⇔DAN | [-400, 0] | dIES | $\Psi_{N1\Leftrightarrow N2}(E) - \Psi_{N1\Leftrightarrow N2}(F)$ | $\gamma_2$ | 0.015 | 23.6 % |
| response | CCN ⇔DAN | [-400, 0] | dIES | $\Psi_{N1\Leftrightarrow N2}(E) - \Psi_{N1\Leftrightarrow N2}(F)$ | $\gamma_3$ | 0.012 | 26.3 % |
| response | CCN ⇔DAN | [-400, 0] | cl. hours | $\Psi_{N1\Leftrightarrow N2}(E) - \Psi_{N1\Leftrightarrow N2}(F)$ | $\beta_4$ | 0.025 | 20.4 % |
| response | CCN ⇔DAN | [0, 400] | dIES | $\Psi_{N1\Leftrightarrow N2}(E) - \Psi_{N1\Leftrightarrow N2}(F)$ | $\alpha_1$ | 0.009 | 28.5 % |
| response | CCN ⇔DAN | [0, 400] | dIES | $\Psi_{N1\Leftrightarrow N2}(E) - \Psi_{N1\Leftrightarrow N2}(F)$ | $\alpha_2$ | 0.009 | 28.3 % |
| response | CCN ⇔DAN | [0, 400] | dIES | $\Psi_{N1\Leftrightarrow N2}(E) - \Psi_{N1\Leftrightarrow N2}(F)$ | $\beta_2$ | 0.006 | 32.6 % |
| response | CCN ⇔DAN | [0, 400] | dIES | $\Psi_{N1\Leftrightarrow N2}(E) - \Psi_{N1\Leftrightarrow N2}(F)$ | $\gamma_3$ | 0.013 | 25.5 % |
| response | CCN ⇔DAN | [0, 400] | cl. hours | $\Psi_{N1\Leftrightarrow N2}(E) - \Psi_{N1\Leftrightarrow N2}(F)$ | $\gamma_3$ | 0.046 | 15.5 % |
| chord | CCN ⇔DAN-core | [0, 400] | dIES | $\Psi_{N1\Leftrightarrow N2}(E) - \Psi_{N1\Leftrightarrow N2}(F)$ | $\beta_4$ | 0.021 | 22.5 % |
| chord | CCN ⇔VN | [0, 400] | dIES | $\Psi_{N1\Leftrightarrow N2}(E) - \Psi_{N1\Leftrightarrow N2}(F)$ | $\beta_3$ | 0.033 | 17.3 % |
| chord | CCN ⇔VN | [0, 400] | dIES | $\Psi_{N1\Leftrightarrow N2}(E) - \Psi_{N1\Leftrightarrow N2}(F)$ | $\gamma_2$ | 0.021 | 23.4 % |
| chord | CCN ⇔VN | [0, 400] | dIES | $\Psi_{N1\Leftrightarrow N2}(E) - \Psi_{N1\Leftrightarrow N2}(F)$ | $\gamma_3$ | 0.026 | 20.2 % |
| response | CCN ⇔DAN-core | [-400, 0] | dIES | $\Psi_{N1\Leftrightarrow N2}(E) - \Psi_{N1\Leftrightarrow N2}(F)$ | $\alpha_1$ | 0.031 | 18.0 % |
| response | CCN ⇔DAN-core | [-400, 0] | dIES | $\Psi_{N1\Leftrightarrow N2}(E) - \Psi_{N1\Leftrightarrow N2}(F)$ | $\alpha_2$ | 0.028 | 18.7 % |
| response | CCN ⇔DAN-core | [-400, 0] | dIES | $\Psi_{N1\Leftrightarrow N2}(E) - \Psi_{N1\Leftrightarrow N2}(F)$ | $\gamma_1$ | 0.018 | 25.6 % |
| response | CCN ⇔DAN-core | [-400, 0] | dIES | $\Psi_{N1\Leftrightarrow N2}(E) - \Psi_{N1\Leftrightarrow N2}(F)$ | $\gamma_2$ | 0.017 | 28.0 % |
| response | CCN ⇔DAN-core | [-400, 0] | dIES | $\Psi_{N1\Leftrightarrow N2}(E) - \Psi_{N1\Leftrightarrow N2}(F)$ | $\gamma_3$ | 0.021 | 21.3 % |
| response | CCN ⇔DAN-core | [-400, 0] | imp. hours | $\Psi_{N1\Leftrightarrow N2}(E) - \Psi_{N1\Leftrightarrow N2}(F)$ | $\alpha_3$ | 0.036 | 16.4 % |
| response | CCN ⇔DAN-core | [-400, 0] | cl. hours | $\Psi_{N1\Leftrightarrow N2}(E) - \Psi_{N1\Leftrightarrow N2}(F)$ | $\beta_4$ | 0.040 | 15.5 % |

Continued on next page

**Table 3 – continued from previous page**

| Locked to | Network | Time window | DV | IV | Frequency band | All musicians $p_{FDR}$ | $R^2$ |
|---|---|---|---|---|---|---|---|
| response | CCN ⇔DAN-core | [0, 400] | dIES | $\Psi_{N1\Leftrightarrow N2}(E) - \Psi_{N1\Leftrightarrow N2}(F)$ | $\alpha_3$ | 0.013 | 30.1 % |
| response | CCN ⇔DAN-core | [0, 400] | dIES | $\Psi_{N1\Leftrightarrow N2}(E) - \Psi_{N1\Leftrightarrow N2}(F)$ | $\beta_2$ | 0.013 | 32.9 % |
| response | CCN ⇔DAN-core | [0, 400] | dIES | $\Psi_{N1\Leftrightarrow N2}(E) - \Psi_{N1\Leftrightarrow N2}(F)$ | $\gamma_3$ | 0.021 | 21.4 % |
| response | CCN ⇔DAN-core | [0, 400] | imp. hours | $\Psi_{N1\Leftrightarrow N2}(E) - \Psi_{N1\Leftrightarrow N2}(F)$ | $\alpha_1$ | 0.028 | 19.1 % |
| response | CCN ⇔DAN-core | [0, 400] | cl. hours | $\Psi_{N1\Leftrightarrow N2}(E) - \Psi_{N1\Leftrightarrow N2}(F)$ | $\gamma_3$ | 0.021 | 21.8 % |
| response | CCN ⇔VN | [-400, 0] | dIES | $\Psi_{N1\Leftrightarrow N2}(E) - \Psi_{N1\Leftrightarrow N2}(F)$ | $\beta_1$ | 0.018 | 25.7 % |
| response | CCN ⇔VN | [-400, 0] | cl. hours | $\Psi_{N1\Leftrightarrow N2}(E) - \Psi_{N1\Leftrightarrow N2}(F)$ | $\beta_3$ | 0.048 | 14.6 % |
| response | CCN ⇔VN | [-400, 0] | cl. hours | $\Psi_{N1\Leftrightarrow N2}(E) - \Psi_{N1\Leftrightarrow N2}(F)$ | $\gamma_1$ | 0.021 | 23.6 % |
| response | CCN ⇔VN | [0, 400] | dIES | $\Psi_{N1\Leftrightarrow N2}(E) - \Psi_{N1\Leftrightarrow N2}(F)$ | $\alpha_2$ | 0.013 | 30.3 % |
| response | CCN ⇔VN | [0, 400] | dIES | $\Psi_{N1\Leftrightarrow N2}(E) - \Psi_{N1\Leftrightarrow N2}(F)$ | $\beta_2$ | 0.018 | 25.6 % |
| response | CCN ⇔VN | [0, 400] | imp. hours | $\Psi_{N1\Leftrightarrow N2}(E) - \Psi_{N1\Leftrightarrow N2}(F)$ | $\gamma_1$ | 0.033 | 17.1 % |
| response | CCN ⇔VN | [0, 400] | imp. hours | $\Psi_{N1\Leftrightarrow N2}(E) - \Psi_{N1\Leftrightarrow N2}(F)$ | $\gamma_3$ | 0.028 | 19.7 % |
| response | CCN ⇔VN | [0, 400] | cl. hours | $\Psi_{N1\Leftrightarrow N2}(E) - \Psi_{N1\Leftrightarrow N2}(F)$ | $\alpha_1$ | 0.028 | 18.6 % |
| chord | DMN ⇔FPN-L | [0, 400] | dIES | $\Psi_{N1\Leftrightarrow N2}(E) - \Psi_{N1\Leftrightarrow N2}(F)$ | $\beta_4$ | 7.65e-04 | 44.2 % |
| chord | DMN ⇔DAN | [0, 400] | dIES | $\Psi_{N1\Leftrightarrow N2}(E) - \Psi_{N1\Leftrightarrow N2}(F)$ | $\beta_4$ | 0.012 | 28.7 % |
| chord | DMN ⇔DAN | [0, 400] | dIES | $\Psi_{N1\Leftrightarrow N2}(E) - \Psi_{N1\Leftrightarrow N2}(F)$ | $\gamma_2$ | 0.032 | 18.9 % |
| response | DMN ⇔FPN-L | [-400, 0] | dIES | $\Psi_{N1\Leftrightarrow N2}(E) - \Psi_{N1\Leftrightarrow N2}(F)$ | $\alpha_2$ | 0.024 | 21.8 % |
| response | DMN ⇔FPN-L | [-400, 0] | dIES | $\Psi_{N1\Leftrightarrow N2}(E) - \Psi_{N1\Leftrightarrow N2}(F)$ | $\beta_2$ | 0.033 | 18.2 % |
| response | DMN ⇔FPN-L | [-400, 0] | dIES | $\Psi_{N1\Leftrightarrow N2}(E) - \Psi_{N1\Leftrightarrow N2}(F)$ | $\gamma_1$ | 0.032 | 18.6 % |
| response | DMN ⇔FPN-L | [-400, 0] | imp. hours | $\Psi_{N1\Leftrightarrow N2}(E) - \Psi_{N1\Leftrightarrow N2}(F)$ | $\alpha_1$ | 0.028 | 20.1 % |
| response | DMN ⇔FPN-L | [0, 400] | dIES | $\Psi_{N1\Leftrightarrow N2}(E) - \Psi_{N1\Leftrightarrow N2}(F)$ | $\beta_2$ | 0.019 | 24.9 % |
| response | DMN ⇔FPN-R | [-400, 0] | dIES | $\Psi_{N1\Leftrightarrow N2}(E) - \Psi_{N1\Leftrightarrow N2}(F)$ | $\alpha_2$ | 0.043 | 15.7 % |
| response | DMN ⇔FPN-R | [-400, 0] | dIES | $\Psi_{N1\Leftrightarrow N2}(E) - \Psi_{N1\Leftrightarrow N2}(F)$ | $\gamma_1$ | 0.023 | 23.6 % |
| response | DMN ⇔FPN-R | [0, 400] | dIES | $\Psi_{N1\Leftrightarrow N2}(E) - \Psi_{N1\Leftrightarrow N2}(F)$ | $\beta_1$ | 0.024 | 22.1 % |
| response | DMN ⇔FPN-R | [0, 400] | dIES | $\Psi_{N1\Leftrightarrow N2}(E) - \Psi_{N1\Leftrightarrow N2}(F)$ | $\beta_3$ | 0.003 | 36.7 % |
| response | DMN ⇔DAN | [-400, 0] | dIES | $\Psi_{N1\Leftrightarrow N2}(E) - \Psi_{N1\Leftrightarrow N2}(F)$ | $\alpha_1$ | 0.019 | 25.4 % |
| response | DMN ⇔DAN | [-400, 0] | dIES | $\Psi_{N1\Leftrightarrow N2}(E) - \Psi_{N1\Leftrightarrow N2}(F)$ | $\alpha_2$ | 0.024 | 21.3 % |
| response | DMN ⇔DAN | [-400, 0] | dIES | $\Psi_{N1\Leftrightarrow N2}(E) - \Psi_{N1\Leftrightarrow N2}(F)$ | $\gamma_1$ | 0.017 | 26.1 % |

Continued on next page

**Table 3 – continued from previous page**

| Locked to | Network | Time window | DV | IV | Frequency band | All musicians $p_{FDR}$ | $R^2$ |
|---|---|---|---|---|---|---|---|
| response | DMN ⇔DAN | [-400, 0] | dIES | $\Psi_{N1\Leftrightarrow N2}(E) - \Psi_{N1\Leftrightarrow N2}(F)$ | $\gamma_2$ | 0.012 | 28.6 % |
| response | DMN ⇔DAN | [-400, 0] | imp. hours | $\Psi_{N1\Leftrightarrow N2}(E) - \Psi_{N1\Leftrightarrow N2}(F)$ | $\alpha_1$ | 0.011 | 30.2 % |
| response | DMN ⇔DAN | [-400, 0] | imp. hours | $\Psi_{N1\Leftrightarrow N2}(E) - \Psi_{N1\Leftrightarrow N2}(F)$ | $\alpha_3$ | 0.024 | 21.9 % |
| response | DMN ⇔DAN | [-400, 0] | cl. hours | $\Psi_{N1\Leftrightarrow N2}(E) - \Psi_{N1\Leftrightarrow N2}(F)$ | $\beta_3$ | 0.024 | 22.6 % |
| response | DMN ⇔DAN | [0, 400] | dIES | $\Psi_{N1\Leftrightarrow N2}(E) - \Psi_{N1\Leftrightarrow N2}(F)$ | $\beta_2$ | 0.001 | 40.1 % |
| response | DMN ⇔DAN | [0, 400] | dIES | $\Psi_{N1\Leftrightarrow N2}(E) - \Psi_{N1\Leftrightarrow N2}(F)$ | $\beta_3$ | 0.006 | 33.1 % |
| response | DMN ⇔DAN | [0, 400] | imp. hours | $\Psi_{N1\Leftrightarrow N2}(E) - \Psi_{N1\Leftrightarrow N2}(F)$ | $\beta_4$ | 0.040 | 16.3 % |
| response | DMN ⇔DAN | [0, 400] | imp. hours | $\Psi_{N1\Leftrightarrow N2}(E) - \Psi_{N1\Leftrightarrow N2}(F)$ | $\gamma_3$ | 0.031 | 19.3 % |
| chord | DMN ⇔DAN-core | [0, 400] | dIES | $\Psi_{N1\Leftrightarrow N2}(E) - \Psi_{N1\Leftrightarrow N2}(F)$ | $\beta_4$ | 0.008 | 30.5 % |
| chord | DMN ⇔DAN-core | [0, 400] | dIES | $\Psi_{N1\Leftrightarrow N2}(E) - \Psi_{N1\Leftrightarrow N2}(F)$ | $\gamma_1$ | 0.045 | 16.0 % |
| chord | DMN ⇔VN | [0, 400] | dIES | $\Psi_{N1\Leftrightarrow N2}(E) - \Psi_{N1\Leftrightarrow N2}(F)$ | $\beta_3$ | 0.001 | 41.1 % |
| chord | DMN ⇔VN | [0, 400] | dIES | $\Psi_{N1\Leftrightarrow N2}(E) - \Psi_{N1\Leftrightarrow N2}(F)$ | $\gamma_1$ | 0.008 | 29.6 % |
| chord | DMN ⇔VN | [0, 400] | dIES | $\Psi_{N1\Leftrightarrow N2}(E) - \Psi_{N1\Leftrightarrow N2}(F)$ | $\gamma_2$ | 0.018 | 23.3 % |
| response | DMN ⇔DAN-core | [-400, 0] | dIES | $\Psi_{N1\Leftrightarrow N2}(E) - \Psi_{N1\Leftrightarrow N2}(F)$ | $\gamma_1$ | 0.016 | 25.0 % |
| response | DMN ⇔DAN-core | [-400, 0] | dIES | $\Psi_{N1\Leftrightarrow N2}(E) - \Psi_{N1\Leftrightarrow N2}(F)$ | $\gamma_2$ | 0.008 | 31.2 % |
| response | DMN ⇔DAN-core | [-400, 0] | imp. hours | $\Psi_{N1\Leftrightarrow N2}(E) - \Psi_{N1\Leftrightarrow N2}(F)$ | $\alpha_1$ | 0.027 | 21.2 % |
| response | DMN ⇔DAN-core | [-400, 0] | cl. hours | $\Psi_{N1\Leftrightarrow N2}(E) - \Psi_{N1\Leftrightarrow N2}(F)$ | $\beta_3$ | 0.042 | 17.3 % |
| response | DMN ⇔DAN-core | [0, 400] | dIES | $\Psi_{N1\Leftrightarrow N2}(E) - \Psi_{N1\Leftrightarrow N2}(F)$ | $\beta_2$ | 0.009 | 28.4 % |
| response | DMN ⇔DAN-core | [0, 400] | imp. hours | $\Psi_{N1\Leftrightarrow N2}(E) - \Psi_{N1\Leftrightarrow N2}(F)$ | $\beta_4$ | 0.033 | 20.0 % |
| response | DMN ⇔VN | [-400, 0] | dIES | $\Psi_{N1\Leftrightarrow N2}(E) - \Psi_{N1\Leftrightarrow N2}(F)$ | $\alpha_2$ | 0.020 | 22.6 % |
| response | DMN ⇔VN | [-400, 0] | dIES | $\Psi_{N1\Leftrightarrow N2}(E) - \Psi_{N1\Leftrightarrow N2}(F)$ | $\beta_2$ | 0.046 | 15.9 % |
| response | DMN ⇔VN | [-400, 0] | imp. hours | $\Psi_{N1\Leftrightarrow N2}(E) - \Psi_{N1\Leftrightarrow N2}(F)$ | $\alpha_2$ | 0.042 | 17.4 % |
| response | DMN ⇔VN | [-400, 0] | imp. hours | $\Psi_{N1\Leftrightarrow N2}(E) - \Psi_{N1\Leftrightarrow N2}(F)$ | $\alpha_3$ | 0.016 | 24.8 % |
| response | DMN ⇔VN | [0, 400] | dIES | $\Psi_{N1\Leftrightarrow N2}(E) - \Psi_{N1\Leftrightarrow N2}(F)$ | $\alpha_3$ | 0.018 | 24.0 % |
| response | DMN ⇔VN | [0, 400] | dIES | $\Psi_{N1\Leftrightarrow N2}(E) - \Psi_{N1\Leftrightarrow N2}(F)$ | $\beta_1$ | 0.016 | 24.9 % |
| response | DMN ⇔VN | [0, 400] | dIES | $\Psi_{N1\Leftrightarrow N2}(E) - \Psi_{N1\Leftrightarrow N2}(F)$ | $\beta_2$ | 0.008 | 29.4 % |
| response | DMN ⇔VN | [0, 400] | dIES | $\Psi_{N1\Leftrightarrow N2}(E) - \Psi_{N1\Leftrightarrow N2}(F)$ | $\beta_3$ | 0.005 | 35.0 % |
| response | DMN ⇔VN | [0, 400] | imp. hours | $\Psi_{N1\Leftrightarrow N2}(E) - \Psi_{N1\Leftrightarrow N2}(F)$ | $\gamma_3$ | 0.034 | 18.6 % |

Continued on next page

**Table 3 – continued from previous page**

| Locked to | Network | Time window | DV | IV | Frequency band | All musicians $p_{FDR}$ | $R^2$ |
|---|---|---|---|---|---|---|---|
| response | DMN ⇔VN | [0, 400] | cl. hours | $\Psi_{N1\Leftrightarrow N2}(E) - \Psi_{N1\Leftrightarrow N2}(F)$ | $\beta_3$ | 0.042 | 17.1 % |
| chord | SN ⇔FPN-L | [0, 400] | dIES | $\Psi_{N1\Leftrightarrow N2}(E) - \Psi_{N1\Leftrightarrow N2}(F)$ | $\beta_2$ | 0.020 | 23.2 % |
| chord | SN ⇔FPN-L | [0, 400] | dIES | $\Psi_{N1\Leftrightarrow N2}(E) - \Psi_{N1\Leftrightarrow N2}(F)$ | $\gamma_1$ | 0.032 | 19.4 % |
| chord | SN ⇔FPN-L | [0, 400] | cl. hours | $\Psi_{N1\Leftrightarrow N2}(E) - \Psi_{N1\Leftrightarrow N2}(F)$ | $\alpha_3$ | 0.047 | 15.6 % |
| chord | SN ⇔FPN-L | [0, 400] | cl. hours | $\Psi_{N1\Leftrightarrow N2}(E) - \Psi_{N1\Leftrightarrow N2}(F)$ | $\gamma_3$ | 0.042 | 17.3 % |
| chord | SN ⇔FPN-R | [0, 400] | dIES | $\Psi_{N1\Leftrightarrow N2}(E) - \Psi_{N1\Leftrightarrow N2}(F)$ | $\beta_3$ | 0.042 | 17.6 % |
| chord | SN ⇔FPN-R | [0, 400] | dIES | $\Psi_{N1\Leftrightarrow N2}(E) - \Psi_{N1\Leftrightarrow N2}(F)$ | $\beta_4$ | 0.012 | 29.5 % |
| chord | SN ⇔FPN-R | [0, 400] | cl. hours | $\Psi_{N1\Leftrightarrow N2}(E) - \Psi_{N1\Leftrightarrow N2}(F)$ | $\delta$ | 0.019 | 23.6 % |
| chord | SN ⇔DAN | [0, 400] | dIES | $\Psi_{N1\Leftrightarrow N2}(E) - \Psi_{N1\Leftrightarrow N2}(F)$ | $\beta_2$ | 0.018 | 24.7 % |
| response | SN ⇔FPN-L | [-400, 0] | dIES | $\Psi_{N1\Leftrightarrow N2}(E) - \Psi_{N1\Leftrightarrow N2}(F)$ | $\alpha_2$ | 0.023 | 22.2 % |
| response | SN ⇔FPN-L | [-400, 0] | dIES | $\Psi_{N1\Leftrightarrow N2}(E) - \Psi_{N1\Leftrightarrow N2}(F)$ | $\gamma_1$ | 0.032 | 20.4 % |
| response | SN ⇔FPN-L | [-400, 0] | dIES | $\Psi_{N1\Leftrightarrow N2}(E) - \Psi_{N1\Leftrightarrow N2}(F)$ | $\gamma_2$ | 0.019 | 23.9 % |
| response | SN ⇔FPN-L | [0, 400] | dIES | $\Psi_{N1\Leftrightarrow N2}(E) - \Psi_{N1\Leftrightarrow N2}(F)$ | $\beta_1$ | 0.012 | 28.2 % |
| response | SN ⇔FPN-L | [0, 400] | dIES | $\Psi_{N1\Leftrightarrow N2}(E) - \Psi_{N1\Leftrightarrow N2}(F)$ | $\beta_2$ | 0.012 | 28.3 % |
| response | SN ⇔FPN-R | [-400, 0] | dIES | $\Psi_{N1\Leftrightarrow N2}(E) - \Psi_{N1\Leftrightarrow N2}(F)$ | $\alpha_2$ | 0.016 | 25.6 % |
| response | SN ⇔FPN-R | [-400, 0] | dIES | $\Psi_{N1\Leftrightarrow N2}(E) - \Psi_{N1\Leftrightarrow N2}(F)$ | $\gamma_1$ | 0.014 | 27.1 % |
| response | SN ⇔FPN-R | [0, 400] | dIES | $\Psi_{N1\Leftrightarrow N2}(E) - \Psi_{N1\Leftrightarrow N2}(F)$ | $\alpha_2$ | 0.012 | 28.7 % |
| response | SN ⇔FPN-R | [0, 400] | dIES | $\Psi_{N1\Leftrightarrow N2}(E) - \Psi_{N1\Leftrightarrow N2}(F)$ | $\gamma_3$ | 0.011 | 31.2 % |
| response | SN ⇔DAN | [-400, 0] | dIES | $\Psi_{N1\Leftrightarrow N2}(E) - \Psi_{N1\Leftrightarrow N2}(F)$ | $\beta_2$ | 0.042 | 17.4 % |
| response | SN ⇔DAN | [-400, 0] | dIES | $\Psi_{N1\Leftrightarrow N2}(E) - \Psi_{N1\Leftrightarrow N2}(F)$ | $\gamma_1$ | 0.007 | 37.1 % |
| response | SN ⇔DAN | [0, 400] | dIES | $\Psi_{N1\Leftrightarrow N2}(E) - \Psi_{N1\Leftrightarrow N2}(F)$ | $\alpha_1$ | 0.018 | 24.6 % |
| response | SN ⇔DAN | [0, 400] | dIES | $\Psi_{N1\Leftrightarrow N2}(E) - \Psi_{N1\Leftrightarrow N2}(F)$ | $\beta_2$ | 0.010 | 33.7 % |
| response | SN ⇔DAN | [0, 400] | dIES | $\Psi_{N1\Leftrightarrow N2}(E) - \Psi_{N1\Leftrightarrow N2}(F)$ | $\beta_3$ | 0.023 | 22.2 % |
| response | SN ⇔DAN | [0, 400] | dIES | $\Psi_{N1\Leftrightarrow N2}(E) - \Psi_{N1\Leftrightarrow N2}(F)$ | $\beta_4$ | 0.032 | 19.6 % |
| response | SN ⇔DAN | [0, 400] | dIES | $\Psi_{N1\Leftrightarrow N2}(E) - \Psi_{N1\Leftrightarrow N2}(F)$ | $\gamma_3$ | 0.010 | 32.3 % |
| response | SN ⇔DAN | [0, 400] | imp. hours | $\Psi_{N1\Leftrightarrow N2}(E) - \Psi_{N1\Leftrightarrow N2}(F)$ | $\alpha_1$ | 0.041 | 17.8 % |
| response | SN ⇔DAN | [0, 400] | imp. hours | $\Psi_{N1\Leftrightarrow N2}(E) - \Psi_{N1\Leftrightarrow N2}(F)$ | $\alpha_2$ | 0.049 | 14.7 % |
| chord | DAN-core ⇔SN-core | [0, 400] | dIES | $\Psi_{N1\Leftrightarrow N2}(E) - \Psi_{N1\Leftrightarrow N2}(F)$ | $\beta_2$ | 0.022 | 24.1 % |

Continued on next page

**Table 3 – continued from previous page**

| Locked to | Network | Time window | DV | IV | Frequency band | All musicians $p_{FDR}$ | $R^2$ |
|---|---|---|---|---|---|---|---|
| chord | DAN-core ⇔SN-core | [0, 400] | dIES | $\Psi_{N1\Leftrightarrow N2}(E) - \Psi_{N1\Leftrightarrow N2}(F)$ | $\beta_4$ | 0.022 | 22.3 % |
| chord | DAN-core ⇔SN-core | [0, 400] | cl. hours | $\Psi_{N1\Leftrightarrow N2}(E) - \Psi_{N1\Leftrightarrow N2}(F)$ | $\gamma_3$ | 0.046 | 15.0 % |
| chord | DAN-core ⇔CON | [0, 400] | dIES | $\Psi_{N1\Leftrightarrow N2}(E) - \Psi_{N1\Leftrightarrow N2}(F)$ | $\beta_4$ | 0.023 | 21.8 % |
| chord | DAN-core ⇔CON | [0, 400] | imp. hours | $\Psi_{N1\Leftrightarrow N2}(E) - \Psi_{N1\Leftrightarrow N2}(F)$ | $\beta_1$ | 0.036 | 17.8 % |
| chord | DAN-core ⇔CON | [0, 400] | cl. hours | $\Psi_{N1\Leftrightarrow N2}(E) - \Psi_{N1\Leftrightarrow N2}(F)$ | $\gamma_3$ | 0.046 | 15.5 % |
| chord | VN ⇔SN-core | [0, 400] | dIES | $\Psi_{N1\Leftrightarrow N2}(E) - \Psi_{N1\Leftrightarrow N2}(F)$ | $\beta_2$ | 0.026 | 20.7 % |
| chord | VN ⇔SN-core | [0, 400] | imp. hours | $\Psi_{N1\Leftrightarrow N2}(E) - \Psi_{N1\Leftrightarrow N2}(F)$ | $\alpha_2$ | 0.044 | 16.3 % |
| chord | VN ⇔SN-core | [0, 400] | imp. hours | $\Psi_{N1\Leftrightarrow N2}(E) - \Psi_{N1\Leftrightarrow N2}(F)$ | $\beta_4$ | 0.017 | 25.5 % |
| chord | VN ⇔CON | [0, 400] | dIES | $\Psi_{N1\Leftrightarrow N2}(E) - \Psi_{N1\Leftrightarrow N2}(F)$ | $\beta_2$ | 0.030 | 19.9 % |
| chord | VN ⇔CON | [0, 400] | dIES | $\Psi_{N1\Leftrightarrow N2}(E) - \Psi_{N1\Leftrightarrow N2}(F)$ | $\gamma_3$ | 0.022 | 22.5 % |
| response | DAN-core ⇔SN-core | [-400, 0] | dIES | $\Psi_{N1\Leftrightarrow N2}(E) - \Psi_{N1\Leftrightarrow N2}(F)$ | $\alpha_2$ | 0.016 | 27.7 % |
| response | DAN-core ⇔SN-core | [-400, 0] | dIES | $\Psi_{N1\Leftrightarrow N2}(E) - \Psi_{N1\Leftrightarrow N2}(F)$ | $\gamma_1$ | 0.022 | 22.8 % |
| response | DAN-core ⇔SN-core | [0, 400] | dIES | $\Psi_{N1\Leftrightarrow N2}(E) - \Psi_{N1\Leftrightarrow N2}(F)$ | $\beta_2$ | 0.009 | 33.9 % |
| response | DAN-core ⇔SN-core | [0, 400] | dIES | $\Psi_{N1\Leftrightarrow N2}(E) - \Psi_{N1\Leftrightarrow N2}(F)$ | $\beta_3$ | 0.023 | 22.0 % |
| response | DAN-core ⇔SN-core | [0, 400] | dIES | $\Psi_{N1\Leftrightarrow N2}(E) - \Psi_{N1\Leftrightarrow N2}(F)$ | $\beta_4$ | 0.016 | 26.0 % |
| response | DAN-core ⇔SN-core | [0, 400] | dIES | $\Psi_{N1\Leftrightarrow N2}(E) - \Psi_{N1\Leftrightarrow N2}(F)$ | $\gamma_3$ | 0.016 | 26.7 % |
| response | DAN-core ⇔SN-core | [0, 400] | imp. hours | $\Psi_{N1\Leftrightarrow N2}(E) - \Psi_{N1\Leftrightarrow N2}(F)$ | $\beta_3$ | 0.032 | 19.0 % |
| response | DAN-core ⇔CON | [-400, 0] | dIES | $\Psi_{N1\Leftrightarrow N2}(E) - \Psi_{N1\Leftrightarrow N2}(F)$ | $\delta$ | 0.016 | 26.4 % |
| response | DAN-core ⇔CON | [-400, 0] | dIES | $\Psi_{N1\Leftrightarrow N2}(E) - \Psi_{N1\Leftrightarrow N2}(F)$ | $\alpha_2$ | 0.016 | 26.4 % |
| response | DAN-core ⇔CON | [-400, 0] | dIES | $\Psi_{N1\Leftrightarrow N2}(E) - \Psi_{N1\Leftrightarrow N2}(F)$ | $\beta_3$ | 0.030 | 20.0 % |
| response | DAN-core ⇔CON | [0, 400] | dIES | $\Psi_{N1\Leftrightarrow N2}(E) - \Psi_{N1\Leftrightarrow N2}(F)$ | $\beta_2$ | 0.003 | 41.2 % |
| response | DAN-core ⇔CON | [0, 400] | dIES | $\Psi_{N1\Leftrightarrow N2}(E) - \Psi_{N1\Leftrightarrow N2}(F)$ | $\beta_4$ | 0.022 | 23.3 % |
| response | DAN-core ⇔CON | [0, 400] | dIES | $\Psi_{N1\Leftrightarrow N2}(E) - \Psi_{N1\Leftrightarrow N2}(F)$ | $\gamma_3$ | 0.009 | 34.6 % |
| response | DAN-core ⇔CON | [0, 400] | imp. hours | $\Psi_{N1\Leftrightarrow N2}(E) - \Psi_{N1\Leftrightarrow N2}(F)$ | $\beta_4$ | 0.022 | 22.7 % |
| response | DAN-core ⇔CON | [0, 400] | cl. hours | $\Psi_{N1\Leftrightarrow N2}(E) - \Psi_{N1\Leftrightarrow N2}(F)$ | $\gamma_3$ | 0.016 | 26.1 % |
| response | VN ⇔SN-core | [-400, 0] | dIES | $\Psi_{N1\Leftrightarrow N2}(E) - \Psi_{N1\Leftrightarrow N2}(F)$ | $\beta_2$ | 0.022 | 22.3 % |
| response | VN ⇔SN-core | [-400, 0] | dIES | $\Psi_{N1\Leftrightarrow N2}(E) - \Psi_{N1\Leftrightarrow N2}(F)$ | $\gamma_1$ | 0.022 | 23.2 % |
| response | VN ⇔SN-core | [0, 400] | dIES | $\Psi_{N1\Leftrightarrow N2}(E) - \Psi_{N1\Leftrightarrow N2}(F)$ | $\alpha_3$ | 0.015 | 31.0 % |

Continued on next page

**Table 3 – continued from previous page**

| Locked to | Network | Time window | DV | IV | Frequency band | All musicians $p_{FDR}$ | $R^2$ |
|---|---|---|---|---|---|---|---|
| response | VN ⇔SN-core | [0, 400] | dIES | $\Psi_{N1\Leftrightarrow N2}(E) - \Psi_{N1\Leftrightarrow N2}(F)$ | $\beta_2$ | 0.030 | 19.8 % |
| response | VN ⇔SN-core | [0, 400] | dIES | $\Psi_{N1\Leftrightarrow N2}(E) - \Psi_{N1\Leftrightarrow N2}(F)$ | $\beta_3$ | 0.023 | 21.6 % |
| response | VN ⇔SN-core | [0, 400] | dIES | $\Psi_{N1\Leftrightarrow N2}(E) - \Psi_{N1\Leftrightarrow N2}(F)$ | $\gamma_1$ | 0.032 | 18.7 % |
| response | VN ⇔SN-core | [0, 400] | dIES | $\Psi_{N1\Leftrightarrow N2}(E) - \Psi_{N1\Leftrightarrow N2}(F)$ | $\gamma_3$ | 0.022 | 22.3 % |
| response | VN ⇔SN-core | [0, 400] | imp. hours | $\Psi_{N1\Leftrightarrow N2}(E) - \Psi_{N1\Leftrightarrow N2}(F)$ | $\beta_2$ | 0.034 | 18.4 % |
| response | VN ⇔SN-core | [0, 400] | imp. hours | $\Psi_{N1\Leftrightarrow N2}(E) - \Psi_{N1\Leftrightarrow N2}(F)$ | $\gamma_1$ | 0.016 | 26.9 % |
| response | VN ⇔SN-core | [0, 400] | cl. hours | $\Psi_{N1\Leftrightarrow N2}(E) - \Psi_{N1\Leftrightarrow N2}(F)$ | $\beta_1$ | 0.050 | 14.5 % |
| response | VN ⇔CON | [-400, 0] | dIES | $\Psi_{N1\Leftrightarrow N2}(E) - \Psi_{N1\Leftrightarrow N2}(F)$ | $\beta_2$ | 0.016 | 26.2 % |
| response | VN ⇔CON | [-400, 0] | dIES | $\Psi_{N1\Leftrightarrow N2}(E) - \Psi_{N1\Leftrightarrow N2}(F)$ | $\gamma_1$ | 0.032 | 19.3 % |
| response | VN ⇔CON | [-400, 0] | cl. hours | $\Psi_{N1\Leftrightarrow N2}(E) - \Psi_{N1\Leftrightarrow N2}(F)$ | $\gamma_1$ | 0.046 | 15.1 % |
| response | VN ⇔CON | [0, 400] | dIES | $\Psi_{N1\Leftrightarrow N2}(E) - \Psi_{N1\Leftrightarrow N2}(F)$ | $\alpha_2$ | 0.016 | 26.6 % |
| response | VN ⇔CON | [0, 400] | dIES | $\Psi_{N1\Leftrightarrow N2}(E) - \Psi_{N1\Leftrightarrow N2}(F)$ | $\beta_2$ | 0.016 | 28.5 % |
| response | VN ⇔CON | [0, 400] | dIES | $\Psi_{N1\Leftrightarrow N2}(E) - \Psi_{N1\Leftrightarrow N2}(F)$ | $\beta_3$ | 0.026 | 20.7 % |
| response | VN ⇔CON | [0, 400] | dIES | $\Psi_{N1\Leftrightarrow N2}(E) - \Psi_{N1\Leftrightarrow N2}(F)$ | $\gamma_1$ | 0.016 | 28.5 % |
| response | VN ⇔CON | [0, 400] | imp. hours | $\Psi_{N1\Leftrightarrow N2}(E) - \Psi_{N1\Leftrightarrow N2}(F)$ | $\beta_3$ | 0.034 | 18.1 % |
| response | VN ⇔CON | [0, 400] | imp. hours | $\Psi_{N1\Leftrightarrow N2}(E) - \Psi_{N1\Leftrightarrow N2}(F)$ | $\gamma_1$ | 0.032 | 18.7 % |
| chord | VN ⇔DAN-core | [0, 400] | dIES | $\Psi_{N1\Leftrightarrow N2}(E) - \Psi_{N1\Leftrightarrow N2}(F)$ | $\delta$ | 0.027 | 21.9 % |
| chord | VN ⇔DAN-core | [0, 400] | dIES | $\Psi_{N1\Leftrightarrow N2}(E) - \Psi_{N1\Leftrightarrow N2}(F)$ | $\beta_3$ | 0.012 | 27.9 % |
| chord | VN ⇔DAN-core | [0, 400] | cl. hours | $\Psi_{N1\Leftrightarrow N2}(E) - \Psi_{N1\Leftrightarrow N2}(F)$ | $\beta_4$ | 0.031 | 20.1 % |
| chord | VN ⇔FPN-L | [0, 400] | dIES | $\Psi_{N1\Leftrightarrow N2}(E) - \Psi_{N1\Leftrightarrow N2}(F)$ | $\theta$ | 0.023 | 23.3 % |
| chord | VN ⇔FPN-L | [0, 400] | imp. hours | $\Psi_{N1\Leftrightarrow N2}(E) - \Psi_{N1\Leftrightarrow N2}(F)$ | $\gamma_1$ | 0.036 | 18.6 % |
| chord | VN ⇔FPN-R | [0, 400] | dIES | $\Psi_{N1\Leftrightarrow N2}(E) - \Psi_{N1\Leftrightarrow N2}(F)$ | $\theta$ | 0.012 | 28.5 % |
| chord | DAN-core ⇔FPN-L | [0, 400] | dIES | $\Psi_{N1\Leftrightarrow N2}(E) - \Psi_{N1\Leftrightarrow N2}(F)$ | $\beta_2$ | 0.031 | 20.3 % |
| chord | DAN-core ⇔FPN-L | [0, 400] | dIES | $\Psi_{N1\Leftrightarrow N2}(E) - \Psi_{N1\Leftrightarrow N2}(F)$ | $\beta_4$ | 0.003 | 39.8 % |
| chord | DAN-core ⇔FPN-R | [0, 400] | imp. hours | $\Psi_{N1\Leftrightarrow N2}(E) - \Psi_{N1\Leftrightarrow N2}(F)$ | $\alpha_3$ | 0.018 | 24.7 % |
| response | VN ⇔DAN-core | [-400, 0] | dIES | $\Psi_{N1\Leftrightarrow N2}(E) - \Psi_{N1\Leftrightarrow N2}(F)$ | $\gamma_3$ | 0.031 | 20.5 % |
| response | VN ⇔DAN-core | [-400, 0] | cl. hours | $\Psi_{N1\Leftrightarrow N2}(E) - \Psi_{N1\Leftrightarrow N2}(F)$ | $\beta_4$ | 0.044 | 16.4 % |
| response | VN ⇔DAN-core | [-400, 0] | cl. hours | $\Psi_{N1\Leftrightarrow N2}(E) - \Psi_{N1\Leftrightarrow N2}(F)$ | $\gamma_1$ | 0.043 | 17.6 % |

Continued on next page

**Table 3 – continued from previous page**

| Locked to | Network | Time window | DV | IV | Frequency band | All musicians $p_{FDR}$ | $R^2$ |
|---|---|---|---|---|---|---|---|
| response | VN ⇔DAN-core | [0, 400] | dIES | $\Psi_{N1\Leftrightarrow N2}(E) - \Psi_{N1\Leftrightarrow N2}(F)$ | $\alpha_1$ | 0.014 | 26.2 % |
| response | VN ⇔DAN-core | [0, 400] | dIES | $\Psi_{N1\Leftrightarrow N2}(E) - \Psi_{N1\Leftrightarrow N2}(F)$ | $\alpha_2$ | 0.012 | 27.9 % |
| response | VN ⇔DAN-core | [0, 400] | dIES | $\Psi_{N1\Leftrightarrow N2}(E) - \Psi_{N1\Leftrightarrow N2}(F)$ | $\alpha_3$ | 0.012 | 29.9 % |
| response | VN ⇔DAN-core | [0, 400] | dIES | $\Psi_{N1\Leftrightarrow N2}(E) - \Psi_{N1\Leftrightarrow N2}(F)$ | $\beta_1$ | 0.014 | 26.3 % |
| response | VN ⇔DAN-core | [0, 400] | imp. hours | $\Psi_{N1\Leftrightarrow N2}(E) - \Psi_{N1\Leftrightarrow N2}(F)$ | $\alpha_3$ | 0.033 | 19.7 % |
| response | VN ⇔DAN-core | [0, 400] | cl. hours | $\Psi_{N1\Leftrightarrow N2}(E) - \Psi_{N1\Leftrightarrow N2}(F)$ | $\beta_3$ | 0.043 | 17.4 % |
| response | VN ⇔FPN-L | [-400, 0] | dIES | $\Psi_{N1\Leftrightarrow N2}(E) - \Psi_{N1\Leftrightarrow N2}(F)$ | $\theta$ | 0.019 | 24.3 % |
| response | VN ⇔FPN-L | [-400, 0] | dIES | $\Psi_{N1\Leftrightarrow N2}(E) - \Psi_{N1\Leftrightarrow N2}(F)$ | $\beta_3$ | 0.012 | 28.3 % |
| response | VN ⇔FPN-L | [-400, 0] | dIES | $\Psi_{N1\Leftrightarrow N2}(E) - \Psi_{N1\Leftrightarrow N2}(F)$ | $\gamma_1$ | 0.030 | 21.0 % |
| response | VN ⇔FPN-L | [-400, 0] | dIES | $\Psi_{N1\Leftrightarrow N2}(E) - \Psi_{N1\Leftrightarrow N2}(F)$ | $\gamma_2$ | 0.034 | 19.2 % |
| response | VN ⇔FPN-L | [-400, 0] | dIES | $\Psi_{N1\Leftrightarrow N2}(E) - \Psi_{N1\Leftrightarrow N2}(F)$ | $\gamma_3$ | 0.031 | 20.4 % |
| response | VN ⇔FPN-L | [-400, 0] | cl. hours | $\Psi_{N1\Leftrightarrow N2}(E) - \Psi_{N1\Leftrightarrow N2}(F)$ | $\gamma_1$ | 0.043 | 17.5 % |
| response | VN ⇔FPN-L | [0, 400] | dIES | $\Psi_{N1\Leftrightarrow N2}(E) - \Psi_{N1\Leftrightarrow N2}(F)$ | $\delta$ | 0.012 | 30.0 % |
| response | VN ⇔FPN-L | [0, 400] | dIES | $\Psi_{N1\Leftrightarrow N2}(E) - \Psi_{N1\Leftrightarrow N2}(F)$ | $\beta_2$ | 0.012 | 31.2 % |
| response | VN ⇔FPN-L | [0, 400] | imp. hours | $\Psi_{N1\Leftrightarrow N2}(E) - \Psi_{N1\Leftrightarrow N2}(F)$ | $\gamma_3$ | 0.044 | 16.7 % |
| response | VN ⇔FPN-R | [-400, 0] | dIES | $\Psi_{N1\Leftrightarrow N2}(E) - \Psi_{N1\Leftrightarrow N2}(F)$ | $\beta_2$ | 0.004 | 37.5 % |
| response | VN ⇔FPN-R | [0, 400] | dIES | $\Psi_{N1\Leftrightarrow N2}(E) - \Psi_{N1\Leftrightarrow N2}(F)$ | $\beta_1$ | 0.024 | 23.0 % |
| response | VN ⇔FPN-R | [0, 400] | dIES | $\Psi_{N1\Leftrightarrow N2}(E) - \Psi_{N1\Leftrightarrow N2}(F)$ | $\beta_2$ | 0.029 | 21.3 % |
| response | VN ⇔FPN-R | [0, 400] | dIES | $\Psi_{N1\Leftrightarrow N2}(E) - \Psi_{N1\Leftrightarrow N2}(F)$ | $\beta_3$ | 0.015 | 25.7 % |
| response | VN ⇔FPN-R | [0, 400] | dIES | $\Psi_{N1\Leftrightarrow N2}(E) - \Psi_{N1\Leftrightarrow N2}(F)$ | $\gamma_1$ | 0.033 | 19.5 % |
| response | VN ⇔FPN-R | [0, 400] | imp. hours | $\Psi_{N1\Leftrightarrow N2}(E) - \Psi_{N1\Leftrightarrow N2}(F)$ | $\gamma_1$ | 0.044 | 16.5 % |
| response | VN ⇔FPN-R | [0, 400] | cl. hours | $\Psi_{N1\Leftrightarrow N2}(E) - \Psi_{N1\Leftrightarrow N2}(F)$ | $\beta_1$ | 0.026 | 22.5 % |
| response | DAN-core ⇔FPN-L | [-400, 0] | imp. hours | $\Psi_{N1\Leftrightarrow N2}(E) - \Psi_{N1\Leftrightarrow N2}(F)$ | $\beta_2$ | 0.027 | 22.0 % |
| response | DAN-core ⇔FPN-L | [0, 400] | dIES | $\Psi_{N1\Leftrightarrow N2}(E) - \Psi_{N1\Leftrightarrow N2}(F)$ | $\beta_2$ | 0.012 | 28.9 % |
| response | DAN-core ⇔FPN-L | [0, 400] | dIES | $\Psi_{N1\Leftrightarrow N2}(E) - \Psi_{N1\Leftrightarrow N2}(F)$ | $\beta_4$ | 0.012 | 28.6 % |
| response | DAN-core ⇔FPN-L | [0, 400] | dIES | $\Psi_{N1\Leftrightarrow N2}(E) - \Psi_{N1\Leftrightarrow N2}(F)$ | $\gamma_3$ | 0.020 | 24.0 % |
| response | DAN-core ⇔FPN-R | [-400, 0] | dIES | $\Psi_{N1\Leftrightarrow N2}(E) - \Psi_{N1\Leftrightarrow N2}(F)$ | $\gamma_1$ | 0.027 | 22.3 % |
| response | DAN-core ⇔FPN-R | [-400, 0] | dIES | $\Psi_{N1\Leftrightarrow N2}(E) - \Psi_{N1\Leftrightarrow N2}(F)$ | $\gamma_2$ | 0.014 | 26.8 % |

Continued on next page

**Table 3 – continued from previous page**

| Locked to | Network | Time window | DV | IV | Frequency band | All musicians $p_{FDR}$ | $R^2$ |
|---|---|---|---|---|---|---|---|
| response | DAN-core ⇔FPN-R | [0, 400] | dIES | $\Psi_{N1\Leftrightarrow N2}(E) - \Psi_{N1\Leftrightarrow N2}(F)$ | $\theta$ | 0.012 | 30.8 % |
| response | DAN-core ⇔FPN-R | [0, 400] | dIES | $\Psi_{N1\Leftrightarrow N2}(E) - \Psi_{N1\Leftrightarrow N2}(F)$ | $\alpha_3$ | 0.014 | 26.7 % |
| response | DAN-core ⇔FPN-R | [0, 400] | dIES | $\Psi_{N1\Leftrightarrow N2}(E) - \Psi_{N1\Leftrightarrow N2}(F)$ | $\beta_1$ | 0.039 | 18.2 % |
| response | DAN-core ⇔FPN-R | [0, 400] | dIES | $\Psi_{N1\Leftrightarrow N2}(E) - \Psi_{N1\Leftrightarrow N2}(F)$ | $\beta_2$ | 0.012 | 27.8 % |
| response | DAN-core ⇔FPN-R | [0, 400] | dIES | $\Psi_{N1\Leftrightarrow N2}(E) - \Psi_{N1\Leftrightarrow N2}(F)$ | $\beta_3$ | 0.014 | 26.4 % |
| response | DAN-core ⇔FPN-R | [0, 400] | dIES | $\Psi_{N1\Leftrightarrow N2}(E) - \Psi_{N1\Leftrightarrow N2}(F)$ | $\gamma_3$ | 0.011 | 33.0 % |
| response | DAN-core ⇔FPN-R | [0, 400] | imp. hours | $\Psi_{N1\Leftrightarrow N2}(E) - \Psi_{N1\Leftrightarrow N2}(F)$ | $\beta_3$ | 1.59e-04 | 50.0 % |
| chord | DMN ⇔SN-core | [0, 400] | dIES | $\Psi_{N1\Leftrightarrow N2}(E) - \Psi_{N1\Leftrightarrow N2}(F)$ | $\beta_2$ | 0.018 | 24.7 % |
| chord | DMN ⇔CON | [0, 400] | dIES | $\Psi_{N1\Leftrightarrow N2}(E) - \Psi_{N1\Leftrightarrow N2}(F)$ | $\beta_2$ | 0.029 | 21.7 % |
| chord | DMN ⇔CON | [0, 400] | dIES | $\Psi_{N1\Leftrightarrow N2}(E) - \Psi_{N1\Leftrightarrow N2}(F)$ | $\beta_4$ | 0.016 | 26.9 % |
| response | DMN ⇔SN-core | [-400, 0] | dIES | $\Psi_{N1\Leftrightarrow N2}(E) - \Psi_{N1\Leftrightarrow N2}(F)$ | $\delta$ | 0.010 | 34.9 % |
| response | DMN ⇔SN-core | [-400, 0] | dIES | $\Psi_{N1\Leftrightarrow N2}(E) - \Psi_{N1\Leftrightarrow N2}(F)$ | $\gamma_1$ | 0.016 | 26.2 % |
| response | DMN ⇔SN-core | [0, 400] | dIES | $\Psi_{N1\Leftrightarrow N2}(E) - \Psi_{N1\Leftrightarrow N2}(F)$ | $\beta_1$ | 0.029 | 21.5 % |
| response | DMN ⇔SN-core | [0, 400] | dIES | $\Psi_{N1\Leftrightarrow N2}(E) - \Psi_{N1\Leftrightarrow N2}(F)$ | $\beta_3$ | 0.016 | 28.5 % |
| response | DMN ⇔CON | [-400, 0] | dIES | $\Psi_{N1\Leftrightarrow N2}(E) - \Psi_{N1\Leftrightarrow N2}(F)$ | $\beta_1$ | 0.016 | 26.6 % |
| response | DMN ⇔CON | [0, 400] | dIES | $\Psi_{N1\Leftrightarrow N2}(E) - \Psi_{N1\Leftrightarrow N2}(F)$ | $\beta_2$ | 0.016 | 25.7 % |
| response | DMN ⇔CON | [0, 400] | dIES | $\Psi_{N1\Leftrightarrow N2}(E) - \Psi_{N1\Leftrightarrow N2}(F)$ | $\gamma_3$ | 0.016 | 26.6 % |

Table 4: Details for significant findings within networks on the level of factors (corresponds to scatter plots). DV indicates the dependent variable, IV the independent variables, where an expression within square brackets represents one factor.

| Locked to | Network | Time window | DV | IV | Frequency band | All musicians p | All musicians $R^2$ | Improvisers p | Improvisers $R^2$ | Classical performers p | Classical performers $R^2$ |
|---|---|---|---|---|---|---|---|---|---|---|---|
| chord | AN ⇔CCN | [0, 400] | dIES | $\Psi_{N1\Leftrightarrow N2}(E) - \Psi_{N1\Leftrightarrow N2}(F)$ | $\beta_4$ | 9.99e-05 | 35.0 % | 0.055 | 21.9 % | 0.055 | 24.1 % |
| chord | AN ⇔CCN | [0, 400] | dIES | $\Psi_{N1\Leftrightarrow N2}(E) - \Psi_{N1\Leftrightarrow N2}(F)$ | $\gamma_1$ | 0.006 | 22.8 % | 0.030 | 23.7 % | 0.405 | 5.8 % |
| chord | AN ⇔CCN | [0, 400] | imp. hours | $\Psi_{N1\Leftrightarrow N2}(E) - \Psi_{N1\Leftrightarrow N2}(F)$ | $\beta_1$ | 0.035 | 23.4 % | 0.409 | 4.5 % | 1.000 | 40.7 % |
| chord | AN ⇔DMN | [0, 400] | dIES | $\Psi_{N1\Leftrightarrow N2}(E) - \Psi_{N1\Leftrightarrow N2}(F)$ | $\beta_4$ | 3.07e-04 | 32.3 % | 0.861 | 1.8 % | 0.120 | 17.3 % |
| chord | CCN ⇔DMN | [0, 400] | dIES | $\Psi_{N1\Leftrightarrow N2}(E) - \Psi_{N1\Leftrightarrow N2}(F)$ | $\beta_4$ | 1.66e-04 | 33.3 % | 0.985 | 1.0 % | 0.015 | 35.4 % |
| chord | DMN ⇔SN | [0, 400] | dIES | $\Psi_{N1\Leftrightarrow N2}(E) - \Psi_{N1\Leftrightarrow N2}(F)$ | $\beta_2$ | 0.001 | 33.4 % | 1.02e-04 | 57.8 % | 0.990 | 0.6 % |
| chord | AN ⇔SN | [0, 400] | dIES | $\Psi_{N1\Leftrightarrow N2}(E) - \Psi_{N1\Leftrightarrow N2}(F)$ | $\beta_2$ | 7.56e-06 | 46.8 % | 1.85e-04 | 55.4 % | 0.124 | 16.9 % |
| chord | CCN ⇔SN | [0, 400] | dIES | $\Psi_{N1\Leftrightarrow N2}(E) - \Psi_{N1\Leftrightarrow N2}(F)$ | $\beta_4$ | 3.56e-05 | 37.6 % | 0.148 | 13.2 % | 2.30e-04 | 64.4 % |
| chord | CCN ⇔SN | [0, 400] | dIES | $\Psi_{N1\Leftrightarrow N2}(E) - \Psi_{N1\Leftrightarrow N2}(F)$ | $\gamma_1$ | 0.015 | 21.0 % | 0.125 | 12.8 % | 0.223 | 10.8 % |
| response | AN ⇔CCN | [-400, 0] | dIES | $\Psi_{N1\Leftrightarrow N2}(E) - \Psi_{N1\Leftrightarrow N2}(F)$ | $\alpha_1$ | 0.002 | 29.3 % | 0.353 | 8.3 % | 0.020 | 33.0 % |
| response | AN ⇔CCN | [-400, 0] | dIES | $\Psi_{N1\Leftrightarrow N2}(E) - \Psi_{N1\Leftrightarrow N2}(F)$ | $\alpha_2$ | 0.006 | 24.6 % | 0.092 | 15.9 % | 0.045 | 25.8 % |
| response | AN ⇔CCN | [-400, 0] | dIES | $\Psi_{N1\Leftrightarrow N2}(E) - \Psi_{N1\Leftrightarrow N2}(F)$ | $\beta_1$ | 0.020 | 30.0 % | 0.155 | 14.8 % | 0.255 | 10.6 % |
| response | AN ⇔CCN | [-400, 0] | dIES | $\Psi_{N1\Leftrightarrow N2}(E) - \Psi_{N1\Leftrightarrow N2}(F)$ | $\gamma_1$ | 0.003 | 29.4 % | 0.012 | 31.0 % | 0.045 | 25.9 % |
| response | AN ⇔CCN | [-400, 0] | dIES | $\Psi_{N1\Leftrightarrow N2}(E) - \Psi_{N1\Leftrightarrow N2}(F)$ | $\gamma_3$ | 0.046 | 19.5 % | 0.371 | 5.9 % | 0.072 | 21.3 % |
| response | AN ⇔CCN | [0, 400] | dIES | $\Psi_{N1\Leftrightarrow N2}(E) - \Psi_{N1\Leftrightarrow N2}(F)$ | $\alpha_3$ | 0.046 | 30.3 % | 0.003 | 43.0 % | 0.974 | 0.5 % |
| response | AN ⇔CCN | [0, 400] | dIES | $\Psi_{N1\Leftrightarrow N2}(E) - \Psi_{N1\Leftrightarrow N2}(F)$ | $\beta_2$ | 0.018 | 33.8 % | 0.007 | 36.0 % | 0.504 | 3.5 % |
| response | AN ⇔CCN | [0, 400] | dIES | $\Psi_{N1\Leftrightarrow N2}(E) - \Psi_{N1\Leftrightarrow N2}(F)$ | $\beta_3$ | 0.037 | 25.8 % | 0.373 | 5.6 % | 0.045 | 25.7 % |
| response | AN ⇔CCN | [0, 400] | dIES | $\Psi_{N1\Leftrightarrow N2}(E) - \Psi_{N1\Leftrightarrow N2}(F)$ | $\gamma_3$ | 0.016 | 26.6 % | 0.074 | 17.4 % | 0.133 | 15.4 % |
| response | AN ⇔CCN | [0, 400] | cl. hours | $\Psi_{N1\Leftrightarrow N2}(E) - \Psi_{N1\Leftrightarrow N2}(F)$ | $\gamma_3$ | 0.008 | 18.0 % | 0.037 | 22.1 % | 0.944 | 0.0 % |
| response | AN ⇔DMN | [-400, 0] | dIES | $\Psi_{N1\Leftrightarrow N2}(E) - \Psi_{N1\Leftrightarrow N2}(F)$ | $\alpha_2$ | 0.004 | 29.9 % | 0.054 | 19.8 % | 0.021 | 32.4 % |
| response | AN ⇔DMN | [-400, 0] | dIES | $\Psi_{N1\Leftrightarrow N2}(E) - \Psi_{N1\Leftrightarrow N2}(F)$ | $\beta_1$ | 1.08e-05 | 46.3 % | 0.037 | 26.3 % | 0.047 | 25.7 % |
| response | AN ⇔DMN | [-400, 0] | dIES | $\Psi_{N1\Leftrightarrow N2}(E) - \Psi_{N1\Leftrightarrow N2}(F)$ | $\gamma_1$ | 1.06e-04 | 39.5 % | 0.039 | 22.6 % | 0.001 | 52.8 % |
| response | AN ⇔DMN | [-400, 0] | dIES | $\Psi_{N1\Leftrightarrow N2}(E) - \Psi_{N1\Leftrightarrow N2}(F)$ | $\gamma_2$ | 0.042 | 28.2 % | 0.454 | 5.0 % | 0.022 | 33.7 % |
| response | AN ⇔DMN | [-400, 0] | imp. hours | $\Psi_{N1\Leftrightarrow N2}(E) - \Psi_{N1\Leftrightarrow N2}(F)$ | $\alpha_1$ | 0.005 | 21.2 % | 0.126 | 16.4 % | 1.000 | 40.9 % |
| response | AN ⇔DMN | [-400, 0] | imp. hours | $\Psi_{N1\Leftrightarrow N2}(E) - \Psi_{N1\Leftrightarrow N2}(F)$ | $\alpha_2$ | 0.006 | 18.8 % | 0.221 | 10.7 % | 1.000 | 40.5 % |
| response | AN ⇔DMN | [-400, 0] | imp. hours | $\Psi_{N1\Leftrightarrow N2}(E) - \Psi_{N1\Leftrightarrow N2}(F)$ | $\alpha_3$ | 0.023 | 19.0 % | 0.849 | 4.3 % | 1.000 | 41.0 % |

Continued on next page

**Table 4 – continued from previous page**

| Locked to | Network | Time window | DV | IV | Frequency band | All musicians p | All musicians $R^2$ | Improvisers p | Improvisers $R^2$ | Classical performers p | Classical performers $R^2$ |
|---|---|---|---|---|---|---|---|---|---|---|---|
| response | AN ⇔DMN | [-400, 0] | cl. hours | $\Psi_{N1\Leftrightarrow N2}(E) - \Psi_{N1\Leftrightarrow N2}(F)$ | $\beta_3$ | 0.005 | 19.8 % | 0.029 | 24.5 % | 0.551 | 2.6 % |
| response | AN ⇔DMN | [0, 400] | dIES | $\Psi_{N1\Leftrightarrow N2}(E) - \Psi_{N1\Leftrightarrow N2}(F)$ | $\alpha_1$ | 0.034 | 26.2 % | 0.092 | 16.3 % | 0.196 | 12.7 % |
| response | AN ⇔DMN | [0, 400] | dIES | $\Psi_{N1\Leftrightarrow N2}(E) - \Psi_{N1\Leftrightarrow N2}(F)$ | $\alpha_3$ | 0.022 | 36.8 % | 0.025 | 28.1 % | 0.520 | 4.4 % |
| response | AN ⇔DMN | [0, 400] | dIES | $\Psi_{N1\Leftrightarrow N2}(E) - \Psi_{N1\Leftrightarrow N2}(F)$ | $\beta_2$ | 0.037 | 25.9 % | 0.029 | 25.2 % | 0.310 | 7.7 % |
| response | AN ⇔DMN | [0, 400] | dIES | $\Psi_{N1\Leftrightarrow N2}(E) - \Psi_{N1\Leftrightarrow N2}(F)$ | $\beta_3$ | 0.002 | 34.0 % | 0.233 | 8.1 % | 0.024 | 31.3 % |
| response | AN ⇔DMN | [0, 400] | dIES | $\Psi_{N1\Leftrightarrow N2}(E) - \Psi_{N1\Leftrightarrow N2}(F)$ | $\gamma_3$ | 0.035 | 24.0 % | 0.026 | 24.8 % | 0.750 | 1.1 % |
| response | AN ⇔DMN | [0, 400] | imp. hours | $\Psi_{N1\Leftrightarrow N2}(E) - \Psi_{N1\Leftrightarrow N2}(F)$ | $\gamma_3$ | 0.037 | 17.4 % | 0.106 | 15.3 % | 1.000 | 40.7 % |
| response | CCN ⇔DMN | [-400, 0] | dIES | $\Psi_{N1\Leftrightarrow N2}(E) - \Psi_{N1\Leftrightarrow N2}(F)$ | $\delta$ | 0.032 | 21.8 % | 0.271 | 7.6 % | 0.333 | 8.0 % |
| response | CCN ⇔DMN | [-400, 0] | dIES | $\Psi_{N1\Leftrightarrow N2}(E) - \Psi_{N1\Leftrightarrow N2}(F)$ | $\alpha_1$ | 0.003 | 31.6 % | 0.055 | 22.6 % | 0.042 | 26.3 % |
| response | CCN ⇔DMN | [-400, 0] | dIES | $\Psi_{N1\Leftrightarrow N2}(E) - \Psi_{N1\Leftrightarrow N2}(F)$ | $\alpha_2$ | 0.003 | 31.3 % | 0.026 | 25.5 % | 0.030 | 29.5 % |
| response | CCN ⇔DMN | [-400, 0] | dIES | $\Psi_{N1\Leftrightarrow N2}(E) - \Psi_{N1\Leftrightarrow N2}(F)$ | $\beta_1$ | 2.22e-05 | 44.2 % | 0.315 | 8.3 % | 0.010 | 39.6 % |
| response | CCN ⇔DMN | [-400, 0] | dIES | $\Psi_{N1\Leftrightarrow N2}(E) - \Psi_{N1\Leftrightarrow N2}(F)$ | $\gamma_1$ | 0.001 | 31.2 % | 0.002 | 42.4 % | 0.146 | 14.5 % |
| response | CCN ⇔DMN | [-400, 0] | dIES | $\Psi_{N1\Leftrightarrow N2}(E) - \Psi_{N1\Leftrightarrow N2}(F)$ | $\gamma_2$ | 0.018 | 32.6 % | 0.049 | 20.4 % | 0.016 | 38.6 % |
| response | CCN ⇔DMN | [-400, 0] | imp. hours | $\Psi_{N1\Leftrightarrow N2}(E) - \Psi_{N1\Leftrightarrow N2}(F)$ | $\alpha_1$ | 0.006 | 22.5 % | 0.244 | 15.2 % | 1.000 | 40.6 % |
| response | CCN ⇔DMN | [-400, 0] | cl. hours | $\Psi_{N1\Leftrightarrow N2}(E) - \Psi_{N1\Leftrightarrow N2}(F)$ | $\beta_3$ | 0.010 | 17.0 % | 0.111 | 14.5 % | 0.780 | 0.6 % |
| response | CCN ⇔DMN | [0, 400] | dIES | $\Psi_{N1\Leftrightarrow N2}(E) - \Psi_{N1\Leftrightarrow N2}(F)$ | $\alpha_3$ | 0.015 | 33.3 % | 0.018 | 31.7 % | 0.339 | 8.0 % |
| response | CCN ⇔DMN | [0, 400] | dIES | $\Psi_{N1\Leftrightarrow N2}(E) - \Psi_{N1\Leftrightarrow N2}(F)$ | $\beta_2$ | 0.050 | 28.4 % | 0.003 | 41.2 % | 0.893 | 0.6 % |
| response | CCN ⇔DMN | [0, 400] | cl. hours | $\Psi_{N1\Leftrightarrow N2}(E) - \Psi_{N1\Leftrightarrow N2}(F)$ | $\gamma_3$ | 0.004 | 20.8 % | 0.049 | 19.9 % | 0.190 | 12.0 % |
| response | DMN ⇔SN | [-400, 0] | dIES | $\Psi_{N1\Leftrightarrow N2}(E) - \Psi_{N1\Leftrightarrow N2}(F)$ | $\alpha_2$ | 0.038 | 22.9 % | 0.491 | 5.0 % | 0.043 | 26.1 % |
| response | DMN ⇔SN | [-400, 0] | dIES | $\Psi_{N1\Leftrightarrow N2}(E) - \Psi_{N1\Leftrightarrow N2}(F)$ | $\beta_1$ | 0.032 | 22.4 % | 0.709 | 2.9 % | 0.019 | 33.4 % |
| response | DMN ⇔SN | [-400, 0] | dIES | $\Psi_{N1\Leftrightarrow N2}(E) - \Psi_{N1\Leftrightarrow N2}(F)$ | $\beta_2$ | 0.006 | 21.2 % | 0.802 | 1.1 % | 0.033 | 29.3 % |
| response | DMN ⇔SN | [-400, 0] | dIES | $\Psi_{N1\Leftrightarrow N2}(E) - \Psi_{N1\Leftrightarrow N2}(F)$ | $\gamma_1$ | 0.004 | 28.9 % | 0.036 | 22.8 % | 0.059 | 23.1 % |
| response | DMN ⇔SN | [-400, 0] | cl. hours | $\Psi_{N1\Leftrightarrow N2}(E) - \Psi_{N1\Leftrightarrow N2}(F)$ | $\delta$ | 0.008 | 17.8 % | 4.47e-06 | 70.4 % | 0.456 | 4.1 % |
| response | DMN ⇔SN | [0, 400] | dIES | $\Psi_{N1\Leftrightarrow N2}(E) - \Psi_{N1\Leftrightarrow N2}(F)$ | $\beta_3$ | 0.006 | 32.0 % | 0.069 | 17.4 % | 0.312 | 7.8 % |
| response | DMN ⇔SN | [0, 400] | dIES | $\Psi_{N1\Leftrightarrow N2}(E) - \Psi_{N1\Leftrightarrow N2}(F)$ | $\gamma_3$ | 0.017 | 27.5 % | 0.086 | 16.1 % | 0.187 | 12.8 % |
| response | AN ⇔SN | [-400, 0] | dIES | $\Psi_{N1\Leftrightarrow N2}(E) - \Psi_{N1\Leftrightarrow N2}(F)$ | $\delta$ | 5.17e-04 | 41.1 % | 0.009 | 37.5 % | 0.007 | 41.4 % |
| response | AN ⇔SN | [-400, 0] | dIES | $\Psi_{N1\Leftrightarrow N2}(E) - \Psi_{N1\Leftrightarrow N2}(F)$ | $\theta$ | 0.026 | 23.8 % | 0.237 | 9.7 % | 0.063 | 22.6 % |
| response | AN ⇔SN | [-400, 0] | dIES | $\Psi_{N1\Leftrightarrow N2}(E) - \Psi_{N1\Leftrightarrow N2}(F)$ | $\alpha_1$ | 0.040 | 20.3 % | 0.314 | 7.4 % | 0.067 | 22.1 % |

Continued on next page

**Table 4 – continued from previous page**

| Locked to | Network | Time window | DV | IV | Frequency band | All musicians p | All musicians $R^2$ | Improvisers p | Improvisers $R^2$ | Classical performers p | Classical performers $R^2$ |
|---|---|---|---|---|---|---|---|---|---|---|---|
| response | AN ⇔SN | [-400, 0] | dIES | $\Psi_{N1\Leftrightarrow N2}(E) - \Psi_{N1\Leftrightarrow N2}(F)$ | $\alpha_2$ | 0.025 | 20.6 % | 0.079 | 18.1 % | 0.105 | 17.7 % |
| response | AN ⇔SN | [-400, 0] | dIES | $\Psi_{N1\Leftrightarrow N2}(E) - \Psi_{N1\Leftrightarrow N2}(F)$ | $\beta_1$ | 0.027 | 25.5 % | 0.206 | 10.7 % | 0.199 | 12.1 % |
| response | AN ⇔SN | [-400, 0] | dIES | $\Psi_{N1\Leftrightarrow N2}(E) - \Psi_{N1\Leftrightarrow N2}(F)$ | $\gamma_1$ | 0.002 | 31.2 % | 0.012 | 30.5 % | 0.009 | 39.3 % |
| response | AN ⇔SN | [0, 400] | dIES | $\Psi_{N1\Leftrightarrow N2}(E) - \Psi_{N1\Leftrightarrow N2}(F)$ | $\alpha_1$ | 0.032 | 29.2 % | 0.039 | 24.0 % | 0.467 | 5.0 % |
| response | AN ⇔SN | [0, 400] | dIES | $\Psi_{N1\Leftrightarrow N2}(E) - \Psi_{N1\Leftrightarrow N2}(F)$ | $\alpha_3$ | 0.036 | 31.1 % | 0.019 | 30.7 % | 0.679 | 2.1 % |
| response | AN ⇔SN | [0, 400] | dIES | $\Psi_{N1\Leftrightarrow N2}(E) - \Psi_{N1\Leftrightarrow N2}(F)$ | $\beta_2$ | 0.013 | 32.4 % | 0.016 | 30.1 % | 0.138 | 15.6 % |
| response | AN ⇔SN | [0, 400] | dIES | $\Psi_{N1\Leftrightarrow N2}(E) - \Psi_{N1\Leftrightarrow N2}(F)$ | $\beta_3$ | 0.048 | 23.4 % | 0.445 | 3.7 % | 0.101 | 18.4 % |
| response | AN ⇔SN | [0, 400] | dIES | $\Psi_{N1\Leftrightarrow N2}(E) - \Psi_{N1\Leftrightarrow N2}(F)$ | $\gamma_2$ | 0.030 | 25.1 % | 0.135 | 12.7 % | 0.104 | 18.0 % |
| response | AN ⇔SN | [0, 400] | dIES | $\Psi_{N1\Leftrightarrow N2}(E) - \Psi_{N1\Leftrightarrow N2}(F)$ | $\gamma_3$ | 0.015 | 27.8 % | 0.013 | 30.8 % | 0.381 | 6.2 % |
| response | AN ⇔SN | [0, 400] | imp. hours | $\Psi_{N1\Leftrightarrow N2}(E) - \Psi_{N1\Leftrightarrow N2}(F)$ | $\gamma_1$ | 0.015 | 17.0 % | 0.435 | 6.4 % | 1.000 | 40.9 % |
| response | CCN ⇔SN | [-400, 0] | dIES | $\Psi_{N1\Leftrightarrow N2}(E) - \Psi_{N1\Leftrightarrow N2}(F)$ | $\alpha_2$ | 0.005 | 35.8 % | 0.251 | 13.7 % | 0.008 | 40.9 % |
| response | CCN ⇔SN | [-400, 0] | dIES | $\Psi_{N1\Leftrightarrow N2}(E) - \Psi_{N1\Leftrightarrow N2}(F)$ | $\gamma_1$ | 8.85e-04 | 38.4 % | 0.005 | 38.6 % | 0.080 | 20.8 % |
| response | CCN ⇔SN | [0, 400] | dIES | $\Psi_{N1\Leftrightarrow N2}(E) - \Psi_{N1\Leftrightarrow N2}(F)$ | $\beta_2$ | 0.013 | 31.8 % | 0.043 | 21.6 % | 0.009 | 47.9 % |
| response | CCN ⇔SN | [0, 400] | dIES | $\Psi_{N1\Leftrightarrow N2}(E) - \Psi_{N1\Leftrightarrow N2}(F)$ | $\gamma_3$ | 2.21e-05 | 46.7 % | 0.001 | 46.2 % | 0.006 | 43.2 % |
| chord | CCN ⇔FPN-L | [0, 400] | dIES | $\Psi_{N1\Leftrightarrow N2}(E) - \Psi_{N1\Leftrightarrow N2}(F)$ | $\beta_2$ | 0.002 | 23.1 % | 0.108 | 14.4 % | 0.670 | 1.7 % |
| chord | CCN ⇔FPN-L | [0, 400] | dIES | $\Psi_{N1\Leftrightarrow N2}(E) - \Psi_{N1\Leftrightarrow N2}(F)$ | $\beta_4$ | 6.36e-05 | 36.6 % | 0.841 | 1.5 % | 0.010 | 39.2 % |
| chord | CCN ⇔FPN-R | [0, 400] | imp. hours | $\Psi_{N1\Leftrightarrow N2}(E) - \Psi_{N1\Leftrightarrow N2}(F)$ | $\alpha_3$ | 0.008 | 19.0 % | 0.072 | 18.3 % | 1.000 | 40.8 % |
| chord | CCN ⇔FPN-R | [0, 400] | cl. hours | $\Psi_{N1\Leftrightarrow N2}(E) - \Psi_{N1\Leftrightarrow N2}(F)$ | $\delta$ | 0.008 | 17.5 % | 0.514 | 4.2 % | 0.030 | 29.4 % |
| chord | CCN ⇔DAN | [0, 400] | dIES | $\Psi_{N1\Leftrightarrow N2}(E) - \Psi_{N1\Leftrightarrow N2}(F)$ | $\beta_4$ | 3.59e-04 | 30.8 % | 0.916 | 1.6 % | 0.081 | 20.4 % |
| response | CCN ⇔FPN-L | [-400, 0] | dIES | $\Psi_{N1\Leftrightarrow N2}(E) - \Psi_{N1\Leftrightarrow N2}(F)$ | $\delta$ | 0.024 | 35.1 % | 0.008 | 37.1 % | 0.064 | 31.3 % |
| response | CCN ⇔FPN-L | [-400, 0] | dIES | $\Psi_{N1\Leftrightarrow N2}(E) - \Psi_{N1\Leftrightarrow N2}(F)$ | $\gamma_1$ | 0.011 | 26.6 % | 0.007 | 35.0 % | 0.257 | 9.4 % |
| response | CCN ⇔FPN-L | [-400, 0] | dIES | $\Psi_{N1\Leftrightarrow N2}(E) - \Psi_{N1\Leftrightarrow N2}(F)$ | $\gamma_3$ | 0.043 | 21.7 % | 0.345 | 6.3 % | 0.107 | 18.1 % |
| response | CCN ⇔FPN-L | [0, 400] | dIES | $\Psi_{N1\Leftrightarrow N2}(E) - \Psi_{N1\Leftrightarrow N2}(F)$ | $\beta_2$ | 0.032 | 32.1 % | 0.006 | 37.1 % | 0.967 | 0.2 % |
| response | CCN ⇔FPN-L | [0, 400] | dIES | $\Psi_{N1\Leftrightarrow N2}(E) - \Psi_{N1\Leftrightarrow N2}(F)$ | $\gamma_3$ | 0.013 | 24.6 % | 0.217 | 9.0 % | 0.063 | 22.6 % |
| response | CCN ⇔FPN-L | [0, 400] | imp. hours | $\Psi_{N1\Leftrightarrow N2}(E) - \Psi_{N1\Leftrightarrow N2}(F)$ | $\beta_4$ | 7.77e-04 | 27.2 % | 0.013 | 31.4 % | 1.000 | 41.9 % |
| response | CCN ⇔FPN-R | [-400, 0] | dIES | $\Psi_{N1\Leftrightarrow N2}(E) - \Psi_{N1\Leftrightarrow N2}(F)$ | $\delta$ | 0.029 | 22.1 % | 0.106 | 15.9 % | 0.258 | 9.7 % |
| response | CCN ⇔FPN-R | [-400, 0] | dIES | $\Psi_{N1\Leftrightarrow N2}(E) - \Psi_{N1\Leftrightarrow N2}(F)$ | $\alpha_1$ | 0.008 | 24.7 % | 0.124 | 13.6 % | 0.052 | 24.3 % |
| response | CCN ⇔FPN-R | [-400, 0] | dIES | $\Psi_{N1\Leftrightarrow N2}(E) - \Psi_{N1\Leftrightarrow N2}(F)$ | $\alpha_2$ | 0.003 | 28.6 % | 0.144 | 11.9 % | 0.014 | 35.8 % |

Continued on next page

**Table 4 – continued from previous page**

| Locked to | Network | Time window | DV | IV | Frequency band | All musicians p | All musicians $R^2$ | Improvisers p | Improvisers $R^2$ | Classical performers p | Classical performers $R^2$ |
|---|---|---|---|---|---|---|---|---|---|---|---|
| response | CCN ⇔FPN-R | [-400, 0] | dIES | $\Psi_{N1\Leftrightarrow N2}(E) - \Psi_{N1\Leftrightarrow N2}(F)$ | $\gamma_2$ | 0.025 | 28.2 % | 0.697 | 2.1 % | 0.007 | 41.4 % |
| response | CCN ⇔FPN-R | [-400, 0] | imp. hours | $\Psi_{N1\Leftrightarrow N2}(E) - \Psi_{N1\Leftrightarrow N2}(F)$ | $\alpha_2$ | 0.003 | 22.0 % | 0.067 | 18.2 % | 1.000 | 41.0 % |
| response | CCN ⇔FPN-R | [0, 400] | dIES | $\Psi_{N1\Leftrightarrow N2}(E) - \Psi_{N1\Leftrightarrow N2}(F)$ | $\alpha_3$ | 0.014 | 32.3 % | 0.054 | 22.8 % | 0.160 | 15.0 % |
| response | CCN ⇔FPN-R | [0, 400] | dIES | $\Psi_{N1\Leftrightarrow N2}(E) - \Psi_{N1\Leftrightarrow N2}(F)$ | $\beta_3$ | 3.62e-05 | 45.5 % | 0.119 | 13.1 % | 0.242 | 9.7 % |
| response | CCN ⇔FPN-R | [0, 400] | dIES | $\Psi_{N1\Leftrightarrow N2}(E) - \Psi_{N1\Leftrightarrow N2}(F)$ | $\gamma_3$ | 7.38e-04 | 35.8 % | 0.006 | 35.6 % | 0.207 | 11.2 % |
| response | CCN ⇔FPN-R | [0, 400] | cl. hours | $\Psi_{N1\Leftrightarrow N2}(E) - \Psi_{N1\Leftrightarrow N2}(F)$ | $\gamma_3$ | 0.004 | 21.5 % | 0.107 | 15.4 % | 0.363 | 6.0 % |
| response | CCN ⇔DAN | [-400, 0] | dIES | $\Psi_{N1\Leftrightarrow N2}(E) - \Psi_{N1\Leftrightarrow N2}(F)$ | $\alpha_2$ | 0.011 | 21.7 % | 0.201 | 9.7 % | 0.056 | 23.7 % |
| response | CCN ⇔DAN | [-400, 0] | dIES | $\Psi_{N1\Leftrightarrow N2}(E) - \Psi_{N1\Leftrightarrow N2}(F)$ | $\beta_1$ | 0.037 | 27.4 % | 0.524 | 5.5 % | 0.016 | 35.2 % |
| response | CCN ⇔DAN | [-400, 0] | dIES | $\Psi_{N1\Leftrightarrow N2}(E) - \Psi_{N1\Leftrightarrow N2}(F)$ | $\gamma_1$ | 0.007 | 30.7 % | 0.012 | 33.0 % | 0.094 | 18.8 % |
| response | CCN ⇔DAN | [-400, 0] | dIES | $\Psi_{N1\Leftrightarrow N2}(E) - \Psi_{N1\Leftrightarrow N2}(F)$ | $\gamma_2$ | 0.028 | 25.6 % | 0.837 | 1.2 % | 0.010 | 39.3 % |
| response | CCN ⇔DAN | [-400, 0] | dIES | $\Psi_{N1\Leftrightarrow N2}(E) - \Psi_{N1\Leftrightarrow N2}(F)$ | $\gamma_3$ | 0.008 | 28.2 % | 0.257 | 8.6 % | 0.023 | 32.1 % |
| response | CCN ⇔DAN | [-400, 0] | cl. hours | $\Psi_{N1\Leftrightarrow N2}(E) - \Psi_{N1\Leftrightarrow N2}(F)$ | $\beta_4$ | 0.002 | 22.5 % | 0.178 | 11.1 % | 0.003 | 48.9 % |
| response | CCN ⇔DAN | [0, 400] | dIES | $\Psi_{N1\Leftrightarrow N2}(E) - \Psi_{N1\Leftrightarrow N2}(F)$ | $\alpha_1$ | 0.030 | 30.3 % | 0.059 | 21.3 % | 0.195 | 15.1 % |
| response | CCN ⇔DAN | [0, 400] | dIES | $\Psi_{N1\Leftrightarrow N2}(E) - \Psi_{N1\Leftrightarrow N2}(F)$ | $\alpha_2$ | 0.028 | 30.2 % | 0.103 | 18.5 % | 0.010 | 38.7 % |
| response | CCN ⇔DAN | [0, 400] | dIES | $\Psi_{N1\Leftrightarrow N2}(E) - \Psi_{N1\Leftrightarrow N2}(F)$ | $\beta_2$ | 0.017 | 34.4 % | 0.008 | 35.0 % | 0.438 | 5.0 % |
| response | CCN ⇔DAN | [0, 400] | dIES | $\Psi_{N1\Leftrightarrow N2}(E) - \Psi_{N1\Leftrightarrow N2}(F)$ | $\gamma_3$ | 0.015 | 27.4 % | 0.246 | 8.1 % | 0.038 | 27.2 % |
| response | CCN ⇔DAN | [0, 400] | cl. hours | $\Psi_{N1\Leftrightarrow N2}(E) - \Psi_{N1\Leftrightarrow N2}(F)$ | $\gamma_3$ | 0.008 | 17.7 % | 0.011 | 30.9 % | 0.977 | 0.0 % |
| chord | CCN ⇔DAN-core | [0, 400] | dIES | $\Psi_{N1\Leftrightarrow N2}(E) - \Psi_{N1\Leftrightarrow N2}(F)$ | $\beta_4$ | 0.002 | 24.6 % | 0.933 | 1.5 % | 0.043 | 26.4 % |
| chord | CCN ⇔VN | [0, 400] | dIES | $\Psi_{N1\Leftrightarrow N2}(E) - \Psi_{N1\Leftrightarrow N2}(F)$ | $\beta_3$ | 0.006 | 19.5 % | 0.968 | 1.1 % | 0.479 | 4.1 % |
| chord | CCN ⇔VN | [0, 400] | dIES | $\Psi_{N1\Leftrightarrow N2}(E) - \Psi_{N1\Leftrightarrow N2}(F)$ | $\gamma_2$ | 0.018 | 25.4 % | 0.113 | 13.5 % | 0.050 | 28.6 % |
| chord | CCN ⇔VN | [0, 400] | dIES | $\Psi_{N1\Leftrightarrow N2}(E) - \Psi_{N1\Leftrightarrow N2}(F)$ | $\gamma_3$ | 0.034 | 22.3 % | 0.380 | 5.1 % | 0.035 | 28.7 % |
| response | CCN ⇔DAN-core | [-400, 0] | dIES | $\Psi_{N1\Leftrightarrow N2}(E) - \Psi_{N1\Leftrightarrow N2}(F)$ | $\alpha_1$ | 0.009 | 20.2 % | 0.598 | 4.6 % | 0.099 | 18.2 % |
| response | CCN ⇔DAN-core | [-400, 0] | dIES | $\Psi_{N1\Leftrightarrow N2}(E) - \Psi_{N1\Leftrightarrow N2}(F)$ | $\alpha_2$ | 0.011 | 20.9 % | 0.364 | 5.3 % | 0.027 | 30.3 % |
| response | CCN ⇔DAN-core | [-400, 0] | dIES | $\Psi_{N1\Leftrightarrow N2}(E) - \Psi_{N1\Leftrightarrow N2}(F)$ | $\gamma_1$ | 0.014 | 27.6 % | 0.044 | 22.6 % | 0.093 | 19.0 % |
| response | CCN ⇔DAN-core | [-400, 0] | dIES | $\Psi_{N1\Leftrightarrow N2}(E) - \Psi_{N1\Leftrightarrow N2}(F)$ | $\gamma_2$ | 0.015 | 29.9 % | 0.469 | 5.3 % | 0.014 | 36.1 % |
| response | CCN ⇔DAN-core | [-400, 0] | dIES | $\Psi_{N1\Leftrightarrow N2}(E) - \Psi_{N1\Leftrightarrow N2}(F)$ | $\gamma_3$ | 0.009 | 23.4 % | 0.501 | 3.9 % | 0.041 | 27.2 % |
| response | CCN ⇔DAN-core | [-400, 0] | imp. hours | $\Psi_{N1\Leftrightarrow N2}(E) - \Psi_{N1\Leftrightarrow N2}(F)$ | $\alpha_3$ | 0.040 | 18.6 % | 0.455 | 5.6 % | 1.000 | 43.3 % |
| response | CCN ⇔DAN-core | [-400, 0] | cl. hours | $\Psi_{N1\Leftrightarrow N2}(E) - \Psi_{N1\Leftrightarrow N2}(F)$ | $\beta_4$ | 0.008 | 17.7 % | 0.217 | 8.8 % | 0.006 | 43.2 % |

Continued on next page

**Table 4 – continued from previous page**

| Locked to | Network | Time window | DV | IV | Frequency band | All musicians p | All musicians $R^2$ | Improvisers p | Improvisers $R^2$ | Classical performers p | Classical performers $R^2$ |
|---|---|---|---|---|---|---|---|---|---|---|---|
| response | CCN ⇔DAN-core | [0, 400] | dIES | $\Psi_{N1\Leftrightarrow N2}(E) - \Psi_{N1\Leftrightarrow N2}(F)$ | $\alpha_3$ | 0.033 | 32.0 % | 0.015 | 31.0 % | 0.699 | 2.1 % |
| response | CCN ⇔DAN-core | [0, 400] | dIES | $\Psi_{N1\Leftrightarrow N2}(E) - \Psi_{N1\Leftrightarrow N2}(F)$ | $\beta_2$ | 0.013 | 34.7 % | 0.002 | 44.1 % | 0.491 | 5.1 % |
| response | CCN ⇔DAN-core | [0, 400] | dIES | $\Psi_{N1\Leftrightarrow N2}(E) - \Psi_{N1\Leftrightarrow N2}(F)$ | $\gamma_3$ | 0.033 | 23.4 % | 0.253 | 7.8 % | 0.211 | 11.0 % |
| response | CCN ⇔DAN-core | [0, 400] | imp. hours | $\Psi_{N1\Leftrightarrow N2}(E) - \Psi_{N1\Leftrightarrow N2}(F)$ | $\alpha_1$ | 0.007 | 21.2 % | 0.410 | 8.9 % | 1.000 | 41.8 % |
| response | CCN ⇔DAN-core | [0, 400] | cl. hours | $\Psi_{N1\Leftrightarrow N2}(E) - \Psi_{N1\Leftrightarrow N2}(F)$ | $\gamma_3$ | 0.002 | 23.9 % | 0.015 | 28.7 % | 0.581 | 2.3 % |
| response | CCN ⇔VN | [-400, 0] | dIES | $\Psi_{N1\Leftrightarrow N2}(E) - \Psi_{N1\Leftrightarrow N2}(F)$ | $\beta_1$ | 0.021 | 27.6 % | 0.079 | 20.8 % | 0.042 | 26.5 % |
| response | CCN ⇔VN | [-400, 0] | cl. hours | $\Psi_{N1\Leftrightarrow N2}(E) - \Psi_{N1\Leftrightarrow N2}(F)$ | $\beta_3$ | 0.010 | 16.8 % | 0.210 | 12.4 % | 0.389 | 5.4 % |
| response | CCN ⇔VN | [-400, 0] | cl. hours | $\Psi_{N1\Leftrightarrow N2}(E) - \Psi_{N1\Leftrightarrow N2}(F)$ | $\gamma_1$ | 0.001 | 25.6 % | 5.89e-04 | 51.1 % | 0.554 | 2.6 % |
| response | CCN ⇔VN | [0, 400] | dIES | $\Psi_{N1\Leftrightarrow N2}(E) - \Psi_{N1\Leftrightarrow N2}(F)$ | $\alpha_2$ | 0.017 | 32.2 % | 0.049 | 23.8 % | 0.011 | 38.0 % |
| response | CCN ⇔VN | [0, 400] | dIES | $\Psi_{N1\Leftrightarrow N2}(E) - \Psi_{N1\Leftrightarrow N2}(F)$ | $\beta_2$ | 0.020 | 27.6 % | 0.010 | 32.7 % | 0.232 | 10.3 % |
| response | CCN ⇔VN | [0, 400] | imp. hours | $\Psi_{N1\Leftrightarrow N2}(E) - \Psi_{N1\Leftrightarrow N2}(F)$ | $\gamma_1$ | 0.014 | 19.2 % | 0.649 | 4.0 % | 1.000 | 40.6 % |
| response | CCN ⇔VN | [0, 400] | imp. hours | $\Psi_{N1\Leftrightarrow N2}(E) - \Psi_{N1\Leftrightarrow N2}(F)$ | $\gamma_3$ | 0.011 | 21.8 % | 0.826 | 4.0 % | 1.000 | 41.1 % |
| response | CCN ⇔VN | [0, 400] | cl. hours | $\Psi_{N1\Leftrightarrow N2}(E) - \Psi_{N1\Leftrightarrow N2}(F)$ | $\alpha_1$ | 0.004 | 20.8 % | 0.049 | 20.5 % | 0.087 | 19.5 % |
| chord | DMN ⇔FPN-L | [0, 400] | dIES | $\Psi_{N1\Leftrightarrow N2}(E) - \Psi_{N1\Leftrightarrow N2}(F)$ | $\beta_4$ | 1.15e-05 | 45.7 % | 0.600 | 2.2 % | 0.022 | 33.4 % |
| chord | DMN ⇔DAN | [0, 400] | dIES | $\Psi_{N1\Leftrightarrow N2}(E) - \Psi_{N1\Leftrightarrow N2}(F)$ | $\beta_4$ | 3.78e-04 | 30.6 % | 0.111 | 18.9 % | 0.019 | 33.4 % |
| chord | DMN ⇔DAN | [0, 400] | dIES | $\Psi_{N1\Leftrightarrow N2}(E) - \Psi_{N1\Leftrightarrow N2}(F)$ | $\gamma_2$ | 0.036 | 21.0 % | 0.309 | 6.0 % | 0.029 | 30.1 % |
| response | DMN ⇔FPN-L | [-400, 0] | dIES | $\Psi_{N1\Leftrightarrow N2}(E) - \Psi_{N1\Leftrightarrow N2}(F)$ | $\alpha_2$ | 0.010 | 23.9 % | 0.169 | 11.3 % | 0.039 | 27.0 % |
| response | DMN ⇔FPN-L | [-400, 0] | dIES | $\Psi_{N1\Leftrightarrow N2}(E) - \Psi_{N1\Leftrightarrow N2}(F)$ | $\beta_2$ | 0.005 | 20.3 % | 0.877 | 0.8 % | 0.354 | 7.2 % |
| response | DMN ⇔FPN-L | [-400, 0] | dIES | $\Psi_{N1\Leftrightarrow N2}(E) - \Psi_{N1\Leftrightarrow N2}(F)$ | $\gamma_1$ | 0.048 | 20.8 % | 0.093 | 15.9 % | 0.299 | 7.8 % |
| response | DMN ⇔FPN-L | [-400, 0] | imp. hours | $\Psi_{N1\Leftrightarrow N2}(E) - \Psi_{N1\Leftrightarrow N2}(F)$ | $\alpha_1$ | 0.004 | 22.2 % | 0.600 | 4.0 % | 1.000 | 40.6 % |
| response | DMN ⇔FPN-L | [0, 400] | dIES | $\Psi_{N1\Leftrightarrow N2}(E) - \Psi_{N1\Leftrightarrow N2}(F)$ | $\beta_2$ | 0.026 | 26.9 % | 0.011 | 32.4 % | 0.465 | 4.1 % |
| response | DMN ⇔FPN-R | [-400, 0] | dIES | $\Psi_{N1\Leftrightarrow N2}(E) - \Psi_{N1\Leftrightarrow N2}(F)$ | $\alpha_2$ | 0.029 | 18.0 % | 0.095 | 14.7 % | 0.259 | 9.1 % |
| response | DMN ⇔FPN-R | [-400, 0] | dIES | $\Psi_{N1\Leftrightarrow N2}(E) - \Psi_{N1\Leftrightarrow N2}(F)$ | $\gamma_1$ | 0.012 | 25.6 % | 0.807 | 2.1 % | 0.003 | 48.8 % |
| response | DMN ⇔FPN-R | [0, 400] | dIES | $\Psi_{N1\Leftrightarrow N2}(E) - \Psi_{N1\Leftrightarrow N2}(F)$ | $\beta_1$ | 0.044 | 24.2 % | 0.399 | 5.1 % | 0.029 | 29.6 % |
| response | DMN ⇔FPN-R | [0, 400] | dIES | $\Psi_{N1\Leftrightarrow N2}(E) - \Psi_{N1\Leftrightarrow N2}(F)$ | $\beta_3$ | 5.04e-04 | 38.4 % | 0.124 | 12.7 % | 0.060 | 23.1 % |
| response | DMN ⇔DAN | [-400, 0] | dIES | $\Psi_{N1\Leftrightarrow N2}(E) - \Psi_{N1\Leftrightarrow N2}(F)$ | $\alpha_1$ | 0.012 | 27.3 % | 0.031 | 27.3 % | 0.159 | 13.7 % |
| response | DMN ⇔DAN | [-400, 0] | dIES | $\Psi_{N1\Leftrightarrow N2}(E) - \Psi_{N1\Leftrightarrow N2}(F)$ | $\alpha_2$ | 0.012 | 23.4 % | 0.074 | 17.0 % | 0.066 | 22.1 % |
| response | DMN ⇔DAN | [-400, 0] | dIES | $\Psi_{N1\Leftrightarrow N2}(E) - \Psi_{N1\Leftrightarrow N2}(F)$ | $\gamma_1$ | 0.005 | 28.1 % | 0.069 | 18.3 % | 0.048 | 25.2 % |

Continued on next page

**Table 4 – continued from previous page**

| Locked to | Network | Time window | DV | IV | Frequency band | All musicians p | All musicians $R^2$ | Improvisers p | Improvisers $R^2$ | Classical performers p | Classical performers $R^2$ |
|---|---|---|---|---|---|---|---|---|---|---|---|
| response | DMN ⇔DAN | [-400, 0] | dIES | $\Psi_{N1\Leftrightarrow N2}(E) - \Psi_{N1\Leftrightarrow N2}(F)$ | $\gamma_2$ | 0.029 | 30.5 % | 0.351 | 7.5 % | 0.017 | 37.3 % |
| response | DMN ⇔DAN | [-400, 0] | imp. hours | $\Psi_{N1\Leftrightarrow N2}(E) - \Psi_{N1\Leftrightarrow N2}(F)$ | $\alpha_1$ | 3.06e-04 | 32.0 % | 0.003 | 45.1 % | 1.000 | 41.3 % |
| response | DMN ⇔DAN | [-400, 0] | imp. hours | $\Psi_{N1\Leftrightarrow N2}(E) - \Psi_{N1\Leftrightarrow N2}(F)$ | $\alpha_3$ | 0.011 | 24.0 % | 0.726 | 3.0 % | 1.000 | 40.8 % |
| response | DMN ⇔DAN | [-400, 0] | cl. hours | $\Psi_{N1\Leftrightarrow N2}(E) - \Psi_{N1\Leftrightarrow N2}(F)$ | $\beta_3$ | 0.001 | 24.6 % | 0.029 | 24.2 % | 0.344 | 6.4 % |
| response | DMN ⇔DAN | [0, 400] | dIES | $\Psi_{N1\Leftrightarrow N2}(E) - \Psi_{N1\Leftrightarrow N2}(F)$ | $\beta_2$ | 0.002 | 41.7 % | 0.002 | 43.4 % | 0.058 | 23.5 % |
| response | DMN ⇔DAN | [0, 400] | dIES | $\Psi_{N1\Leftrightarrow N2}(E) - \Psi_{N1\Leftrightarrow N2}(F)$ | $\beta_3$ | 9.05e-04 | 34.9 % | 0.004 | 38.2 % | 0.113 | 17.6 % |
| response | DMN ⇔DAN | [0, 400] | imp. hours | $\Psi_{N1\Leftrightarrow N2}(E) - \Psi_{N1\Leftrightarrow N2}(F)$ | $\beta_4$ | 0.007 | 18.5 % | 0.272 | 8.0 % | 1.000 | 44.8 % |
| response | DMN ⇔DAN | [0, 400] | imp. hours | $\Psi_{N1\Leftrightarrow N2}(E) - \Psi_{N1\Leftrightarrow N2}(F)$ | $\gamma_3$ | 0.010 | 21.5 % | 0.562 | 3.8 % | 1.000 | 41.4 % |
| chord | DMN ⇔DAN-core | [0, 400] | dIES | $\Psi_{N1\Leftrightarrow N2}(E) - \Psi_{N1\Leftrightarrow N2}(F)$ | $\beta_4$ | 2.18e-04 | 32.3 % | 0.987 | 1.9 % | 0.031 | 29.0 % |
| chord | DMN ⇔DAN-core | [0, 400] | dIES | $\Psi_{N1\Leftrightarrow N2}(E) - \Psi_{N1\Leftrightarrow N2}(F)$ | $\gamma_1$ | 0.031 | 18.2 % | 0.243 | 8.1 % | 0.094 | 19.0 % |
| chord | DMN ⇔VN | [0, 400] | dIES | $\Psi_{N1\Leftrightarrow N2}(E) - \Psi_{N1\Leftrightarrow N2}(F)$ | $\beta_3$ | 7.56e-06 | 42.6 % | 0.491 | 5.1 % | 0.250 | 9.8 % |
| chord | DMN ⇔VN | [0, 400] | dIES | $\Psi_{N1\Leftrightarrow N2}(E) - \Psi_{N1\Leftrightarrow N2}(F)$ | $\gamma_1$ | 0.024 | 31.4 % | 0.041 | 25.0 % | 0.206 | 12.0 % |
| chord | DMN ⇔VN | [0, 400] | dIES | $\Psi_{N1\Leftrightarrow N2}(E) - \Psi_{N1\Leftrightarrow N2}(F)$ | $\gamma_2$ | 0.032 | 25.3 % | 0.309 | 6.1 % | 0.047 | 26.2 % |
| response | DMN ⇔DAN-core | [-400, 0] | dIES | $\Psi_{N1\Leftrightarrow N2}(E) - \Psi_{N1\Leftrightarrow N2}(F)$ | $\gamma_1$ | 0.012 | 27.0 % | 0.063 | 18.4 % | 0.082 | 20.1 % |
| response | DMN ⇔DAN-core | [-400, 0] | dIES | $\Psi_{N1\Leftrightarrow N2}(E) - \Psi_{N1\Leftrightarrow N2}(F)$ | $\gamma_2$ | 0.020 | 33.0 % | 0.293 | 8.8 % | 0.012 | 40.4 % |
| response | DMN ⇔DAN-core | [-400, 0] | imp. hours | $\Psi_{N1\Leftrightarrow N2}(E) - \Psi_{N1\Leftrightarrow N2}(F)$ | $\alpha_1$ | 0.007 | 23.3 % | 0.072 | 24.8 % | 1.000 | 40.5 % |
| response | DMN ⇔DAN-core | [-400, 0] | cl. hours | $\Psi_{N1\Leftrightarrow N2}(E) - \Psi_{N1\Leftrightarrow N2}(F)$ | $\beta_3$ | 0.005 | 19.5 % | 0.277 | 9.3 % | 0.396 | 5.2 % |
| response | DMN ⇔DAN-core | [0, 400] | dIES | $\Psi_{N1\Leftrightarrow N2}(E) - \Psi_{N1\Leftrightarrow N2}(F)$ | $\beta_2$ | 0.019 | 30.3 % | 0.001 | 47.5 % | 0.299 | 7.8 % |
| response | DMN ⇔DAN-core | [0, 400] | imp. hours | $\Psi_{N1\Leftrightarrow N2}(E) - \Psi_{N1\Leftrightarrow N2}(F)$ | $\beta_4$ | 0.003 | 22.1 % | 0.507 | 6.4 % | 1.000 | 44.1 % |
| response | DMN ⇔VN | [-400, 0] | dIES | $\Psi_{N1\Leftrightarrow N2}(E) - \Psi_{N1\Leftrightarrow N2}(F)$ | $\alpha_2$ | 0.020 | 24.6 % | 0.089 | 16.6 % | 0.044 | 25.9 % |
| response | DMN ⇔VN | [-400, 0] | dIES | $\Psi_{N1\Leftrightarrow N2}(E) - \Psi_{N1\Leftrightarrow N2}(F)$ | $\beta_2$ | 0.046 | 18.1 % | 0.445 | 5.4 % | 0.025 | 31.3 % |
| response | DMN ⇔VN | [-400, 0] | imp. hours | $\Psi_{N1\Leftrightarrow N2}(E) - \Psi_{N1\Leftrightarrow N2}(F)$ | $\alpha_2$ | 0.010 | 19.5 % | 0.050 | 25.8 % | 1.000 | 40.9 % |
| response | DMN ⇔VN | [-400, 0] | imp. hours | $\Psi_{N1\Leftrightarrow N2}(E) - \Psi_{N1\Leftrightarrow N2}(F)$ | $\alpha_3$ | 0.002 | 26.8 % | 0.008 | 42.4 % | 1.000 | 40.9 % |
| response | DMN ⇔VN | [0, 400] | dIES | $\Psi_{N1\Leftrightarrow N2}(E) - \Psi_{N1\Leftrightarrow N2}(F)$ | $\alpha_3$ | 0.050 | 26.0 % | 0.026 | 28.6 % | 0.171 | 13.2 % |
| response | DMN ⇔VN | [0, 400] | dIES | $\Psi_{N1\Leftrightarrow N2}(E) - \Psi_{N1\Leftrightarrow N2}(F)$ | $\beta_1$ | 0.035 | 26.9 % | 0.139 | 14.6 % | 0.116 | 16.9 % |
| response | DMN ⇔VN | [0, 400] | dIES | $\Psi_{N1\Leftrightarrow N2}(E) - \Psi_{N1\Leftrightarrow N2}(F)$ | $\beta_2$ | 0.008 | 31.3 % | 0.060 | 19.3 % | 0.033 | 28.7 % |
| response | DMN ⇔VN | [0, 400] | dIES | $\Psi_{N1\Leftrightarrow N2}(E) - \Psi_{N1\Leftrightarrow N2}(F)$ | $\beta_3$ | 1.23e-04 | 36.7 % | 0.004 | 37.4 % | 0.037 | 27.7 % |
| response | DMN ⇔VN | [0, 400] | imp. hours | $\Psi_{N1\Leftrightarrow N2}(E) - \Psi_{N1\Leftrightarrow N2}(F)$ | $\gamma_3$ | 0.007 | 20.8 % | 0.694 | 5.9 % | 1.000 | 41.1 % |

Continued on next page

**Table 4 – continued from previous page**

| Locked to | Network | Time window | DV | IV | Frequency band | All musicians p | All musicians $R^2$ | Improvisers p | Improvisers $R^2$ | Classical performers p | Classical performers $R^2$ |
|---|---|---|---|---|---|---|---|---|---|---|---|
| response | DMN ⇔VN | [0, 400] | cl. hours | $\Psi_{N1\Leftrightarrow N2}(E) - \Psi_{N1\Leftrightarrow N2}(F)$ | $\beta_3$ | 0.005 | 19.3 % | 0.198 | 11.4 % | 0.005 | 44.8 % |
| chord | SN ⇔FPN-L | [0, 400] | dIES | $\Psi_{N1\Leftrightarrow N2}(E) - \Psi_{N1\Leftrightarrow N2}(F)$ | $\beta_2$ | 0.022 | 25.2 % | 0.028 | 24.2 % | 0.483 | 4.3 % |
| chord | SN ⇔FPN-L | [0, 400] | dIES | $\Psi_{N1\Leftrightarrow N2}(E) - \Psi_{N1\Leftrightarrow N2}(F)$ | $\gamma_1$ | 0.026 | 21.5 % | 0.616 | 5.0 % | 0.012 | 53.4 % |
| chord | SN ⇔FPN-L | [0, 400] | cl. hours | $\Psi_{N1\Leftrightarrow N2}(E) - \Psi_{N1\Leftrightarrow N2}(F)$ | $\alpha_3$ | 0.008 | 17.8 % | 0.048 | 21.3 % | 0.268 | 8.7 % |
| chord | SN ⇔FPN-L | [0, 400] | cl. hours | $\Psi_{N1\Leftrightarrow N2}(E) - \Psi_{N1\Leftrightarrow N2}(F)$ | $\gamma_3$ | 0.005 | 19.4 % | 0.017 | 28.9 % | 0.004 | 46.3 % |
| chord | SN ⇔FPN-R | [0, 400] | dIES | $\Psi_{N1\Leftrightarrow N2}(E) - \Psi_{N1\Leftrightarrow N2}(F)$ | $\beta_3$ | 0.007 | 19.8 % | 0.817 | 1.1 % | 0.213 | 11.1 % |
| chord | SN ⇔FPN-R | [0, 400] | dIES | $\Psi_{N1\Leftrightarrow N2}(E) - \Psi_{N1\Leftrightarrow N2}(F)$ | $\beta_4$ | 3.13e-04 | 31.3 % | 0.276 | 7.0 % | 0.023 | 31.9 % |
| chord | SN ⇔FPN-R | [0, 400] | cl. hours | $\Psi_{N1\Leftrightarrow N2}(E) - \Psi_{N1\Leftrightarrow N2}(F)$ | $\delta$ | 0.001 | 25.6 % | 0.005 | 40.1 % | 0.062 | 22.7 % |
| chord | SN ⇔DAN | [0, 400] | dIES | $\Psi_{N1\Leftrightarrow N2}(E) - \Psi_{N1\Leftrightarrow N2}(F)$ | $\beta_2$ | 0.005 | 26.7 % | 0.001 | 46.4 % | 0.799 | 1.5 % |
| response | SN ⇔FPN-L | [-400, 0] | dIES | $\Psi_{N1\Leftrightarrow N2}(E) - \Psi_{N1\Leftrightarrow N2}(F)$ | $\alpha_2$ | 0.034 | 24.2 % | 0.048 | 28.5 % | 0.047 | 25.3 % |
| response | SN ⇔FPN-L | [-400, 0] | dIES | $\Psi_{N1\Leftrightarrow N2}(E) - \Psi_{N1\Leftrightarrow N2}(F)$ | $\gamma_1$ | 0.022 | 22.5 % | 0.030 | 26.1 % | 0.176 | 12.9 % |
| response | SN ⇔FPN-L | [-400, 0] | dIES | $\Psi_{N1\Leftrightarrow N2}(E) - \Psi_{N1\Leftrightarrow N2}(F)$ | $\gamma_2$ | 0.039 | 25.9 % | 0.649 | 2.2 % | 0.008 | 43.6 % |
| response | SN ⇔FPN-L | [0, 400] | dIES | $\Psi_{N1\Leftrightarrow N2}(E) - \Psi_{N1\Leftrightarrow N2}(F)$ | $\beta_1$ | 0.032 | 30.1 % | 0.023 | 29.0 % | 0.405 | 6.7 % |
| response | SN ⇔FPN-L | [0, 400] | dIES | $\Psi_{N1\Leftrightarrow N2}(E) - \Psi_{N1\Leftrightarrow N2}(F)$ | $\beta_2$ | 0.024 | 30.2 % | 0.013 | 33.3 % | 0.215 | 10.8 % |
| response | SN ⇔FPN-R | [-400, 0] | dIES | $\Psi_{N1\Leftrightarrow N2}(E) - \Psi_{N1\Leftrightarrow N2}(F)$ | $\alpha_2$ | 0.005 | 27.6 % | 0.154 | 11.5 % | 0.004 | 47.9 % |
| response | SN ⇔FPN-R | [-400, 0] | dIES | $\Psi_{N1\Leftrightarrow N2}(E) - \Psi_{N1\Leftrightarrow N2}(F)$ | $\gamma_1$ | 0.009 | 29.1 % | 0.179 | 12.0 % | 0.023 | 31.8 % |
| response | SN ⇔FPN-R | [0, 400] | dIES | $\Psi_{N1\Leftrightarrow N2}(E) - \Psi_{N1\Leftrightarrow N2}(F)$ | $\alpha_2$ | 0.015 | 30.6 % | 0.014 | 33.9 % | 0.306 | 7.5 % |
| response | SN ⇔FPN-R | [0, 400] | dIES | $\Psi_{N1\Leftrightarrow N2}(E) - \Psi_{N1\Leftrightarrow N2}(F)$ | $\gamma_3$ | 0.004 | 33.0 % | 0.005 | 39.1 % | 0.110 | 17.6 % |
| response | SN ⇔DAN | [-400, 0] | dIES | $\Psi_{N1\Leftrightarrow N2}(E) - \Psi_{N1\Leftrightarrow N2}(F)$ | $\beta_2$ | 0.011 | 19.6 % | 0.604 | 2.8 % | 0.035 | 28.1 % |
| response | SN ⇔DAN | [-400, 0] | dIES | $\Psi_{N1\Leftrightarrow N2}(E) - \Psi_{N1\Leftrightarrow N2}(F)$ | $\gamma_1$ | 2.95e-04 | 38.8 % | 0.002 | 43.8 % | 0.068 | 21.9 % |
| response | SN ⇔DAN | [0, 400] | dIES | $\Psi_{N1\Leftrightarrow N2}(E) - \Psi_{N1\Leftrightarrow N2}(F)$ | $\alpha_1$ | 0.045 | 26.6 % | 0.046 | 21.9 % | 0.594 | 3.1 % |
| response | SN ⇔DAN | [0, 400] | dIES | $\Psi_{N1\Leftrightarrow N2}(E) - \Psi_{N1\Leftrightarrow N2}(F)$ | $\beta_2$ | 0.004 | 35.4 % | 0.013 | 31.1 % | 0.036 | 28.3 % |
| response | SN ⇔DAN | [0, 400] | dIES | $\Psi_{N1\Leftrightarrow N2}(E) - \Psi_{N1\Leftrightarrow N2}(F)$ | $\beta_3$ | 0.036 | 24.2 % | 0.357 | 4.9 % | 0.235 | 10.3 % |
| response | SN ⇔DAN | [0, 400] | dIES | $\Psi_{N1\Leftrightarrow N2}(E) - \Psi_{N1\Leftrightarrow N2}(F)$ | $\beta_4$ | 0.042 | 21.8 % | 0.001 | 45.2 % | 0.680 | 1.5 % |
| response | SN ⇔DAN | [0, 400] | dIES | $\Psi_{N1\Leftrightarrow N2}(E) - \Psi_{N1\Leftrightarrow N2}(F)$ | $\gamma_3$ | 0.002 | 34.1 % | 0.020 | 27.4 % | 0.051 | 24.5 % |
| response | SN ⇔DAN | [0, 400] | imp. hours | $\Psi_{N1\Leftrightarrow N2}(E) - \Psi_{N1\Leftrightarrow N2}(F)$ | $\alpha_1$ | 0.009 | 20.0 % | 0.129 | 18.4 % | 1.000 | 42.7 % |
| response | SN ⇔DAN | [0, 400] | imp. hours | $\Psi_{N1\Leftrightarrow N2}(E) - \Psi_{N1\Leftrightarrow N2}(F)$ | $\alpha_2$ | 0.031 | 16.9 % | 0.508 | 6.9 % | 1.000 | 40.7 % |
| chord | DAN-core ⇔SN-core | [0, 400] | dIES | $\Psi_{N1\Leftrightarrow N2}(E) - \Psi_{N1\Leftrightarrow N2}(F)$ | $\beta_2$ | 0.012 | 26.1 % | 0.023 | 27.1 % | 0.086 | 19.6 % |

Continued on next page

**Table 4 – continued from previous page**

| Locked to | Network | Time window | DV | IV | Frequency band | All musicians p | All musicians $R^2$ | Improvisers p | Improvisers $R^2$ | Classical performers p | Classical performers $R^2$ |
|---|---|---|---|---|---|---|---|---|---|---|---|
| chord | DAN-core ⇔SN-core | [0, 400] | dIES | $\Psi_{N1\Leftrightarrow N2}(E) - \Psi_{N1\Leftrightarrow N2}(F)$ | $\beta_4$ | 0.003 | 24.3 % | 0.345 | 6.3 % | 0.003 | 47.4 % |
| chord | DAN-core ⇔SN-core | [0, 400] | cl. hours | $\Psi_{N1\Leftrightarrow N2}(E) - \Psi_{N1\Leftrightarrow N2}(F)$ | $\gamma_3$ | 0.009 | 17.2 % | 0.014 | 29.1 % | 0.020 | 33.2 % |
| chord | DAN-core ⇔CON | [0, 400] | dIES | $\Psi_{N1\Leftrightarrow N2}(E) - \Psi_{N1\Leftrightarrow N2}(F)$ | $\beta_4$ | 0.004 | 23.9 % | 0.264 | 8.3 % | 0.002 | 54.9 % |
| chord | DAN-core ⇔CON | [0, 400] | imp. hours | $\Psi_{N1\Leftrightarrow N2}(E) - \Psi_{N1\Leftrightarrow N2}(F)$ | $\beta_1$ | 0.017 | 19.9 % | 0.042 | 30.3 % | 1.000 | 46.9 % |
| chord | DAN-core ⇔CON | [0, 400] | cl. hours | $\Psi_{N1\Leftrightarrow N2}(E) - \Psi_{N1\Leftrightarrow N2}(F)$ | $\gamma_3$ | 0.008 | 17.7 % | 0.258 | 8.3 % | 0.005 | 44.7 % |
| chord | VN ⇔SN-core | [0, 400] | dIES | $\Psi_{N1\Leftrightarrow N2}(E) - \Psi_{N1\Leftrightarrow N2}(F)$ | $\beta_2$ | 0.023 | 22.8 % | 0.022 | 25.9 % | 0.479 | 3.9 % |
| chord | VN ⇔SN-core | [0, 400] | imp. hours | $\Psi_{N1\Leftrightarrow N2}(E) - \Psi_{N1\Leftrightarrow N2}(F)$ | $\alpha_2$ | 0.010 | 18.5 % | 0.014 | 31.8 % | 1.000 | 55.3 % |
| chord | VN ⇔SN-core | [0, 400] | imp. hours | $\Psi_{N1\Leftrightarrow N2}(E) - \Psi_{N1\Leftrightarrow N2}(F)$ | $\beta_4$ | 0.002 | 27.5 % | 0.909 | 3.6 % | 1.000 | 45.7 % |
| chord | VN ⇔CON | [0, 400] | dIES | $\Psi_{N1\Leftrightarrow N2}(E) - \Psi_{N1\Leftrightarrow N2}(F)$ | $\beta_2$ | 0.015 | 22.1 % | 0.016 | 28.2 % | 0.615 | 2.7 % |
| chord | VN ⇔CON | [0, 400] | dIES | $\Psi_{N1\Leftrightarrow N2}(E) - \Psi_{N1\Leftrightarrow N2}(F)$ | $\gamma_3$ | 0.025 | 24.5 % | 0.206 | 10.4 % | 0.086 | 19.9 % |
| response | DAN-core ⇔SN-core | [-400, 0] | dIES | $\Psi_{N1\Leftrightarrow N2}(E) - \Psi_{N1\Leftrightarrow N2}(F)$ | $\alpha_2$ | 0.019 | 29.6 % | 0.572 | 2.3 % | 0.011 | 38.3 % |
| response | DAN-core ⇔SN-core | [-400, 0] | dIES | $\Psi_{N1\Leftrightarrow N2}(E) - \Psi_{N1\Leftrightarrow N2}(F)$ | $\gamma_1$ | 0.031 | 24.9 % | 0.026 | 27.5 % | 0.208 | 11.2 % |
| response | DAN-core ⇔SN-core | [0, 400] | dIES | $\Psi_{N1\Leftrightarrow N2}(E) - \Psi_{N1\Leftrightarrow N2}(F)$ | $\beta_2$ | 0.011 | 35.6 % | 0.015 | 30.5 % | 0.449 | 5.5 % |
| response | DAN-core ⇔SN-core | [0, 400] | dIES | $\Psi_{N1\Leftrightarrow N2}(E) - \Psi_{N1\Leftrightarrow N2}(F)$ | $\beta_3$ | 0.016 | 24.0 % | 0.108 | 13.8 % | 0.284 | 8.3 % |
| response | DAN-core ⇔SN-core | [0, 400] | dIES | $\Psi_{N1\Leftrightarrow N2}(E) - \Psi_{N1\Leftrightarrow N2}(F)$ | $\beta_4$ | 0.022 | 28.0 % | 0.013 | 29.8 % | 0.188 | 12.7 % |
| response | DAN-core ⇔SN-core | [0, 400] | dIES | $\Psi_{N1\Leftrightarrow N2}(E) - \Psi_{N1\Leftrightarrow N2}(F)$ | $\gamma_3$ | 0.017 | 28.6 % | 0.046 | 20.9 % | 0.169 | 14.0 % |
| response | DAN-core ⇔SN-core | [0, 400] | imp. hours | $\Psi_{N1\Leftrightarrow N2}(E) - \Psi_{N1\Leftrightarrow N2}(F)$ | $\beta_3$ | 0.009 | 21.1 % | 0.137 | 13.2 % | 1.000 | 40.5 % |
| response | DAN-core ⇔CON | [-400, 0] | dIES | $\Psi_{N1\Leftrightarrow N2}(E) - \Psi_{N1\Leftrightarrow N2}(F)$ | $\delta$ | 0.042 | 28.3 % | 0.026 | 28.7 % | 0.464 | 5.9 % |
| response | DAN-core ⇔CON | [-400, 0] | dIES | $\Psi_{N1\Leftrightarrow N2}(E) - \Psi_{N1\Leftrightarrow N2}(F)$ | $\alpha_2$ | 0.021 | 28.3 % | 0.132 | 12.2 % | 0.004 | 46.6 % |
| response | DAN-core ⇔CON | [-400, 0] | dIES | $\Psi_{N1\Leftrightarrow N2}(E) - \Psi_{N1\Leftrightarrow N2}(F)$ | $\beta_3$ | 0.043 | 22.1 % | 0.207 | 8.8 % | 0.231 | 11.6 % |
| response | DAN-core ⇔CON | [0, 400] | dIES | $\Psi_{N1\Leftrightarrow N2}(E) - \Psi_{N1\Leftrightarrow N2}(F)$ | $\beta_2$ | 4.98e-04 | 42.8 % | 0.004 | 39.3 % | 0.036 | 27.8 % |
| response | DAN-core ⇔CON | [0, 400] | dIES | $\Psi_{N1\Leftrightarrow N2}(E) - \Psi_{N1\Leftrightarrow N2}(F)$ | $\beta_4$ | 0.028 | 25.3 % | 0.137 | 12.5 % | 0.609 | 2.5 % |
| response | DAN-core ⇔CON | [0, 400] | dIES | $\Psi_{N1\Leftrightarrow N2}(E) - \Psi_{N1\Leftrightarrow N2}(F)$ | $\gamma_3$ | 0.002 | 36.3 % | 0.009 | 33.6 % | 0.163 | 13.9 % |
| response | DAN-core ⇔CON | [0, 400] | imp. hours | $\Psi_{N1\Leftrightarrow N2}(E) - \Psi_{N1\Leftrightarrow N2}(F)$ | $\beta_4$ | 0.005 | 24.7 % | 0.669 | 5.3 % | 1.000 | 40.6 % |
| response | DAN-core ⇔CON | [0, 400] | cl. hours | $\Psi_{N1\Leftrightarrow N2}(E) - \Psi_{N1\Leftrightarrow N2}(F)$ | $\gamma_3$ | 9.27e-04 | 28.0 % | 0.036 | 22.5 % | 0.867 | 0.2 % |
| response | VN ⇔SN-core | [-400, 0] | dIES | $\Psi_{N1\Leftrightarrow N2}(E) - \Psi_{N1\Leftrightarrow N2}(F)$ | $\beta_2$ | 0.004 | 24.4 % | 0.845 | 1.4 % | 0.010 | 38.7 % |
| response | VN ⇔SN-core | [-400, 0] | dIES | $\Psi_{N1\Leftrightarrow N2}(E) - \Psi_{N1\Leftrightarrow N2}(F)$ | $\gamma_1$ | 0.011 | 25.2 % | 0.046 | 21.6 % | 0.391 | 5.6 % |
| response | VN ⇔SN-core | [0, 400] | dIES | $\Psi_{N1\Leftrightarrow N2}(E) - \Psi_{N1\Leftrightarrow N2}(F)$ | $\alpha_3$ | 0.022 | 32.8 % | 0.002 | 47.2 % | 0.454 | 5.5 % |

Continued on next page

**Table 4 – continued from previous page**

| Locked to | Network | Time window | DV | IV | Frequency band | All musicians p | All musicians $R^2$ | Improvisers p | Improvisers $R^2$ | Classical performers p | Classical performers $R^2$ |
|---|---|---|---|---|---|---|---|---|---|---|---|
| response | VN ⇔SN-core | [0, 400] | dIES | $\Psi_{N1\Leftrightarrow N2}(E) - \Psi_{N1\Leftrightarrow N2}(F)$ | $\beta_2$ | 0.037 | 21.9 % | 0.063 | 18.6 % | 0.281 | 8.7 % |
| response | VN ⇔SN-core | [0, 400] | dIES | $\Psi_{N1\Leftrightarrow N2}(E) - \Psi_{N1\Leftrightarrow N2}(F)$ | $\beta_3$ | 0.025 | 23.6 % | 0.270 | 7.6 % | 0.228 | 11.0 % |
| response | VN ⇔SN-core | [0, 400] | dIES | $\Psi_{N1\Leftrightarrow N2}(E) - \Psi_{N1\Leftrightarrow N2}(F)$ | $\gamma_1$ | 0.036 | 20.9 % | 0.042 | 24.3 % | 0.332 | 6.9 % |
| response | VN ⇔SN-core | [0, 400] | dIES | $\Psi_{N1\Leftrightarrow N2}(E) - \Psi_{N1\Leftrightarrow N2}(F)$ | $\gamma_3$ | 0.030 | 24.3 % | 0.059 | 19.0 % | 0.314 | 7.4 % |
| response | VN ⇔SN-core | [0, 400] | imp. hours | $\Psi_{N1\Leftrightarrow N2}(E) - \Psi_{N1\Leftrightarrow N2}(F)$ | $\beta_2$ | 0.004 | 20.5 % | 0.007 | 34.1 % | 1.000 | 41.4 % |
| response | VN ⇔SN-core | [0, 400] | imp. hours | $\Psi_{N1\Leftrightarrow N2}(E) - \Psi_{N1\Leftrightarrow N2}(F)$ | $\gamma_1$ | 0.003 | 28.8 % | 0.467 | 8.2 % | 1.000 | 42.7 % |
| response | VN ⇔SN-core | [0, 400] | cl. hours | $\Psi_{N1\Leftrightarrow N2}(E) - \Psi_{N1\Leftrightarrow N2}(F)$ | $\beta_1$ | 0.011 | 16.8 % | 0.036 | 22.5 % | 0.143 | 14.7 % |
| response | VN ⇔CON | [-400, 0] | dIES | $\Psi_{N1\Leftrightarrow N2}(E) - \Psi_{N1\Leftrightarrow N2}(F)$ | $\beta_2$ | 6.33e-04 | 28.2 % | 0.747 | 1.8 % | 7.39e-04 | 57.7 % |
| response | VN ⇔CON | [-400, 0] | dIES | $\Psi_{N1\Leftrightarrow N2}(E) - \Psi_{N1\Leftrightarrow N2}(F)$ | $\gamma_1$ | 0.040 | 21.5 % | 0.067 | 19.3 % | 0.381 | 5.7 % |
| response | VN ⇔CON | [-400, 0] | cl. hours | $\Psi_{N1\Leftrightarrow N2}(E) - \Psi_{N1\Leftrightarrow N2}(F)$ | $\gamma_1$ | 0.009 | 17.3 % | 0.734 | 7.4 % | 0.046 | 25.8 % |
| response | VN ⇔CON | [0, 400] | dIES | $\Psi_{N1\Leftrightarrow N2}(E) - \Psi_{N1\Leftrightarrow N2}(F)$ | $\alpha_2$ | 0.032 | 28.5 % | 0.095 | 19.3 % | 0.014 | 35.9 % |
| response | VN ⇔CON | [0, 400] | dIES | $\Psi_{N1\Leftrightarrow N2}(E) - \Psi_{N1\Leftrightarrow N2}(F)$ | $\beta_2$ | 0.007 | 30.3 % | 0.029 | 24.4 % | 0.035 | 28.1 % |
| response | VN ⇔CON | [0, 400] | dIES | $\Psi_{N1\Leftrightarrow N2}(E) - \Psi_{N1\Leftrightarrow N2}(F)$ | $\beta_3$ | 0.017 | 22.8 % | 0.439 | 4.3 % | 0.196 | 11.8 % |
| response | VN ⇔CON | [0, 400] | dIES | $\Psi_{N1\Leftrightarrow N2}(E) - \Psi_{N1\Leftrightarrow N2}(F)$ | $\gamma_1$ | 7.42e-04 | 30.4 % | 0.005 | 36.8 % | 0.078 | 20.6 % |
| response | VN ⇔CON | [0, 400] | imp. hours | $\Psi_{N1\Leftrightarrow N2}(E) - \Psi_{N1\Leftrightarrow N2}(F)$ | $\beta_3$ | 0.038 | 20.3 % | 0.369 | 12.9 % | 1.000 | 43.3 % |
| response | VN ⇔CON | [0, 400] | imp. hours | $\Psi_{N1\Leftrightarrow N2}(E) - \Psi_{N1\Leftrightarrow N2}(F)$ | $\gamma_1$ | 0.008 | 20.9 % | 0.239 | 13.3 % | 1.000 | 44.0 % |
| chord | VN ⇔DAN-core | [0, 400] | dIES | $\Psi_{N1\Leftrightarrow N2}(E) - \Psi_{N1\Leftrightarrow N2}(F)$ | $\delta$ | 0.024 | 23.9 % | 0.373 | 7.4 % | 0.117 | 16.7 % |
| chord | VN ⇔DAN-core | [0, 400] | dIES | $\Psi_{N1\Leftrightarrow N2}(E) - \Psi_{N1\Leftrightarrow N2}(F)$ | $\beta_3$ | 3.40e-04 | 29.8 % | 0.698 | 2.3 % | 0.475 | 4.0 % |
| chord | VN ⇔DAN-core | [0, 400] | cl. hours | $\Psi_{N1\Leftrightarrow N2}(E) - \Psi_{N1\Leftrightarrow N2}(F)$ | $\beta_4$ | 0.003 | 22.2 % | 0.007 | 34.2 % | 0.029 | 29.6 % |
| chord | VN ⇔FPN-L | [0, 400] | dIES | $\Psi_{N1\Leftrightarrow N2}(E) - \Psi_{N1\Leftrightarrow N2}(F)$ | $\theta$ | 0.030 | 25.3 % | 0.005 | 36.0 % | 0.440 | 5.7 % |
| chord | VN ⇔FPN-L | [0, 400] | imp. hours | $\Psi_{N1\Leftrightarrow N2}(E) - \Psi_{N1\Leftrightarrow N2}(F)$ | $\gamma_1$ | 0.044 | 20.8 % | 0.958 | 4.1 % | 1.000 | 44.6 % |
| chord | VN ⇔FPN-R | [0, 400] | dIES | $\Psi_{N1\Leftrightarrow N2}(E) - \Psi_{N1\Leftrightarrow N2}(F)$ | $\theta$ | 0.038 | 30.3 % | 0.162 | 12.5 % | 0.317 | 8.8 % |
| chord | DAN-core ⇔FPN-L | [0, 400] | dIES | $\Psi_{N1\Leftrightarrow N2}(E) - \Psi_{N1\Leftrightarrow N2}(F)$ | $\beta_2$ | 0.010 | 22.4 % | 0.063 | 18.5 % | 0.365 | 6.0 % |
| chord | DAN-core ⇔FPN-L | [0, 400] | dIES | $\Psi_{N1\Leftrightarrow N2}(E) - \Psi_{N1\Leftrightarrow N2}(F)$ | $\beta_4$ | 1.23e-05 | 41.3 % | 0.383 | 4.8 % | 0.017 | 34.4 % |
| chord | DAN-core ⇔FPN-R | [0, 400] | imp. hours | $\Psi_{N1\Leftrightarrow N2}(E) - \Psi_{N1\Leftrightarrow N2}(F)$ | $\alpha_3$ | 0.001 | 26.7 % | 0.040 | 23.6 % | 1.000 | 40.7 % |
| response | VN ⇔DAN-core | [-400, 0] | dIES | $\Psi_{N1\Leftrightarrow N2}(E) - \Psi_{N1\Leftrightarrow N2}(F)$ | $\gamma_3$ | 0.043 | 22.6 % | 0.866 | 1.1 % | 0.034 | 28.6 % |
| response | VN ⇔DAN-core | [-400, 0] | cl. hours | $\Psi_{N1\Leftrightarrow N2}(E) - \Psi_{N1\Leftrightarrow N2}(F)$ | $\beta_4$ | 0.006 | 18.6 % | 0.369 | 7.4 % | 0.002 | 52.2 % |
| response | VN ⇔DAN-core | [-400, 0] | cl. hours | $\Psi_{N1\Leftrightarrow N2}(E) - \Psi_{N1\Leftrightarrow N2}(F)$ | $\gamma_1$ | 0.005 | 19.7 % | 0.009 | 33.1 % | 0.380 | 5.6 % |

Continued on next page

**Table 4 – continued from previous page**

| | | Time | | | Frequency | All musicians | | Improvisers | | Classical performers | |
|---|---|---|---|---|---|---|---|---|---|---|---|
| Locked to | Network | window | DV | IV | band | p | $R^2$ | p | $R^2$ | p | $R^2$ |
| response | VN ⇔DAN-core | [0, 400] | dIES | $\Psi_{N1\Leftrightarrow N2}(E) - \Psi_{N1\Leftrightarrow N2}(F)$ | $\alpha_1$ | 0.007 | 28.2 % | 0.069 | 18.3 % | 0.045 | 25.7 % |
| response | VN ⇔DAN-core | [0, 400] | dIES | $\Psi_{N1\Leftrightarrow N2}(E) - \Psi_{N1\Leftrightarrow N2}(F)$ | $\alpha_2$ | 0.032 | 29.8 % | 0.060 | 22.1 % | 0.022 | 32.2 % |
| response | VN ⇔DAN-core | [0, 400] | dIES | $\Psi_{N1\Leftrightarrow N2}(E) - \Psi_{N1\Leftrightarrow N2}(F)$ | $\alpha_3$ | 0.004 | 31.8 % | 0.020 | 28.5 % | 0.037 | 27.7 % |
| response | VN ⇔DAN-core | [0, 400] | dIES | $\Psi_{N1\Leftrightarrow N2}(E) - \Psi_{N1\Leftrightarrow N2}(F)$ | $\beta_1$ | 0.017 | 28.3 % | 0.207 | 9.4 % | 0.019 | 33.4 % |
| response | VN ⇔DAN-core | [0, 400] | imp. hours | $\Psi_{N1\Leftrightarrow N2}(E) - \Psi_{N1\Leftrightarrow N2}(F)$ | $\alpha_3$ | 0.004 | 21.8 % | 0.004 | 40.9 % | 1.000 | 48.9 % |
| response | VN ⇔DAN-core | [0, 400] | cl. hours | $\Psi_{N1\Leftrightarrow N2}(E) - \Psi_{N1\Leftrightarrow N2}(F)$ | $\beta_3$ | 0.005 | 19.6 % | 0.591 | 3.9 % | 0.046 | 25.7 % |
| response | VN ⇔FPN-L | [-400, 0] | dIES | $\Psi_{N1\Leftrightarrow N2}(E) - \Psi_{N1\Leftrightarrow N2}(F)$ | $\theta$ | 0.036 | 26.3 % | 0.075 | 19.4 % | 0.135 | 15.4 % |
| response | VN ⇔FPN-L | [-400, 0] | dIES | $\Psi_{N1\Leftrightarrow N2}(E) - \Psi_{N1\Leftrightarrow N2}(F)$ | $\beta_3$ | 0.030 | 30.2 % | 0.030 | 25.8 % | 0.117 | 20.2 % |
| response | VN ⇔FPN-L | [-400, 0] | dIES | $\Psi_{N1\Leftrightarrow N2}(E) - \Psi_{N1\Leftrightarrow N2}(F)$ | $\gamma_1$ | 0.007 | 23.1 % | 0.154 | 15.9 % | 0.421 | 4.8 % |
| response | VN ⇔FPN-L | [-400, 0] | dIES | $\Psi_{N1\Leftrightarrow N2}(E) - \Psi_{N1\Leftrightarrow N2}(F)$ | $\gamma_2$ | 0.018 | 21.3 % | 0.781 | 1.2 % | 0.009 | 40.1 % |
| response | VN ⇔FPN-L | [-400, 0] | dIES | $\Psi_{N1\Leftrightarrow N2}(E) - \Psi_{N1\Leftrightarrow N2}(F)$ | $\gamma_3$ | 0.043 | 22.5 % | 0.509 | 3.1 % | 0.003 | 49.4 % |
| response | VN ⇔FPN-L | [-400, 0] | cl. hours | $\Psi_{N1\Leftrightarrow N2}(E) - \Psi_{N1\Leftrightarrow N2}(F)$ | $\gamma_1$ | 0.005 | 19.7 % | 0.006 | 36.4 % | 0.505 | 3.2 % |
| response | VN ⇔FPN-L | [0, 400] | dIES | $\Psi_{N1\Leftrightarrow N2}(E) - \Psi_{N1\Leftrightarrow N2}(F)$ | $\delta$ | 0.021 | 31.8 % | 0.983 | 1.7 % | 6.82e-04 | 61.0 % |
| response | VN ⇔FPN-L | [0, 400] | dIES | $\Psi_{N1\Leftrightarrow N2}(E) - \Psi_{N1\Leftrightarrow N2}(F)$ | $\beta_2$ | 0.012 | 33.0 % | 0.018 | 29.3 % | 0.295 | 7.9 % |
| response | VN ⇔FPN-L | [0, 400] | imp. hours | $\Psi_{N1\Leftrightarrow N2}(E) - \Psi_{N1\Leftrightarrow N2}(F)$ | $\gamma_3$ | 0.014 | 18.9 % | 0.099 | 24.4 % | 1.000 | 44.4 % |
| response | VN ⇔FPN-R | [-400, 0] | dIES | $\Psi_{N1\Leftrightarrow N2}(E) - \Psi_{N1\Leftrightarrow N2}(F)$ | $\beta_2$ | 2.94e-04 | 39.1 % | 0.887 | 1.5 % | 0.011 | 38.3 % |
| response | VN ⇔FPN-R | [0, 400] | dIES | $\Psi_{N1\Leftrightarrow N2}(E) - \Psi_{N1\Leftrightarrow N2}(F)$ | $\beta_1$ | 0.033 | 25.0 % | 0.741 | 1.9 % | 0.008 | 40.2 % |
| response | VN ⇔FPN-R | [0, 400] | dIES | $\Psi_{N1\Leftrightarrow N2}(E) - \Psi_{N1\Leftrightarrow N2}(F)$ | $\beta_2$ | 0.032 | 23.4 % | 0.068 | 17.5 % | 0.163 | 13.6 % |
| response | VN ⇔FPN-R | [0, 400] | dIES | $\Psi_{N1\Leftrightarrow N2}(E) - \Psi_{N1\Leftrightarrow N2}(F)$ | $\beta_3$ | 0.003 | 27.7 % | 0.020 | 26.6 % | 0.114 | 17.2 % |
| response | VN ⇔FPN-R | [0, 400] | dIES | $\Psi_{N1\Leftrightarrow N2}(E) - \Psi_{N1\Leftrightarrow N2}(F)$ | $\gamma_1$ | 0.008 | 21.7 % | 0.154 | 11.4 % | 0.488 | 3.9 % |
| response | VN ⇔FPN-R | [0, 400] | imp. hours | $\Psi_{N1\Leftrightarrow N2}(E) - \Psi_{N1\Leftrightarrow N2}(F)$ | $\gamma_1$ | 0.015 | 18.7 % | 0.106 | 21.7 % | 1.000 | 41.3 % |
| response | VN ⇔FPN-R | [0, 400] | cl. hours | $\Psi_{N1\Leftrightarrow N2}(E) - \Psi_{N1\Leftrightarrow N2}(F)$ | $\beta_1$ | 0.001 | 24.5 % | 0.031 | 23.5 % | 0.023 | 32.9 % |
| response | DAN-core ⇔FPN-L | [-400, 0] | imp. hours | $\Psi_{N1\Leftrightarrow N2}(E) - \Psi_{N1\Leftrightarrow N2}(F)$ | $\beta_2$ | 0.004 | 24.1 % | 0.197 | 13.1 % | 1.000 | 40.9 % |
| response | DAN-core ⇔FPN-L | [0, 400] | dIES | $\Psi_{N1\Leftrightarrow N2}(E) - \Psi_{N1\Leftrightarrow N2}(F)$ | $\beta_2$ | 0.028 | 30.8 % | 0.010 | 34.2 % | 0.553 | 4.5 % |
| response | DAN-core ⇔FPN-L | [0, 400] | dIES | $\Psi_{N1\Leftrightarrow N2}(E) - \Psi_{N1\Leftrightarrow N2}(F)$ | $\beta_4$ | 7.64e-04 | 30.5 % | 0.253 | 8.0 % | 0.079 | 20.4 % |
| response | DAN-core ⇔FPN-L | [0, 400] | dIES | $\Psi_{N1\Leftrightarrow N2}(E) - \Psi_{N1\Leftrightarrow N2}(F)$ | $\gamma_3$ | 0.014 | 26.0 % | 0.321 | 6.0 % | 0.029 | 29.9 % |
| response | DAN-core ⇔FPN-R | [-400, 0] | dIES | $\Psi_{N1\Leftrightarrow N2}(E) - \Psi_{N1\Leftrightarrow N2}(F)$ | $\gamma_1$ | 0.030 | 24.3 % | 0.314 | 6.0 % | 4.35e-05 | 76.2 % |
| response | DAN-core ⇔FPN-R | [-400, 0] | dIES | $\Psi_{N1\Leftrightarrow N2}(E) - \Psi_{N1\Leftrightarrow N2}(F)$ | $\gamma_2$ | 0.022 | 28.8 % | 0.990 | 2.6 % | 0.007 | 44.1 % |

Continued on next page

**Table 4 – continued from previous page**

| Locked to | Network | Time window | DV | IV | Frequency band | All musicians p | All musicians $R^2$ | Improvisers p | Improvisers $R^2$ | Classical performers p | Classical performers $R^2$ |
|---|---|---|---|---|---|---|---|---|---|---|---|
| response | DAN-core ⇔FPN-R | [0, 400] | dIES | $\Psi_{N1\Leftrightarrow N2}(E) - \Psi_{N1\Leftrightarrow N2}(F)$ | $\theta$ | 0.018 | 32.6 % | 0.003 | 40.6 % | 0.718 | 1.9 % |
| response | DAN-core ⇔FPN-R | [0, 400] | dIES | $\Psi_{N1\Leftrightarrow N2}(E) - \Psi_{N1\Leftrightarrow N2}(F)$ | $\alpha_3$ | 0.042 | 28.6 % | 0.078 | 20.0 % | 0.198 | 12.0 % |
| response | DAN-core ⇔FPN-R | [0, 400] | dIES | $\Psi_{N1\Leftrightarrow N2}(E) - \Psi_{N1\Leftrightarrow N2}(F)$ | $\beta_1$ | 0.048 | 20.4 % | 0.195 | 11.6 % | 0.027 | 30.4 % |
| response | DAN-core ⇔FPN-R | [0, 400] | dIES | $\Psi_{N1\Leftrightarrow N2}(E) - \Psi_{N1\Leftrightarrow N2}(F)$ | $\beta_2$ | 0.013 | 29.7 % | 0.012 | 31.6 % | 0.183 | 12.6 % |
| response | DAN-core ⇔FPN-R | [0, 400] | dIES | $\Psi_{N1\Leftrightarrow N2}(E) - \Psi_{N1\Leftrightarrow N2}(F)$ | $\beta_3$ | 0.013 | 28.3 % | 0.205 | 9.0 % | 0.779 | 1.2 % |
| response | DAN-core ⇔FPN-R | [0, 400] | dIES | $\Psi_{N1\Leftrightarrow N2}(E) - \Psi_{N1\Leftrightarrow N2}(F)$ | $\gamma_3$ | 0.002 | 34.7 % | 0.021 | 27.0 % | 0.227 | 10.4 % |
| response | DAN-core ⇔FPN-R | [0, 400] | imp. hours | $\Psi_{N1\Leftrightarrow N2}(E) - \Psi_{N1\Leftrightarrow N2}(F)$ | $\beta_3$ | 1.50e-06 | 51.3 % | 0.339 | 8.6 % | 1.000 | 41.7 % |
| chord | DMN ⇔SN-core | [0, 400] | dIES | $\Psi_{N1\Leftrightarrow N2}(E) - \Psi_{N1\Leftrightarrow N2}(F)$ | $\beta_2$ | 0.014 | 26.7 % | 0.002 | 43.8 % | 0.857 | 0.7 % |
| chord | DMN ⇔CON | [0, 400] | dIES | $\Psi_{N1\Leftrightarrow N2}(E) - \Psi_{N1\Leftrightarrow N2}(F)$ | $\beta_2$ | 0.025 | 23.7 % | 3.50e-04 | 51.8 % | 0.368 | 7.1 % |
| chord | DMN ⇔CON | [0, 400] | dIES | $\Psi_{N1\Leftrightarrow N2}(E) - \Psi_{N1\Leftrightarrow N2}(F)$ | $\beta_4$ | 0.001 | 28.9 % | 0.812 | 1.6 % | 0.053 | 25.4 % |
| response | DMN ⇔SN-core | [-400, 0] | dIES | $\Psi_{N1\Leftrightarrow N2}(E) - \Psi_{N1\Leftrightarrow N2}(F)$ | $\delta$ | 0.012 | 36.6 % | 0.038 | 24.2 % | 0.963 | 0.7 % |
| response | DMN ⇔SN-core | [-400, 0] | dIES | $\Psi_{N1\Leftrightarrow N2}(E) - \Psi_{N1\Leftrightarrow N2}(F)$ | $\gamma_1$ | 0.005 | 28.2 % | 0.023 | 26.8 % | 0.070 | 21.6 % |
| response | DMN ⇔SN-core | [0, 400] | dIES | $\Psi_{N1\Leftrightarrow N2}(E) - \Psi_{N1\Leftrightarrow N2}(F)$ | $\beta_1$ | 0.042 | 23.6 % | 0.460 | 4.0 % | 0.019 | 33.4 % |
| response | DMN ⇔SN-core | [0, 400] | dIES | $\Psi_{N1\Leftrightarrow N2}(E) - \Psi_{N1\Leftrightarrow N2}(F)$ | $\beta_3$ | 0.008 | 30.3 % | 0.093 | 15.1 % | 0.207 | 12.4 % |
| response | DMN ⇔CON | [-400, 0] | dIES | $\Psi_{N1\Leftrightarrow N2}(E) - \Psi_{N1\Leftrightarrow N2}(F)$ | $\beta_1$ | 0.003 | 28.5 % | 0.466 | 4.6 % | 0.035 | 28.2 % |
| response | DMN ⇔CON | [0, 400] | dIES | $\Psi_{N1\Leftrightarrow N2}(E) - \Psi_{N1\Leftrightarrow N2}(F)$ | $\beta_2$ | 0.019 | 27.7 % | 0.010 | 32.2 % | 0.515 | 3.6 % |
| response | DMN ⇔CON | [0, 400] | dIES | $\Psi_{N1\Leftrightarrow N2}(E) - \Psi_{N1\Leftrightarrow N2}(F)$ | $\gamma_3$ | 0.017 | 28.5 % | 0.040 | 22.5 % | 0.417 | 5.0 % |
